# Fine-mapping of 150 breast cancer risk regions identifies 178 high confidence target genes

**DOI:** 10.1101/521054

**Authors:** Laura Fachal, Hugues Aschard, Jonathan Beesley, Daniel R. Barnes, Jamie Allen, Siddhartha Kar, Karen A. Pooley, Joe Dennis, Kyriaki Michailidou, Constance Turman, Penny Soucy, Audrey Lemaçon, Michael Lush, Jonathan P. Tyrer, Maya Ghoussaini, Mahdi Moradi Marjaneh, Xia Jiang, Simona Agata, Kristiina Aittomäki, M. Rosario Alonso, Irene L. Andrulis, Hoda Anton-Culver, Natalia N. Antonenkova, Adalgeir Arason, Volker Arndt, Kristan J. Aronson, Banu K. Arun, Bernd Auber, Paul L. Auer, Jacopo Azzollini, Judith Balmaña, Rosa B. Barkardottir, Daniel Barrowdale, Alicia Beeghly-Fadiel, Javier Benitez, Marina Bermisheva, Katarzyna Bialkowska, Amie M. Blanco, Carl Blomqvist, William Blot, Natalia V. Bogdanova, Stig E. Bojesen, Manjeet K. Bolla, Bernardo Bonanni, Ake Borg, Kristin Bosse, Hiltrud Brauch, Hermann Brenner, Ignacio Briceno, Ian W. Brock, Angela Brooks-Wilson, Thomas Brüning, Barbara Burwinkel, Saundra S. Buys, Qiuyin Cai, Trinidad Caldés, Maria A. Caligo, Nicola J. Camp, Ian Campbell, Federico Canzian, Jason S. Carroll, Brian D. Carter, Jose E. Castelao, Jocelyne Chiquette, Hans Christiansen, Wendy K. Chung, Kathleen B.M. Claes, Christine L. Clarke, GEMO Study Collaborators, EMBRACE Collaborators, J. Margriet Collée, Sten Cornelissen, Fergus J. Couch, Angela Cox, Simon S. Cross, Cezary Cybulski, Kamila Czene, Mary B. Daly, Miguel de la Hoya, Peter Devilee, Orland Diez, Yuan Chun Ding, Gillian S. Dite, Susan M. Domchek, Thilo Dörk, Isabel dos-Santos-Silva, Arnaud Droit, Stéphane Dubois, Martine Dumont, Mercedes Duran, Lorraine Durcan, Miriam Dwek, Diana M. Eccles, Christoph Engel, Mikael Eriksson, D. Gareth Evans, Peter A. Fasching, Olivia Fletcher, Giuseppe Floris, Henrik Flyger, Lenka Foretova, William D. Foulkes, Eitan Friedman, Lin Fritschi, Debra Frost, Marike Gabrielson, Manuela Gago-Dominguez, Gaetana Gambino, Patricia A. Ganz, Susan M. Gapstur, Judy Garber, José A. García-Sáenz, Mia M. Gaudet, Vassilios Georgoulias, Graham G. Giles, Gord Glendon, Andrew K. Godwin, Mark S. Goldberg, David E. Goldgar, Anna González-Neira, Mark H. Greene, Mervi Grip, Jacek Gronwald, Anne Grundy, Pascal Guénel, Eric Hahnen, Christopher A. Haiman, Niclas Håkansson, Per Hall, Ute Hamann, Patricia A. Harrington, Jaana M. Hartikainen, Mikael Hartman, Wei He, Catherine S. Healey, Bernadette A.M. Heemskerk-Gerritsen, Jane Heyworth, Peter Hillemanns, Frans B.L. Hogervorst, Antoinette Hollestelle, Maartje J. Hooning, John L. Hopper, Anthony Howell, Guanmengqian Huang, Peter J. Hulick, Evgeny N. Imyanitov, ABCTB Investigators, KConFab Investigators, HEBON Investigators, Claudine Isaacs, Motoki Iwasaki, Agnes Jager, Milena Jakimovska, Anna Jakubowska, Paul James, Ramunas Janavicius, Rachel C. Jankowitz, Esther M. John, Nichola Johnson, Michael E. Jones, Arja Jukkola-Vuorinen, Audrey Jung, Rudolf Kaaks, Daehee Kang, Beth Y. Karlan, Renske Keeman, Michael J. Kerin, Elza Khusnutdinova, Johanna I. Kiiski, Judy Kirk, Cari M. Kitahara, Yon-Dschun Ko, Irene Konstantopoulou, Veli-Matti Kosma, Stella Koutros, Katerina Kubelka-Sabit, Ava Kwong, Kyriacos Kyriacou, Yael Laitman, Diether Lambrechts, Eunjung Lee, Goska Leslie, Jenny Lester, Fabienne Lesueur, Annika Lindblom, Wing-Yee Lo, Jirong Long, Artitaya Lophatananon, Jennifer T. Loud, Jan Lubinski, Robert J. MacInnis, Tom Maishman, Enes Makalic, Arto Mannermaa, Mehdi Manoochehri, Siranoush Manoukian, Sara Margolin, Maria Elena Martinez, Keitaro Matsuo, Tabea Maurer, Dimitrios Mavroudis, Rebecca Mayes, Lesley McGuffog, Catriona McLean, Noura Mebirouk, Alfons Meindl, Pooja Middha, Nicola Miller, Austin Miller, Marco Montagna, Fernando Moreno, Anna Marie Mulligan, Victor M. Muñoz-Garzon, Taru A. Muranen, Steven A. Narod, Rami Nassir, Katherine L. Nathanson, Susan L. Neuhausen, Heli Nevanlinna, Patrick Neven, Finn C. Nielsen, Liene Nikitina-Zake, Aaron Norman, Kenneth Offit, Edith Olah, Olufunmilayo I. Olopade, Håkan Olsson, Nick Orr, Ana Osorio, V. Shane Pankratz, Janos Papp, Sue K. Park, Tjoung-Won Park-Simon, Michael T. Parsons, James Paul, Inge Sokilde Pedersen, Bernard Peissel, Beth Peshkin, Paolo Peterlongo, Julian Peto, Dijana Plaseska-Karanfilska, Karolina Prajzendanz, Ross Prentice, Nadege Presneau, Darya Prokofyeva, Miquel Angel Pujana, Katri Pylkäs, Paolo Radice, Susan J. Ramus, Johanna Rantala, Rohini Rau-Murthy, Gad Rennert, Harvey A. Risch, Mark Robson, Atocha Romero, Caroline Maria Rossing, Emmanouil Saloustros, Estela Sánchez-Herrero, Dale P. Sandler, Marta Santamariña, Christobel Saunders, Elinor J. Sawyer, Maren T. Scheuner, Daniel F. Schmidt, Rita K. Schmutzler, Andreas Schneeweiss, Minouk J. Schoemaker, Ben Schöttker, Peter Schürmann, Christopher Scott, Rodney J. Scott, Leigha Senter, Caroline MD Seynaeve, Mitul Shah, Priyanka Sharma, Chen-Yang Shen, Xiao-Ou Shu, Christian F. Singer, Thomas P. Slavin, Snezhana Smichkoska, Melissa C. Southey, John J. Spinelli, Amanda B. Spurdle, Jennifer Stone, Dominique Stoppa-Lyonnet, Christian Sutter, Anthony J. Swerdlow, Rulla M. Tamimi, Yen Yen Tan, William J. Tapper, Jack A. Taylor, Manuel R. Teixeira, Maria Tengström, Soo H. Teo, Mary Beth Terry, Alex Teulé, Mads Thomassen, Darcy L. Thull, Maria Grazia Tibiletti, Marc Tischkowitz, Amanda E. Toland, Rob A.E.M. Tollenaar, Ian Tomlinson, Diana Torres, Gabriela Torres-Mejía, Melissa A. Troester, Nadine Tung, Maria Tzardi, Hans-Ulrich Ulmer, Celine M. Vachon, Christi J. van Asperen, Lizet E. van der Kolk, Elizabeth J. van Rensburg, Ana Vega, Alessandra Viel, Joseph Vijai, Maatje J. Vogel, Qin Wang, Barbara Wappenschmidt, Clarice R. Weinberg, Jeffrey N. Weitzel, Camilla Wendt, Hans Wildiers, Robert Winqvist, Alicja Wolk, Anna H. Wu, Drakoulis Yannoukakos, Yan Zhang, Wei Zheng, Paul D.P. Pharoah, Jenny Chang-Claude, Montserrat García-Closas, Marjanka K. Schmidt, Roger L. Milne, Vessela N. Kristensen, Juliet D. French, Stacey L. Edwards, Antonis C. Antoniou, Georgia Chenevix-Trench, Jacques Simard, Douglas F. Easton, Peter Kraft, Alison M. Dunning

## Abstract

Genome-wide association studies have identified breast cancer risk variants in over 150 genomic regions, but the mechanisms underlying risk remain largely unknown. These regions were explored by combining association analysis with *in silico* genomic feature annotations. We defined 205 independent risk-associated signals with the set of credible causal variants (CCVs) in each one. In parallel, we used a Bayesian approach (PAINTOR) that combines genetic association, linkage disequilibrium, and enriched genomic features to determine variants with high posterior probabilities (HPPs) of being causal. Potentially causal variants were significantly over-represented in active gene regulatory regions and transcription factor binding sites. We applied our INQUSIT pipeline for prioritizing genes as targets of potentially causal variants, using gene expression (eQTL), chromatin interaction and functional annotations. Known cancer drivers, transcription factors and genes in the developmental, apoptosis, immune system and DNA integrity checkpoint gene ontology pathways, were over-represented among the 178 highest confidence target genes.

## INTRODUCTION

Genome-wide association studies (GWAS) have identified genetic variants associated with breast cancer risk in more than 150 genomic regions ^1,2^. However, the variants and genes driving these associations are mostly unknown, with fewer than 20 regions studied in detail ^3-20^. Here, we aimed to fine-map all known breast cancer susceptibility regions using dense genotype data on > 217K subjects participating in the Breast Cancer Association Consortium (BCAC) and the Consortium of Investigators of Modifiers of *BRCA1/2* (CIMBA). All samples were genotyped using the OncoArray^TM 1,2,21^ or the iCOGS chip ^22,23^. Stepwise multinomial logistic regression was used to identify independent association signals in each region and define credible causal variants (CCVs) within each signal. We found genomic features significantly overlapping the CCVs and then used a Bayesian approach, integrating informative genomic features and genetic associations, to refine the set of likely causal variants and calculate their posterior probabilities. Finally, we integrated genetic and *in silico* epigenetic, expression and chromatin conformation data to infer the likely target genes of each signal.

## RESULTS

### Most breast cancer genomic regions contain multiple independent risk-associated signals

We included 109,900 breast cancer cases and 88,937 controls, all of European ancestry, from 75 studies in the BCAC. Genotypes (directly observed or imputed) were available for 639,118 single nucleotide polymorphisms (SNPs), deletion/insertions, and copy number variants (CNVs) with minor allele frequency (MAF) ≥ 0.1% within 152, previously defined, risk-associated regions (**Supplementary Table 1; Figure 1**). Multivariate logistic regression confirmed associations for 150/152 regions at a p-value < 10^−4^ significance threshold (**Supplementary Table 2A**). To determine the number of independent risk signals within each region we applied stepwise multinomial logistic regression, deriving the association of each variant, conditional on the more significant ones, in order of statistical significance. Finally, we defined CCVs in each signal as variants with conditional p-values within two orders of magnitude of the index variant ^24^. We classified the evidence for each independent signal, and it’s CCVs, as either *strong* (conditional p-values <10^−6^) or *moderate* (10^−6^ < conditional p-values <10^−4^).

**Figure 1.**
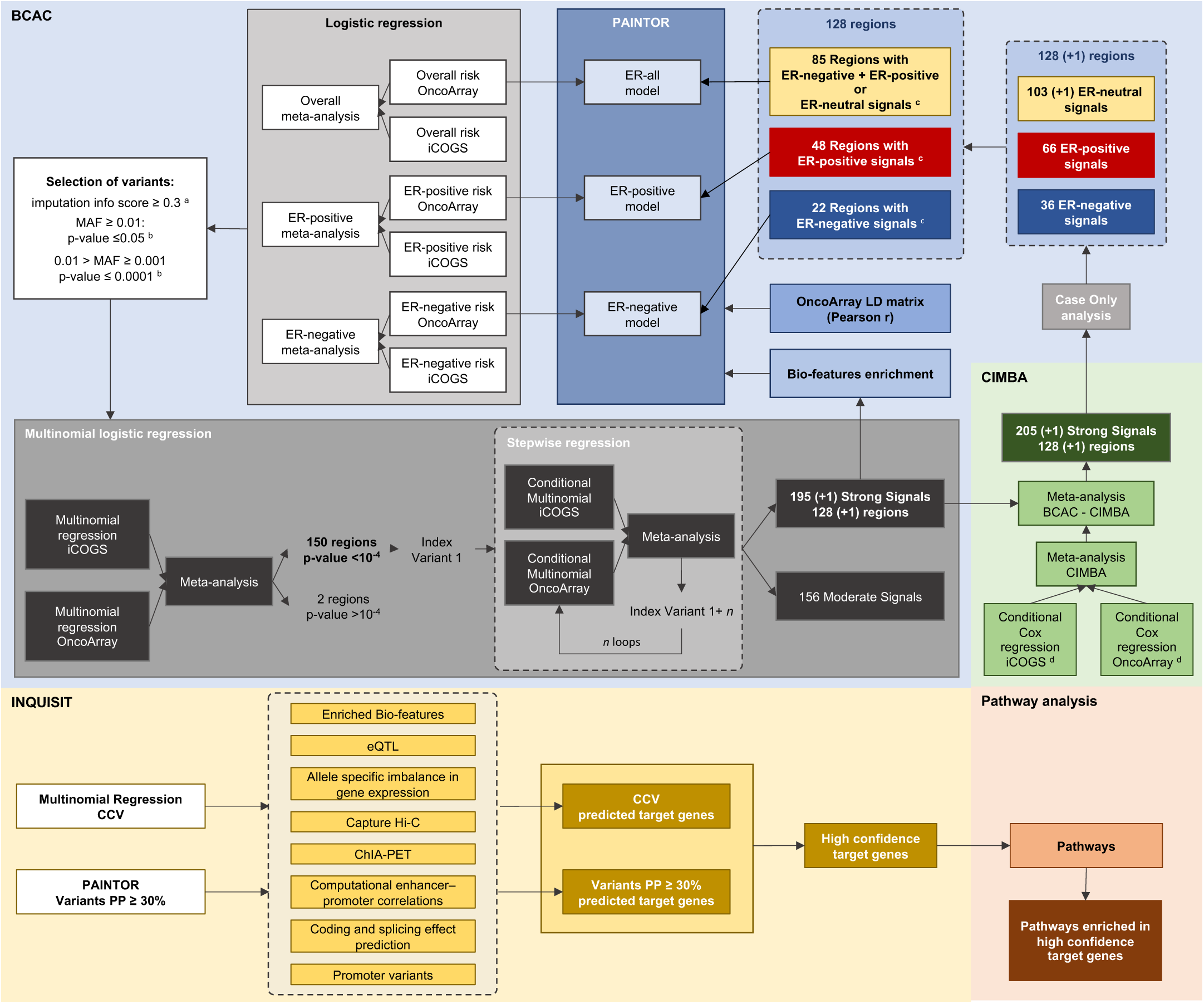
Flowchart summarizing the study design. ^a^ In iCOGS and in OncoArray; ^b^ for at least one phenotype; ^c^ regions containing signals associated with more than one phenotype were evaluated with more than one PAINTOR model; ^d^ conditional on the index variants from BCAC strong signals.

From the 150 genomic regions we identified 352 independent risk signals containing 13,367 CCVs, 7,394 of these were within the 196 strong-evidence signals across 129 regions (**Figures 2A-B**). The number of signals per region ranged from 1 to 9, with 79 (53%) containing multiple signals. We noted a wide range of CCVs per signal, but in 42 signals there was only a single CCV and this was assumed to be causal (**Figures 2C-D, Table 1**).

**Figure 2.**
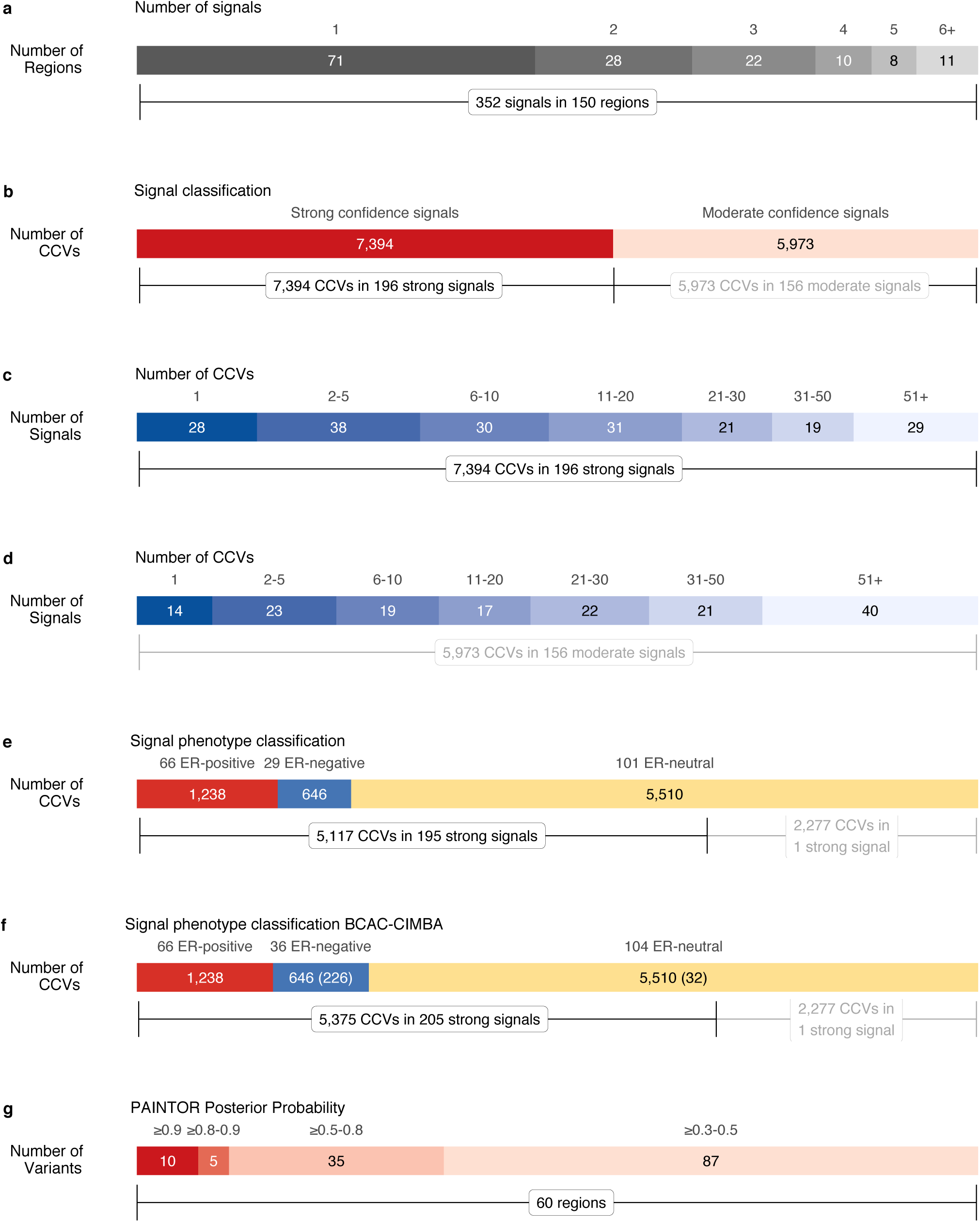
Determining independent risk signals and credible candidate variants (CCVs). (a) Number of independent signals per region identified through multinomial stepwise logistic regression. (b) Signal classification according to their confidence into strong and moderate confidence signals. (c) Number of CCVs per signal at strong confidence signals identified through multinomial stepwise logistic regression. (d) Number of CCVs per signal at moderate confidence signals identified through multinomial stepwise regression. (e) Subtype classification of strong signals into ER-positive, ER-negative and signals equally associated with both phenotypes (ER-neutral) from BCAC analysis. (f) Subtype classification from the meta-analysis of BCAC and CIMBA. Between brackets, number of CCVs from the meta-analysis of BCAC and CIMBA. (g) Number of variants at different posterior probability thresholds. 15 variants reach a PP ≥ 80% by at least one of the three models (ER-all, ER-positive, ER-negative).

**Table 1.** Signals with single CCVs and variants with PP > 80%.

| Fine-mapping region<sup>a</sup> | Variant <sup>b</sup> | Ref/Alt <sup>c</sup> | EAF<sup>d</sup> | PP<sup>e</sup> | Model<sup>f</sup> | Signal<sup>g</sup> | N CCV<sup>h</sup> | ER-negative |  | ER-positive |  | P-value<sup>i</sup> | FP<sup>j</sup> | Predicted target gene(s)<sup>k</sup> | Confidence<sup>l</sup> |
| --- | --- | --- | --- | --- | --- | --- | --- | --- | --- | --- | --- | --- | --- | --- | --- |
|  |  |  |  |  |  |  |  | OR<sup>i</sup> | (95%CI) | OR<sup>i</sup> | (95%CI) |  |  |  |  |
| chr1:120723447-121780613 | rs11249433 | A/G | 0.42 | 0.57 | ERALL | Signal 1 | 1 | 1.02 | (0.99-1.04) | 1.13 | (1.11-1.15) | 8.11x10<sup>-60</sup> | na | na |  |
| chr1:200937832-201937832 | rs35383942 | C/T | 0.06 | 0.96 | ERALL | Signal 1 | 2 | 1.10 | (1.05-1.16) | 1.09 | (1.06-1.13) | 1.14x10<sup>-7</sup> | D | *TNNI1* | Level 1 |
| chr2:201681247-202681247 | rs3769821 | C/T | 0.66 | 0.40 | ERALL | Signal 1 | 1 | 0.94 | (0.92-0.97) | 0.95 | (0.93-0.96) | 1.46x10<sup>-12</sup> | D | *ALS2CR11*, *ALS2CR12*, *CASP8<sup>m</sup>*, *MPP4*, *NIF3L1*, *STRADB*, *TMEM237* | Level 2 |
| chr2:217405832-218796508 | rs4442975 | G/T | 0.48 | 0.84 | ERALL | Signal 1 | 1 | 0.94 | (0.92-0.97) | 0.86 | (0.85-0.87) | 2.50x10<sup>-90</sup> | D | *IGFBP5<sup>m</sup>* | Level 1 |
| chr4:105569013-106856761 | esv3601665 | -/Alu | 0.07 | 0.95 | ERPOS |  |  | 1.01 | (0.95-1.08) | 1.10 | (1.06-1.14) | 3.27x10<sup>-6</sup> | D | *ARHGEF38*, *AC004066.3* | Level 1 |
| chr5:779790-1797488 | rs10069690 | C/T | 0.27 | 0.58 | ERNEG | Signal 1 | 1 | 1.18 | (1.15-1.21) | 1.03 | (1.01-1.05) | 1.20x10<sup>-34</sup> | D | *CLPTM1L*, *TERT<sup>m</sup>* | Level 1 |
| chr5:44013304-45206498 | rs10941679 | A/G | 0.26 | 0.00 | ERPOS | Signal 1 | 1 | 1.04 | (1.02-1.07) | 1.17 | (1.15-1.19) | 1.50x10<sup>-77</sup> | D | *MRPS30* | Level 2 |
|  | rs5867671 | A/- | 0.77 | 0.01 | ERPOS | Signal 2 | 1 | 0.91 | (0.89-0.94) | 0.99 | (0.97-1.01) | 2.25x10<sup>-9</sup> | na | na |  |
| chr5:44013304-45206498 | rs190443933 | T/C | 0.01 | 0.00 | ERALL | Signal 4 | 1 | 1.30 | (1.14-1.48) | 1.26 | (1.16-1.37) | 2.32x10<sup>-8</sup> | na | na |  |
| chr5:55531884-56587883 | rs984113 | G/C | 0.61 | 0.81 | ERPOS | Signal 2 | 1 | 0.96 | (0.93-0.98) | 0.96 | (0.94-0.97) | 3.51x10<sup>-8</sup> | D | *MAP3K1<sup>m</sup>* | Level 1 |
|  | rs889310 | C/T | 0.56 | 0.84 | ERPOS | (Signal 6) | 15 | 1.03 | (1.00-1.05) | 1.05 | (1.03-1.06) | 1.75x10<sup>-7</sup> | D | *MAP3K1<sup>m</sup>* | Level 1 |
| chr6:15899557-16899557 | rs3819405 | C/T | 0.32 | 0.96 | ERALL | Signal 1 | 1 | 0.97 | (0.95-1.00) | 0.95 | (0.94-0.97) | 1.14x10<sup>-7</sup> | D | *ATXN1*, *RP1-151F17.1*, *RP1-151F17.2* | Level 2 |
| chr6:151418856-152937016 | rs12173562 | C/T | 0.08 | 0.10 | ERNEG | Signal 1 | 1 | 1.30 | (1.25-1.36) | 1.14 | (1.11-1.18) | 3.98x10<sup>-40</sup> | D | *ESR1<sup>m</sup>* | Level 1 |
|  | rs34133739 | -/C | 0.53 | 0.25 | ERALL | Signal 2 | 1 | 1.11 | (1.09-1.14) | 1.05 | (1.04-1.07) | 2.36x10<sup>-22</sup> | D | *ESR1<sup>m</sup>*, *SYNE1* | Level 2 |
|  | rs851984 | G/A | 0.40 | 0.73 | ERALL | Signal 3 | 1 | 1.07 | (1.04-1.09) | 1.05 | (1.04-1.07) | 3.69x10<sup>-13</sup> | D | *ESR1<sup>m</sup>* | Level 1 |
| chr7:130167121-131167121 | rs68056147 | G/A | 0.30 | 0.84 | ERALL |  |  | 1.04 | (1.01-1.07) | 1.05 | (1.03-1.06) | 3.07x10<sup>-7</sup> | D | *MKLN1* | Level 2 |
| chr8:127424659-130041931 | rs35961416 | -/A | 0.41 | 0.68 | ERALL | Signal 3 | 1 | 0.97 | (0.94-0.99) | 0.95 | (0.93-0.96) | 9.97x10<sup>-11</sup> | D | *CASC11*, *MYC<sup>m</sup>* | Level 2 |
| chr9:21247803-22624477 | rs539723051 | AAAA/- | 0.33 | 0.43 | ERALL | Signal 1 | 1 | 1.08 | (1.05-1.11) | 1.06 | (1.04-1.08) | 1.81x10<sup>-15</sup> | na | na |  |
| chr9:109803808-111395353 | rs10816625 | A/G | 0.07 | 0.95 | ERPOS | Signal 3 | 1 | 1.06 | (1.01-1.11) | 1.13 | (1.10-1.16) | 3.62x10<sup>-15</sup> | D | *KLF4<sup>m</sup>* | Level 2 |
|  | rs13294895 | C/T | 0.18 | 0.93 | ERPOS | Signal 4 | 1 | 1.01 | (0.98-1.05) | 1.09 | (1.07-1.11) | 4.00x10<sup>-17</sup> | D | *KLF4<sup>m</sup>* | Level 1 |
| chr9:109803808-111395353 | rs60037937 | AA/- | 0.22 | 0.68 | ERPOS | Signal 2 | 1 | 1.02 | (0.99-1.06) | 1.11 | (1.09-1.13) | 3.17x10<sup>-26</sup> | D | *KLF4<sup>m</sup>, RAD23B* | Level 2 |
| chr10:63758684-65063702 | rs10995201 | A/G | 0.15 | 0.31 | ERALL | Signal 1 | 1 | 0.91 | (0.88-0.94) | 0.87 | (0.85-0.89) | 1.40x10<sup>-37</sup> | na | na |  |
| chr10:122593901-123849324 | rs45631563 | A/T | 0.04 | 0.93 | ERPOS | Signal 3 | 1 | 0.97 | (0.92-1.03) | 0.76 | (0.73-0.79) | 4.84x10<sup>-44</sup> | C | *FGFR2<sup>m</sup>* | Level 1 |
| chr10:122593901-123849324 | rs35054928 | C/- | 0.56 | 0.60 | ERALL | Signal 1 | 1 | 0.96 | (0.94-0.98) | 0.74 | (0.73-0.76) | 6.55x10<sup>-342</sup> | D | *FGFR2<sup>m</sup>* | Level 1 |
|  | rs45631563 | A/T | 0.04 | 0.93 | ERPOS | Signal 3 | 1 | 0.97 | (0.92-1.03) | 0.76 | (0.73-0.79) | 4.84x10<sup>-44</sup> | C | *FGFR2<sup>m</sup>* | Level 1 |
|  | rs7899765 | T/C | 0.06 | 0.02 | ERALL | Signal 5 | 1 | 1.01 | (0.97-1.06) | 0.87 | (0.84-0.90) | 2.21x10<sup>-18</sup> | D | *FGFR2<sup>m</sup>* | Level 1 |
| chr11:68831418-69879161 | rs78540526 | C/T | 0.09 | 0.91 | ERPOS | Signal 1 | 1 | 1.01 | (0.97-1.06) | 1.40 | (1.36-1.44) | 2.77x10<sup>-145</sup> | D | *CCND1<sup>m</sup>, MYEOV* | Level 1 |
| chr12:27639846-29034415 | rs7297051 | C/T | 0.23 | 0.23 | ERALL | Signal 1 | 1 | 0.87 | (0.85-0.90) | 0.89 | (0.88-0.91) | 3.12x10<sup>-43</sup> | D | *CCDC91<sup>m</sup>, PTHLH<sup>m</sup>, RP11-967K21.1* | Level 2 |
| chr12:115336522-116336522 | rs35422 | G/A | 0.57 | 0.58 | ERPOS | Signal 2 | 1 | 0.98 | (0.96-1.01) | 1.05 | (1.03-1.07) | 4.85x10<sup>-10</sup> | D | *TBX3* | Level 1 |
| chr14:91341069-92368623 | rs7153397 | C/T | 0.70 | 0.81 | ERPOS | Signal 1 | 3 | 1.01 | (0.99-1.04) | 1.06 | (1.04-1.08) | 3.25x10<sup>-11</sup> | D,C | *CCDC88C, CTD-2547L24.4, GPR68, RP11-73M18.7, RP11-895M11.3* | Level 2 |
| chr16:52038825-53038825 | rs4784227 | C/T | 0.27 | 0.95 | ERPOS | Signal 1 | 1 | 1.15 | (1.12-1.18) | 1.26 | (1.24-1.28) | 4.63x10<sup>-160</sup> | D | *TOX3<sup>m</sup>* | Level 1 |
| chr18:23832476-25075396 | rs180952292 | T/C | 0.01 | 0.01 | ERNEG | Signal 4 | 1 | 1.24 | (1.12-1.37) | 0.98 | (0.92-1.05) | 2.07x10<sup>-5</sup> | na | na |  |
| chr18:41899590-42899590 | rs9952980 | T/C | 0.34 | 0.95 | ERALL | Signal 2 | 3 | 0.97 | (0.94-0.99) | 0.95 | (0.93-0.96) | 7.43x10<sup>-12</sup> | D | *SETBP1* | Level 2 |
| chr20:5448227-6448227 | rs16991615 | G/A | 0.07 | 0.97 | ERALL | Signal 1 | 1 | 1.09 | (1.04-1.15) | 1.07 | (1.04-1.11) | 7.89x10<sup>-7</sup> | D, C | *GPCPD1, MCM8* | Level 2 |
| chr22:45783297-46783297 | rs184070480 | C/T | 0.01 | 0.00 | ERALL | Signal 2 | 1 | 1.40 | (1.20-1.64) | 1.01 | (0.91-1.12) | 5.02x10<sup>-5</sup> | D | *ATXN10, WNT7B* | Level 2 |

<sup>a</sup> GRCh37/hg19, bp
<sup>b</sup> Current reference ID
<sup>c</sup> Reference (Ref) versus Alternative (Alt) Allele
<sup>d</sup> Effect allele (Alt allele) frequency in OncoArray
<sup>e</sup> PP: Posterior probability. Largest posterior probability in all evaluated models
<sup>f</sup> Model where the variant reaches the largest posterior probability
<sup>g</sup> Signal where the variant is included. Between brackets moderate confidence signals.
<sup>h</sup> Number of CCVs in the signal
<sup>i</sup> Multinomial regression summary statistics. Single variant analysis
<sup>j</sup> D: Distal regulation, P: proximal regulation, C: coding; na: prediction non available
<sup>k</sup> Predicted target genes with the largest confidence level for each variant. Between brackets, largest confidence level. na: prediction non available
<sup>l</sup> INQUISIT level of confidence
<sup>m</sup> Target genes with functional follow up

The majority of breast tumors express the estrogen receptor (ER-positive), but ∼20% do not (ER-negative); these two tumor types have distinct biological and clinical characteristics ^25^. Using a case-only analysis for the 196 strong-evidence signals, we found 66 signals (34%; containing 1,238 CCVs) where the lead variant conferred a greater relative-risk of developing ER-positive tumors (false discovery rate, FDR 5%), and 29 (15%; 646 CCVs) where the lead variant conferred a greater risk of ER-negative cancer tumors (FDR 5%) (**Supplementary Table 2B, Figure 2E**). The remaining 101 signals (51%, 5,510 CCVs) showed no difference by ER status (referred to as ER-neutral).

Patients with *BRCA1* mutations are more likely to develop ER-negative tumors ^26^. Hence, to increase our power to identify ER-negative signals, we performed a fixed-effects meta-analysis, combining association results from *BRCA1* mutation carriers in CIMBA with the BCAC ER-negative association results. This meta-analysis identified ten additional signals, seven ER-negative and three ER-neutral, making 206 strong-evidence signals (17% ER-negative) containing 7,652 CCVs in total (**Figure 2F**). More than one quarter of the CCVs (2,277) were accounted for by one signal, resulting from strong linkage disequilibrium with a copy number variant. The remaining analyses focused on the other 205 strong signals across 128 regions (**Supplementary Table 2C**).

### CCVs are over-represented in active gene-regulatory regions and transcription factor binding sites (TFBSs)

We constructed a database of mapped genomic-features in seven primary cells derived from normal breast and 19 breast cell lines using publicly available data, resulting in 811 annotation tracks in total. These ranged from general features, such as whether a variant was in an exon or in open chromatin, to more specific features, such a cell-specific TF binding or histone mark (determined through ChIP-Seq experiments) in breast-derived cells or cell lines. Using logistic regression, we examined the overlap of these genomic-features with the positions of 5,117 CCVs in the 195 strong-evidence BCAC signals versus the positions of 622,903 variants excluded as credible candidates in the same regions (**Supplementary Figure 1A, Supplementary Table 3**). We found significant enrichment of CCVs (FDR 5%) in the following genomic-features:

i. Open chromatin (determined by DNase-seq and FAIRE-seq) in ER-positive breast cancer cell-lines and normal breast (**Figure 3A**). Conversely, we found depletion of CCVs within heterochromatin (determined by the H3K9me3 mark in normal breast, and by chromatin-state in ER-positive cells ^27^).
ii. Actively transcribed genes in normal breast and ER-positive cell lines (defined by H3K36me3 or H3K79me2 histone marks, **Figure 3A**). Enrichment was larger for ER-neutral CCVs than for those affecting either ER-positive or ER-negative tumors.
iii. Gene regulatory regions. CCVs overlapped distal gene regulatory elements in ER-positive breast cancer cells lines (defined by H3K4me1 or H3K27ac marks, **Figure 3B**). This was confirmed using the ENCODE definition of active enhancers in MCF-7 cells (enhancer-like regions defined by combining DNase and H3K27ac marks), as well as the definition of ^28^ and ^27^ (Supplementary Table 3). Under these more stringent definitions, enrichment among ER-positive CCVs was significantly larger than ER-negative or ER-neutral CCVs. Data from ^27^, showed that 73% of active enhancer regions overlapped by ER-positive CCVs in ER-positive cells (MCF-7), are inactive in the normal HMEC breast cell line; thus, these enhancers appear to be MCF-7-specific. We also detected significant enrichment of CCVs in active promoters in ER-positive cells (defined by H3K4me3 marks in T-47D), although the evidence for this effect was weaker than for distal regulatory elements (defined by H3K27ac marks in MCF-7, **Figure 3B**). Only ER-positive CCVs were significantly enriched in T-47D active promoters. Conversely, CCVs were depleted among repressed gene-regulatory elements (defined by H3K27me3 marks) in normal breast (**Figure 3B**). As a control, we performed similar analyses with autoimmune disease CCVs ^29^ (Methods) and relevant B and T cells (**Figures 3B-E**). The strongest evidence of enrichment of breast cancer CCVs was found at regulatory regions active in ER-positive cells (**Figure 3B**), whereas enrichment of autoimmune CCVs was in regulatory regions active in B and T cells (**Figure 3E**). We also compared the enrichment of our CCVs in enhancer-like and promoter-like regions (defined by ENCODE; **Supplementary Figure 1B**). The strongest evidence of enrichment of ER-positive CCVs in enhancer-like regions was found in MCF-7 cells, the only ER-positive cell line in ENCODE (**Supplementary Figure 1B**). These results highlight both the tissue- and disease-specificity of these histone marked gene regulatory regions.
iv. We observed significant enrichment of CCVs in the binding sites for 40 TFBSs determined by ChIP-Seq (**Figures 3F-H**). The majority of the experiments were performed in ER-positive cell lines (90 TFBSs, 20 with data in ER-negative cell lines, 76 in ER-positive cell lines, and 16 in normal breast). These TFBSs overlap each other and histone marks of active regulatory regions (**Supplementary Figure 2**). Enrichment in five TFBSs (ESR1, FOXA1, GATA3, TCF7L2, E2F1) has been previously reported ^2,30^. All 40 TFBSs were significantly enriched in ER-positive CCVs (**Figure 3F**), seven were also enriched in ER-negative CCVs and nine in ER-neutral CCVs (**Figures 3G-H**). ESR1, FOXA1, GATA3 and EP300 TFBSs were enriched in all CCV ER-subtypes. However, the enrichment for ESR1, FOXA1 or GATA3 was stronger for ER-positive CCVs than for ER-negative or ER-neutral.

**Figure 3.**
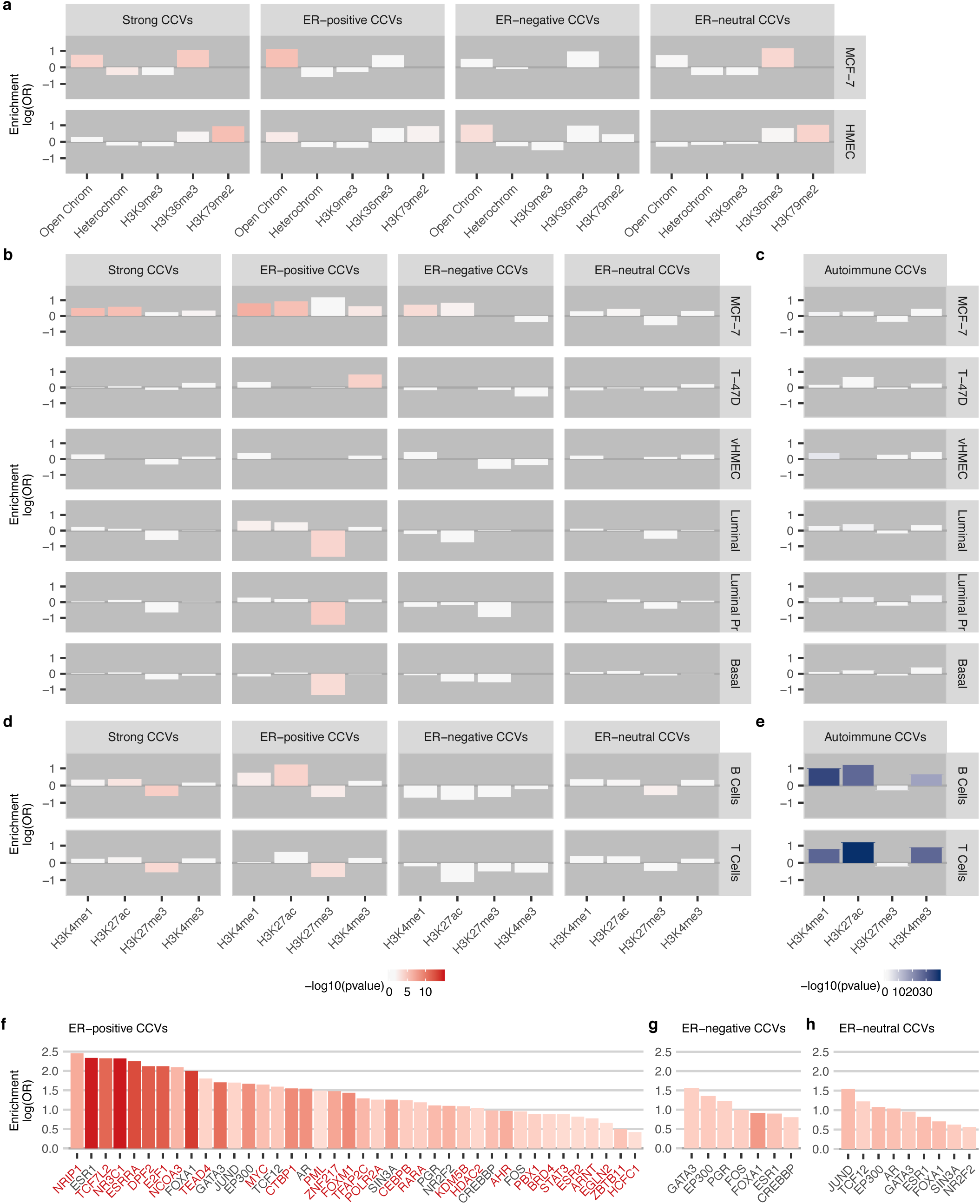
Overlap of CCVs with gene regulatory regions gene bodies and transcription factor binding sites. (a) Breast cancer CCVs overlap with chromatin states and broad breast cells epigenetic marks. (b) Breast cancer CCVs overlap with breast cells epigenetic marks. (c) Autoimmune CCVs overlap with breast cells epigenetic marks. (d) Breast cancer CCVs overlap with autoimmune-related epigenetic marks. (e) Autoimmune CCVs overlap with autoimmune-related epigenetic marks. (f) Significant ER-positive CCVs overlap with transcription factors binding sites. TFBSs found significant for ER-positive CCVs are highlighted in red (x axis labels). (g) Significant ER-negative CCVs overlap with transcription factors binding sites. (h) Significant ER-neutral CCVs overlap with transcription factors binding sites. Strong column: analysis with all CCVs at strong signals. ER-positive, ER-negative, ER-neutral: analysis of CCVs at strong signals stratified by phenotype.

### CCVs significantly overlap consensus transcription factor binding motifs

We investigated whether CCVs were also enriched within consensus TF binding motifs by conducting a motif-search within active regulatory regions (ER-positive CCVs at H3K4me1 marks in MCF-7). We identified 30 motifs, from eight TF families, with enrichment in ER-positive CCVs (FDR 10%, **Supplementary Table 4A**) and a further five motifs depleted among ER-positive CCVs. To assess whether the motifs appeared more frequently than by chance at active regulatory regions overlapped by our ER-positive CCVs, we compared motif-presence in a set of randomized control sequences (Methods). Thirteen of 30 motifs were more frequent at active regulatory regions with ER-positive CCV enrichment; these included seven homeodomain motifs and two fork head factors (**Supplementary Table 4B**).

When we looked at the change in predicted binding affinity, 57 ER-positive signals (86%) included at least one CCV predicted to modify the binding affinity of the enriched TFBSs (≥2-fold, **Supplementary Table 4C**). Forty-eight ER-positive signals (73%) had at least one CCV predicted to modify the binding affinity >10-fold. This analysis validates previous reports of breast cancer causal variants that alter DNA binding affinity for FOXA1 ^3,30^

### Bayesian fine-mapping incorporating functional annotations and linkage disequilibrium

As an alternative statistical approach, we applied PAINTOR ^31^ to the same 128 regions (**Figure 1**). In brief, PAINTOR integrates genetic association results, linkage disequilibrium (LD) structure, and enriched genomic features in an empirical Bayes framework and derives the posterior probability (PP) of each variant being causal, conditional on available data. We identified seven variants with high posterior probability (HPP ≥ 80%) of being causal for all breast cancer types and ten for the ER-positive subtype (**Table 1**); two of these had HPP > 80% for both ER-positive and all breast cancer types. These 15 HPP variants (HPPVs; ≥ 80%) were distributed across 13 regions. We also identified an additional 35 variants in 25 regions with HPP (≥ 50% and < 80%) for ER-positive, ER-negative, or overall breast cancer (**Figure 2G**).

Consistent with the CCV analysis, we found evidence that most regions contained multiple HPPVs; the sum of PP across all variants in a region (an estimate of the number of distinct causal variants in the region) was > 2.0 for 84/86 regions analyzed for overall breast cancer, with a maximum of 16.1 and a mean of 6.4. For ER-positive cancer, 46/47 regions had total PP > 2.0 (maximum 18.3, mean 6.5) and for ER-negative, 17/23 regions had total PP > 2.0 (maximum 9.1, mean 3.2).

Although for many regions we were not able to identify HPP variants, we were able to reduce the proportion of variants needed to account for 80% of the total PP in a region to under 5% for 65 regions for overall, 43 for ER-positive, and 18 for ER-negative breast cancer (**Figures S3A-C**). PAINTOR analyses were also able to reduce the set of likely causal variants in many cases. After summing the PPs for CCVs in each of the overall breast cancer signals, 39/100 strong-evidence signals had a total PP > 1.0. The number of CCVs in these signals ranged from 1 to 375 (median 24), but the number of variants needed to capture 95% of the total PP in each signal ranged from 1 to 115 (median 12), representing an average reduction of 43% in the number of variants needed to capture the signal.

PAINTOR and CCV analyses were generally consistent, yet complementary. Only 3.3% of variants outside of the set of strong-signal CCVs for overall breast cancer had PP > 1%, and only 48 (0.013%) of these had PP > 30% (**Supplementary Figure 3D**). At ER-positive and ER-negative signals respectively, 3.1% and 1.6% of the non-CCVs at strong signals had PP > 1%, and 40 (0.019%) and 3 (0.003%) of these had PP > 30% (**Figures S3E-F**). For the non-CCVs at strong-evidence signals with PP > 30%, the relatively high PP may be driven by the addition of functional annotation. The incorporation of functional annotation (relative to a PAINTOR model with no functional annotation) more than doubled PP for 64/88 variants.

### CCVs co-localize with variants controlling local gene expression

We used four breast-specific expression quantitative trait loci (eQTL) data sets; to identify variants associated with differences in gene expression (eVariants): tumor tissue from the Nurses’ Health Study (NHS) ^32^ and The Cancer Genome Atlas (TCGA) ^33^, and normal breast tissue from the NHS and the Molecular Taxonomy of Breast Cancer International Consortium (METABRIC) ^34^. We then examined the overlap of eVariants (defined as an eVariant within two orders of magnitude of the most significant eVariant for a particular gene) with CCVs (Methods). There was significant overlap of CCVs with eVariants from both the NHS normal and breast cancer tissue studies (normal breast OR = 2.70, p-value = 1.7×10^−5^; tumor tissue OR = 2.34, p-value = 2.6×10^−4^; **Supplementary Table 3**). ER-neutral CCVs overlapped with eVariants in normal tissue more frequently than did ER-positive and ER-negative CCVs (OR_ER-neutral_ = 3.51, p-value = 1.3×10^−5^). Cancer risk CCVs overlapped credible eVariants in 128/205 (62%) signals in at least one of the datasets (**Tables S5A-B**). Sixteen additional variants with PP ≥ 30%, not included among the CCVs, also overlapped with a credible eVariant (**Tables S5A-B**).

### Transcription factors and known somatic breast cancer drivers are overrepresented among prioritized target genes

We assumed that causal variants function by affecting the behavior of a local target gene. However, it is challenging to define target genes or to determine how they may be affected by the causal variant. Few potentially causal variants directly affect protein coding: we observed 67/5,375 CCVs, and 19/137 HPPVs (≥ 30%) in protein-coding regions. Of these, 33 (0.61%) were predicted to create a missense change, one a frameshift, and another a stop-gain, while 30 were synonymous (0.59%, **Supplementary Table 5C**). Four hundred and ninety-nine CCVs at 94 signals, and four additional HPPV (≥ 30%), are predicted to create new splice sites or activate cryptic splice sites in 126 genes (**Supplementary Table 5D**). These results are consistent with previous observations that majority of common susceptibility variants are regulatory.

We applied an updated version of our pipeline INQUISIT - **in**tegrated expression **qu**antitative trait and ***i****n-****s****ilico* prediction of GWAS **t**argets) ^2^ to prioritize potential target genes from 5,375 CCVs in strong signals and all 138 HPPVs (≥ 30%; **Supplementary Table 2C**). The pipeline predicted 1,204 target genes from 124/128 genomic regions examined. As a validation we examined the overlap between INQUISIT predictions and 278 established breast cancer driver genes ^35-39^. Cancer driver genes were over-represented among high confidence (Level 1) targets; a 6-fold increase over expected from CCVs and 16-fold from HPPVs; p-value= 1×10^−6^; **Supplementary Figure 4A**). Notably, ten cancer driver genes (*CCND1, CHEK2, ESR1, FGFR2, GATA3, MAP3K1, MYC, TBX3, XBP1* and *ZFP36L1)* were predicted from the HPPVs derived from PAINTOR. Cancer driver gene status was consequently included as an additional weighting factor in the INQUISIT pipeline. TF genes ^40^ were also enriched amongst high-confidence targets predicted from both CCVs (2-fold, p-value = 4×10^−4^) and HPPVs (3-fold, p-value = 7×10^−3^, **Supplementary Figure 4A**).

In total INQUISIT prioritized 178 high-confidence target genes (**Table 2, Supplementary Table 6**). Significantly more genes were targeted by multiple independent signals (N = 165) than expected by chance (p-value = 4.6×10^−8^, **Supplementary Figure 4B, Figure 4A**). Six high-confidence predictions came only from HPPVs, although three of these (*IGFBP5, POMGNT1* and *WDYHV1*) had been predicted at lower confidence from CCVs. Target genes included 20 that were prioritized via potential coding/splicing changes (**Supplementary Table 7**), nine via promoter variants (**Supplementary Table 8**), and 166 via distal regulatory variants (**Supplementary Table 9**). We illustrate genes prioritized via multiple lines of evidence in **Figure 4A**.

**Table 2.**
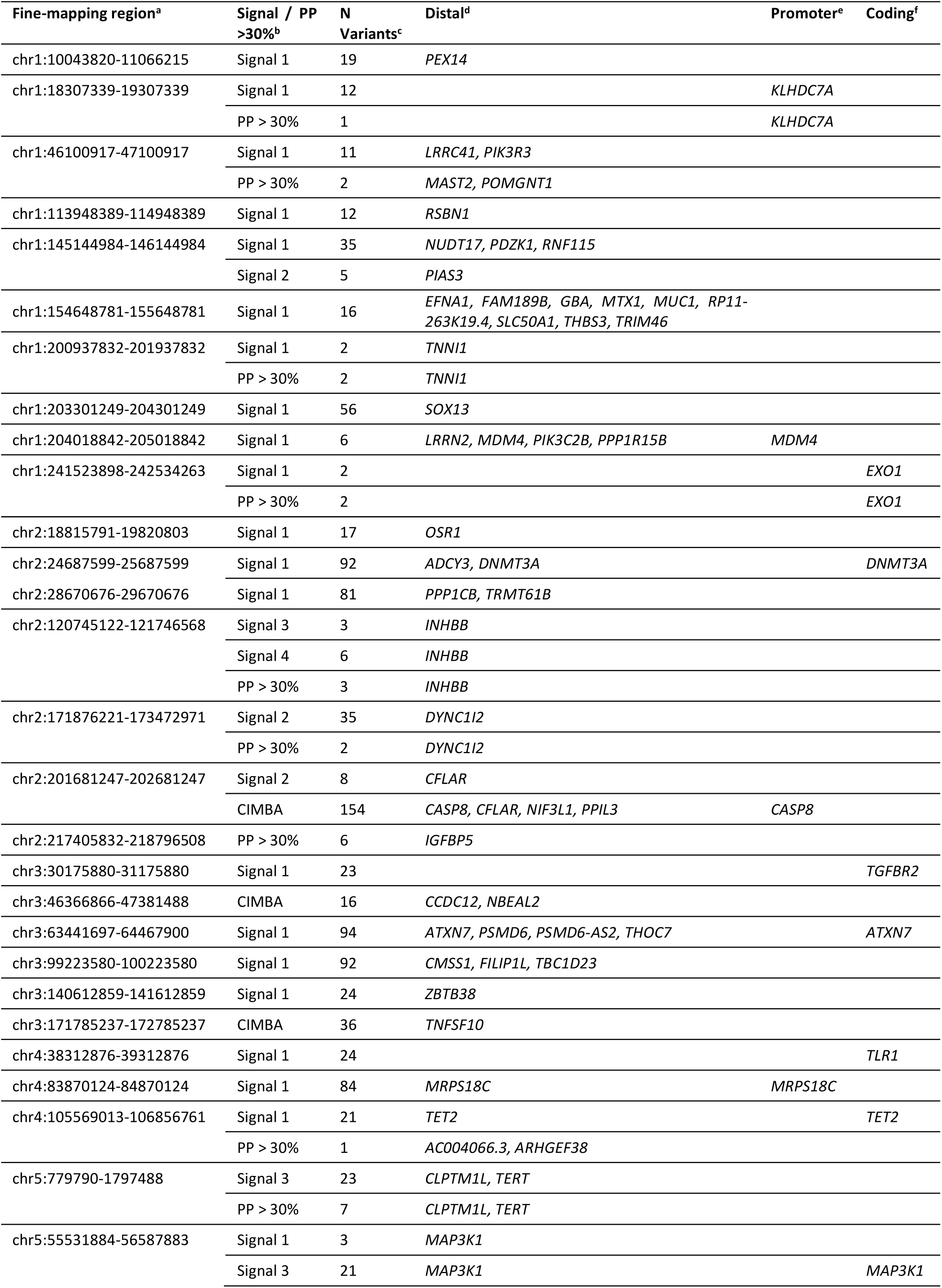
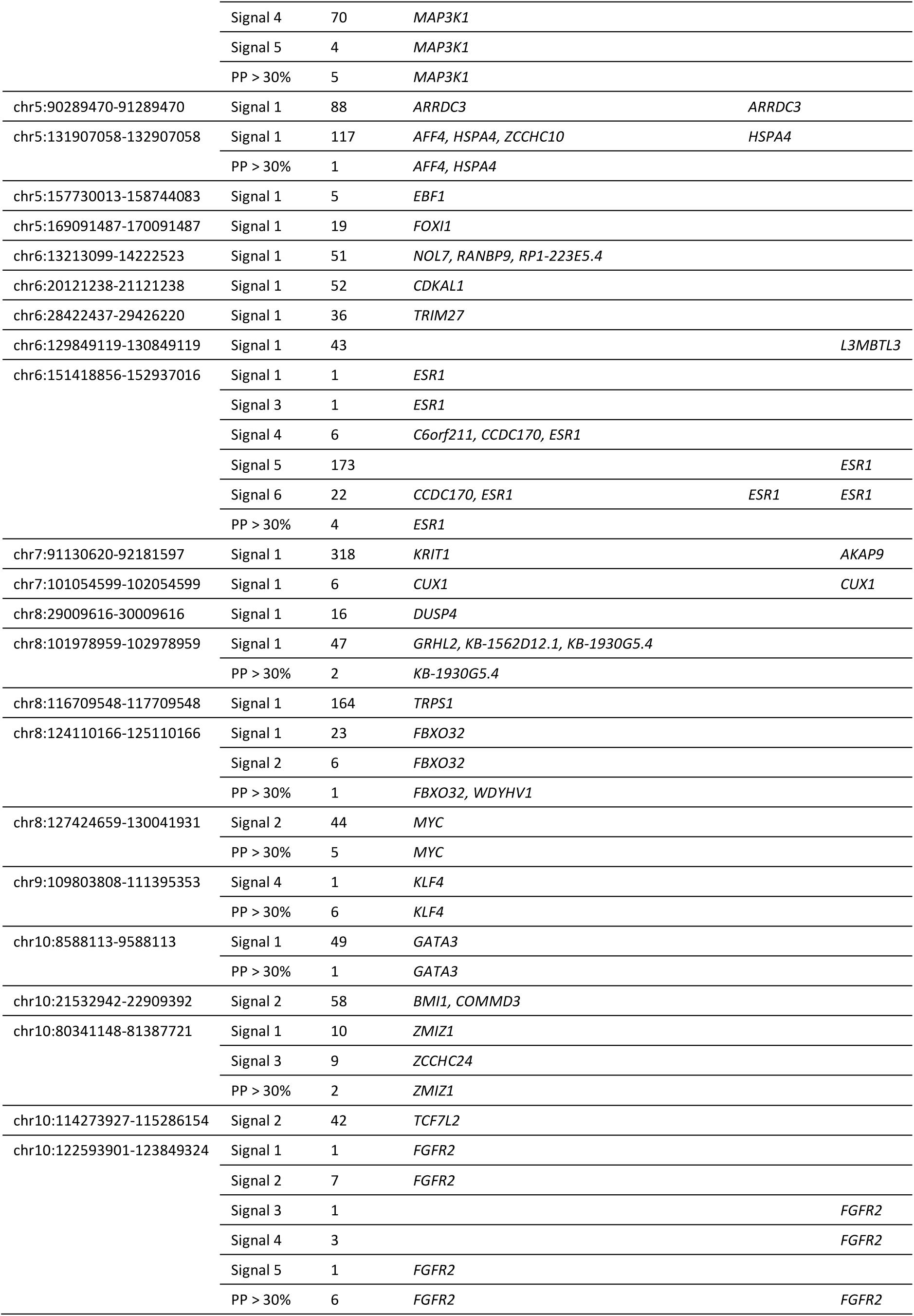
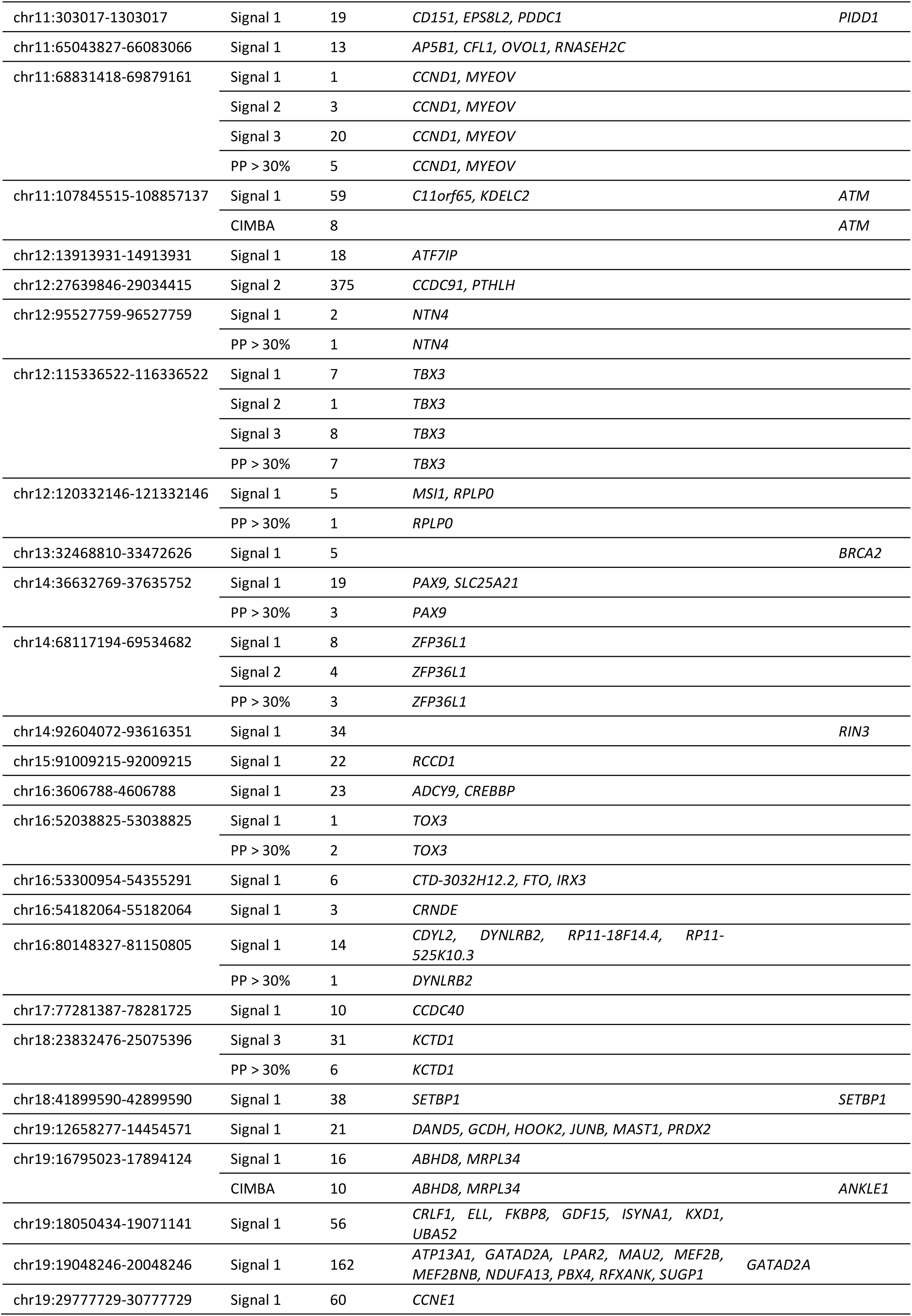
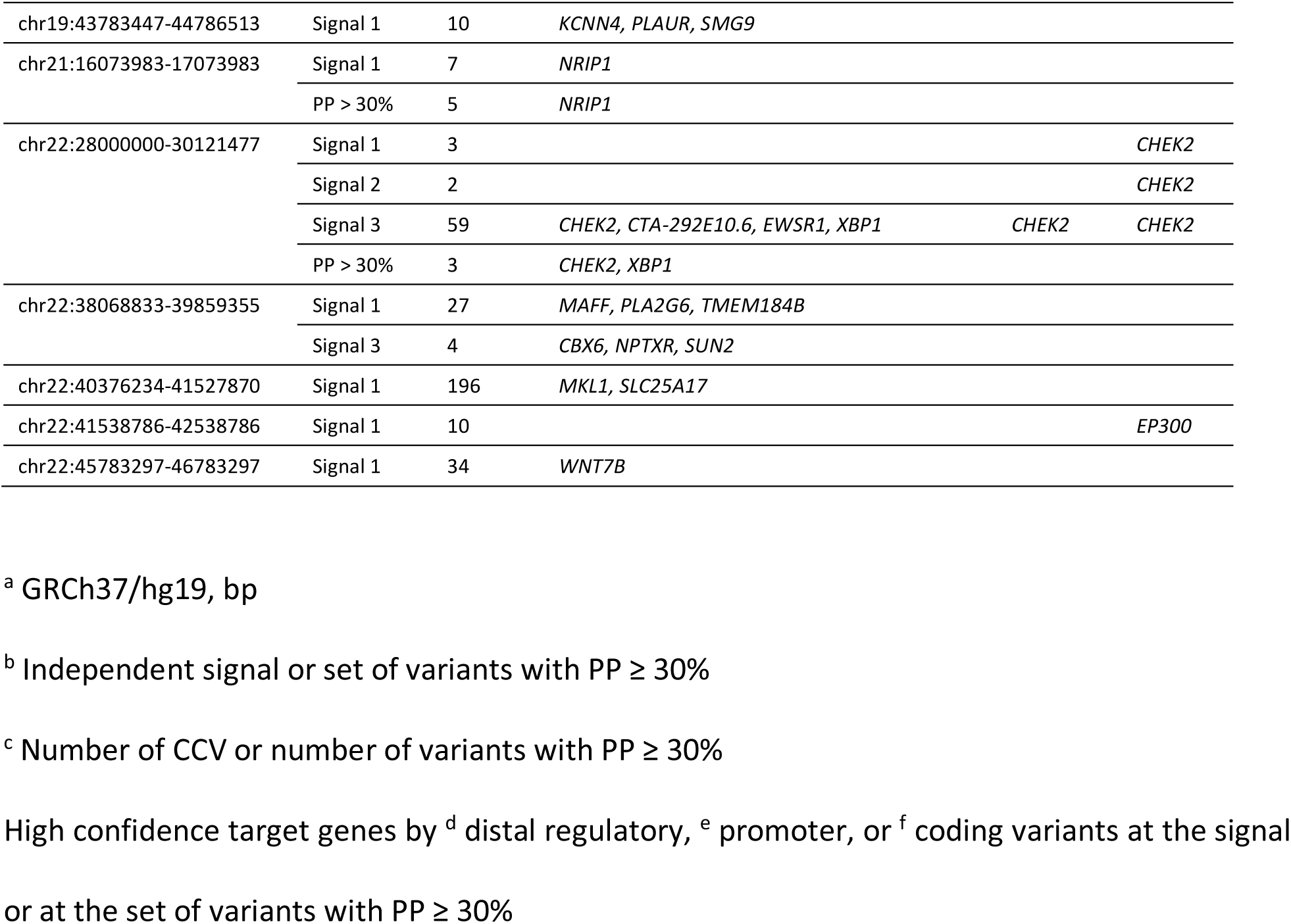
Regions in which target genes are predicted with high confidence.

**Figure 4.**
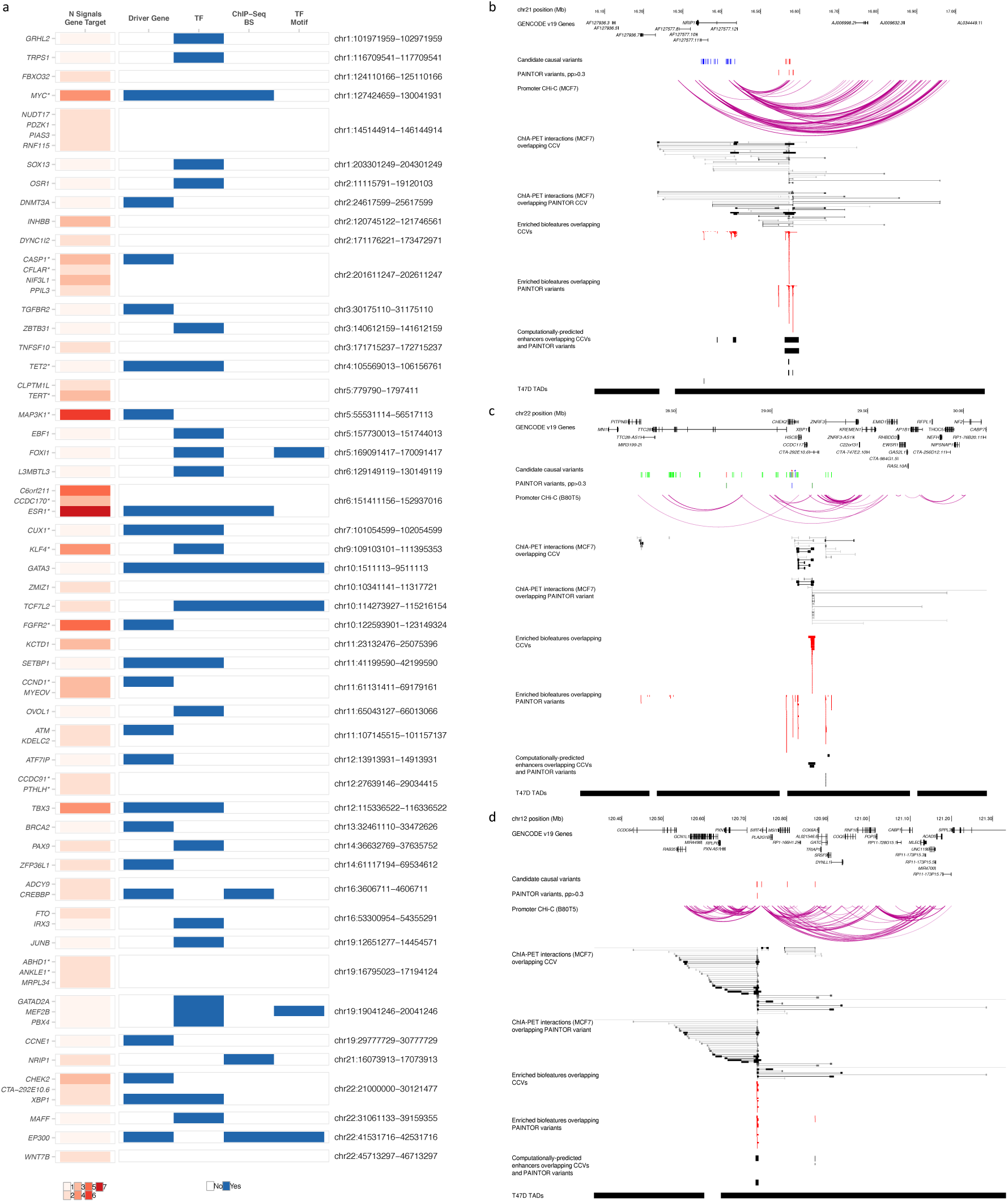
Predicted target genes are enriched in known breast cancer driver genes and transcription factors. (a) 79 target genes that fulfil at least one of the following criteria: are targeted by more than one independent signal, are known driver genes, transcription factor genes, or their binding sites (ChIP-Seq BS) or consensus motif (TF Motif) are significantly overlapped by CCVs. *Genes with published functional follow up. (b-d) In each panel, CCVs and PAINTOR variants with posterior probability >0.3 are shown, with independent signals in different colors. Chromatin interactions are shown as arcs (Capture Hi-C from selected breast cell lines) or boxes connected by lines, colored with gray-scale according to interaction score (ENCODE ChIA-PET). Biofeatures which overlap CCVs from the global genomic enrichment analysis are depicted as red boxes. Computationally predicted enhancers including PreSTIGE, FANTOM5 and super-enhancers which overlap risk variants are represented by black boxes. High confidence INQUISIT target gene predictions include *NRIP1* (b), *CHEK2* and *XBP1* (c), and *RPLP0* and *MSI1* (d).

Three examples of INQUISIT using genomic features to identify predict target genes. Based on capture Hi-C and ChIA-PET chromatin interaction data, *NRIP1* is a predicted target of intergenic CCVs and HPPVs at chr21q21 (**Figure 4B**). Multiple target genes were predicted at chr22q12, including the driver genes *CHEK2* and *XBP1* (**Figure 4C**). A third example at chr12q24.31 is a more complicated scenario with two Level 1 targets: *RPLP0*^41^ and a modulator of mammary progenitor cell expansion, *MSI1*^42^ (Figure 4D).

### Target gene pathways include DNA integrity-checkpoint, apoptosis, developmental processes and the immune system

We performed pathway analysis to identify common processes using INQUSIT high confidence target protein-coding genes (**Figure 5A**) and identified 457 Gene Ontology (GO) terms and 265 pathways at an FDR of 5% (**Supplementary Table 10**). These were grouped into 99 themes by common ancestor GO terms, pathways, or TF classes (**Figure 5B**). We found that 24% (14/58) of the ER-positive target genes were classified within developmental process pathways (including mammary development), 17% in immune system and a further 16% in cell differentiation pathways. Of genes targeted by ER-neutral signals, 19% (16/85) were classified in developmental process pathways, 18% in apoptotic process, and a further 16% in immune system pathways. The top themes of genes targeted by ER-negative signals were DNA integrity checkpoint (20%, 7/35), immune system, response to stimulus, and apoptotic processes, each containing 17% (6/35) of the genes.

**Figure 5.**
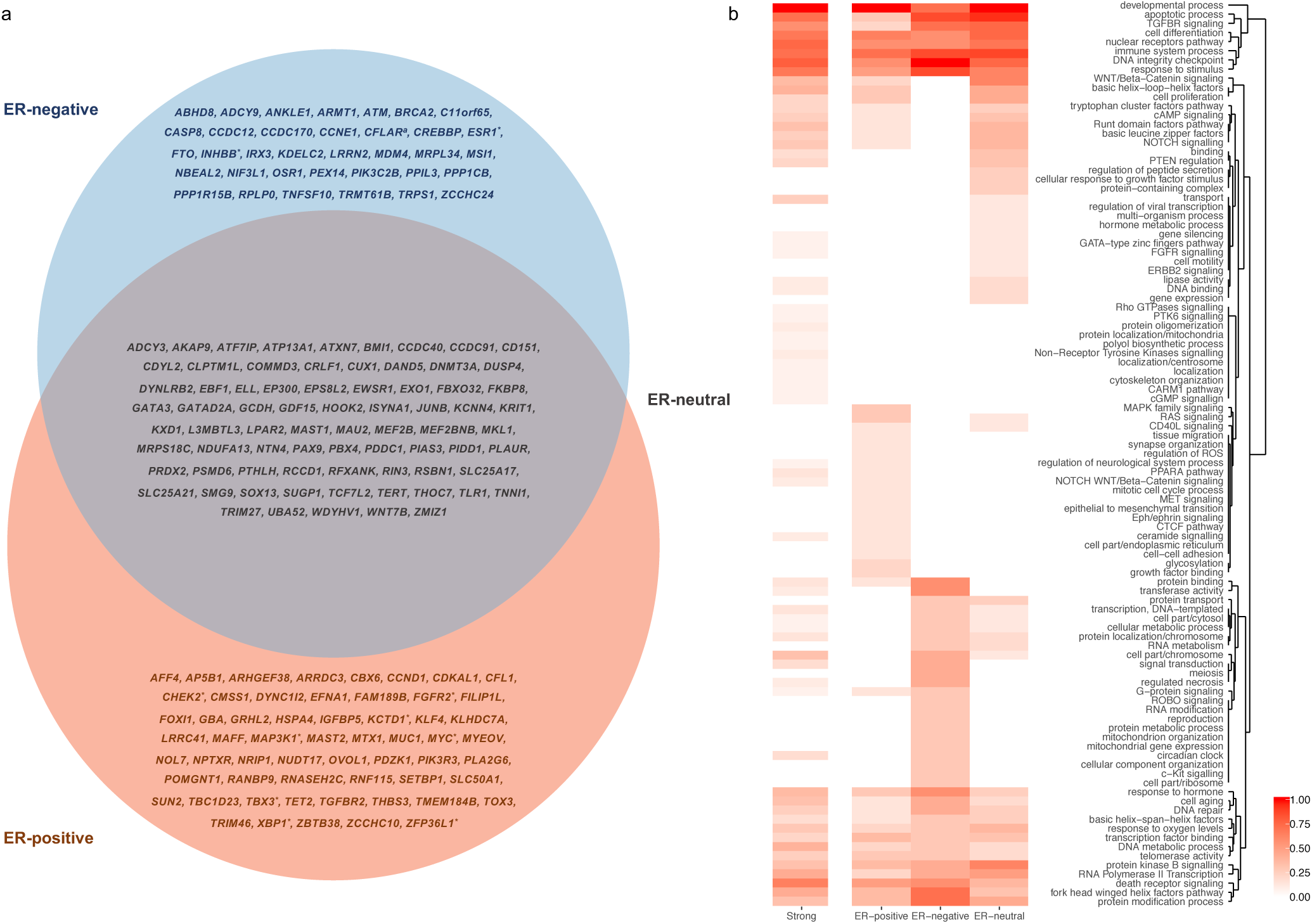
Predicted target genes by phenotype and significantly enriched pathways. (a) Venn diagram showing the associated phenotype (ER-positive, ER-negative, ER-neutral) for the Level 1 target genes, predicted by the CCVs and HPPVs. * ER-positive or ER-negative target genes also targeted by ER-neutral signals. (b) Heatmap showing clustering of pathway themes over-represented by INQUISIT Level 1 target genes. Color represents the relative number of genes per phenotype within enriched pathways, grouped by common themes. ER-positive, ER-negative, ER-neutral, and all phenotypes together (strong).

Novel pathways revealed by this study include TNF-related apoptosis-inducing ligand (TRAIL) signaling, the AP-2 transcription factors pathway, and regulation of IκB kinase/NF-κB signaling. Of note, the latter of these is specifically overrepresented among ER-negative target genes. We also found significant overrepresentation of additional carcinogenesis-linked pathways including cAMP, NOTCH, PI3K, RAS, WNT/Beta-catenin, and of receptor tyrosine kinases signaling, including FGFR, ERBB, or TGFBR ^43-47^. Also of note, our target genes are significantly overrepresented in DNA damage checkpoint, DNA repair pathways, as well as programmed cell death pathways, such as apoptotic process, regulated necrosis, and death receptor signaling-related pathways.

## DISCUSSION

We have performed multiple, complementary analyses on 150 breast cancer associated regions, originally found by GWAS, and identified 362 independent risk signals, 205 of these with high confidence (p-value < 10^−6^). We observed most regions contain multiple independent signals, the greatest number (nine) in the region surrounding *ESR1* and its co-regulated genes, and on 2q35, where *IGFBP5* appears to be a key target. We have used two complementary approaches to identify likely causal variants within each region: a Bayesian approach, PAINTOR, which integrated genetic associations, LD and informative genomic features, confirmed most associations found by the more traditional, multinomial regression approach, and also identified additional variants. Specifically, the Bayesian method highlighted 15 variants that are highly likely to be causal (HPP ≥ 80%). From these approaches we have identified a single variant, likely to be causal, at each of 34/205 signals (**Table 1**). Of these, only rs16991615 (*MCM8* p.E341K) and rs7153397 (*CCDC88C,* a cryptic splice-donor site) were predicted to affect protein-coding sequences. However, in other signals we also identified four coding changes previously recognized as deleterious, including the stop-gain rs11571833 (*BRCA2* p.K3326*, Meeks et al., 2016)^48^ and two *CHEK2* coding variants; the frameshift rs555607708 ^49,50^, and a missense variant, rs17879961 ^51,52^. In addition, a splicing variant, rs10069690, in *TERT* results in the truncated protein INS1b ^19^, decreased telomerase activity, telomere shortening, and increased DNA damage response ^53^

Having identified potential causal variants within each signal, we aimed to uncover their functions at the DNA level and as well as trying to predict their target gene(s). Looking across all 150 regions, a notable feature is that many likely causal variants implicated in ER-positive cancer risk, lie in gene-regulatory regions marked as open and active in ER-positive breast cells, but not in other cell types. Moreover, a significant proportion of potential causal variants overlap the binding sites for transcription factor proteins (n=40 from ChIP-Seq) and co-regulators (n=64 with addition of computationally derived motifs). Furthermore, nine proteins also appear in the list of high-confidence target genes, hence the following genes and their products have been implicated by two different approaches: *CREBBP, EP300, ESR1, FOXI1, GATA3, MEF2B, MYC, NRIP1* and *TCF7L2*. Most of these proteins already have established roles in estrogen signaling. *CREBBP, EP300, ESR1, GATA3*, and *MYC* are also known cancer driver genes that are frequently somatically mutated in breast tumors.

In contrast to ER-positive signals, we identified fewer genomic features enriched in ER-negative signals. This may reflect the common molecular mechanisms underlying their development, but the power of this study was limited, despite including as many patients with ER-negative tumors as possible from the BCAC and CIMBA consortia. Less than 20% of genomic signals confer a greater risk of ER-negative cancer and there is little publicly available ChIP-Seq data on ER-negative breast cancer cell lines. The heterogeneity of ER-negative tumors may also have limited our power. Nevertheless, we have identified 35 target genes for ER-negative likely causal variants. Some of these already had functional evidence supporting their role: including *CASP8*^54^ and *MDM4*^55^. Most targets, however, currently have no reported function in ER-negative breast cancer development.

Finally, we examined the gene-ontology pathways in which target genes most often lie. Of note, 12% (20/167) of all high-confidence target genes and 17% of ER-negative target predictions are in immune system pathways. Among the significantly enriched pathways were T cell activation, interleukin signaling, Toll-like receptor cascades, and I-κB kinase/NF-κB signaling, as well as processes leading to activation and perpetuation of the innate immune system. The link between immunity, inflammation and tumorigenesis has been extensively studied ^56^, although not primarily in the context of susceptibility. Four ER-negative high confidence target genes (*CASP8, CFLAR, ESR1, TNFSF10*) lie in the I-κB kinase/NF-κB signaling pathway. Interestingly, ER-negative cells have high levels of NF-kB activity when compared to ER-positive ^57^. A recent expression–methylation analysis on breast cancer tumor tissue also identified clusters of genes correlated with DNA methylation levels, one enriched in ER signaling genes, and a second in immune pathway genes ^58^.

These analyses provide strong evidence for more than 200 independent breast cancer risk signals, identify the plausible cancer variants and define likely target genes for the majority of these. However, notwithstanding the enrichment of certain pathways and transcription factors, the biological basis underlying most of these signals remains poorly understood. Our analyses provide a rational basis for such future studies into the biology underlying breast cancer susceptibility.

## METHODS

### Study samples

Epidemiological data for European women were obtained from 75 breast cancer case-control studies participating in the Breast Cancer Association Consortium (BCAC) (cases: 40,285 iCOGS, 69,615 OncoArray; cases with ER status available: 29,561 iCOGS, 55,081 OncoArray); controls: 38,058 iCOGS, 50,879 OncoArray). Details of the participating studies, genotyping calling and quality control are given in ^2,22,23^, respectively. Epidemiological data for *BRCA1* mutation carriers were obtained from 60 studies providing data to the Consortium of Investigators of Modifiers of *BRCA1* and *BRCA2* (CIMBA) (affected 1,591 iCOGS, 7,772 OncoArray; unaffected 1,665 iCOGS, 7,780 OncoArray). This dataset has been described in detail previously ^1,59,60^. All studies provided samples of European ancestry. Any non-European samples were excluded from analyses.

### Variant selection, genotyping and imputation

A dense set of variants surrounding susceptibility variants known at the time of design was selected for genotyping on the iCOGS array. To select markers for comprehensive interrogation of these densely-genotyped regions, all known variants from the March 2010 release of the 1000 Genomes Project with MAF > 0.02 in Europeans were identified, and all variants that were correlated with the published GWAS variants at r2 > 0.1 together with a set of variants designed to tag all remaining variants at r2 > 0.9 were selected. Twenty-one regions were densely genotyped with iCOGS (http://ccge.medschl.cam.ac.uk/files/2014/03/iCOGS_detailed_lists_ALL1.pdf). In total 26,978 iCOGS genotyped variants across the 152 mapped regions passed QC criteria. Similarly, all known associated risk variants at the time of the design of the OncoArray were included in the genotyping array. Variants for fine-mapping were selected defining a 1Mb interval surrounding a known hit. Overlapping regions were merged into a single interval including 500kb either side of each lead signal. From among designable variants, all variants correlated with the known hits at r^2^ > 0.6, all variants from lists of potentially functional variants, defined through RegulomeDB, and a set of variants designed to tag all remaining variants at r^2^ > 0.9 were selected. Seventy-three regions were densely genotyped with OncoArray. In total 58,339 OncoArray genotyped variants across the 152 mapped regions passed QC criteria.

We imputed genotypes for the remaining variants at the genomic regions defined in the OncoArray design using IMPUTE2 ^61^, and the October 2014 release of the 1000 Genomes Project as a reference. To improve the accuracy at low frequency variants we used the standard IMPUTE2 MCMC algorithm for follow-up imputation (more detailed description of the parameters used can be found in ^21^)We genotyped or successfully imputed 639,118 variants all having imputation info score ≥ 0.3 and minor allele frequencies (MAF) ≥ 0.001 in both the iCOGS and OncoArray datasets. Imputation summaries, and coverage for each of the analyzed regions stratified by allele frequency can be found in **Supplementary Table 1B**.

### BCAC Statistical analyses

Per-allele odds ratios (OR) and standard errors (SE) were estimated for each variant using logistic regression. We ran this analysis separately for iCOGS and OncoArray, and separately for overall, ER-positive and ER-negative breast cancer. The association between each variant and breast cancer risk was adjusted by study (iCOGS) or country (OncoArray), and eight (iCOGS) or ten (OncoArray) ancestry-informative principal components. The statistical significance for each variant was derived using a Wald test.

#### Defining the significance threshold

To establish an appropriate significance threshold for independent signals, all variants evaluated in the meta-analysis were included in logistic forward selection regression analyses for overall breast cancer risk in iCOGS cohort, run independently for each of the regions. We evaluated five p-value thresholds for inclusion: < 1×10^−4^, < 1×10^−5^, < 1×10^−6^, < 1×10^−7^, and < 1×10^−8^. The most parsimonious iCOGS models were tested in OncoArray, and the false discovery rate (FDR) at 1% level for each one of the thresholds was estimated using the Benjamini-Hochberg procedure ^62^. At a 1% FDR threshold, 72% of variants of variants significant at p<10^−4^ replicated on the OncoArray, whereas 94% of variants significant at p<10^−6^ replicated. Based on these analyses, two categories were defined: strong-evidence signals (conditional p-values <10^−6^ when the signal is included in the final model fit using the inclusion threshold p<10^−4^), and moderate-evidence signals (conditional p-values <10^−4^ and ≥10^−6^ in the final model).

#### Identification of independent signals

To identify independent signals, we ran multinomial stepwise regression analyses, separately in iCOGS and OncoArray, for all the variants displaying evidence of association (N_variants_ = 202,749). We selected two sets of well imputed variants (imputation info score ≥ 0.3 in both iCOGS and OncoArray): (a) common and low frequency variants (MAF ≥ 0.01) with a threshold for inclusion of logistic regression p-value ≤0.05 in either the iCOGS or OncoArray datasets for at least one of the three phenotypes: overall, ER-positive and ER-negative breast cancer; and (b) rarer variants (MAF ≥ 0.001 and < 0.01), with a threshold for inclusion of a logistic regression p-value ≤ 0.0001. The same parameters used for adjustment in logistic regression were used in the multinomial regression analysis (R function *multinom*). The multinomial regression estimates were combined using a fixed-effects meta-analysis weighted by the inverse variance ^63^. Variants with the lowest conditional p-value from the meta-analysis of both European cohorts at each step were included into the multinomial regression model. However, if the new variant to be included in the model caused collinearity problems due to be highly correlated with an already selected variant, or showed high heterogeneity (p-value < 10^−4^) between iCOGS and OncoArray after being conditioned by the variant(s) in the model; we dropped the new variant and repeated this process.

The main signal at 105 of 152 evaluated regions demonstrated genome-wide significance, while 44 were marginally significant (9.89×10^−5^ ≥ p-value > 5×10^−8^). For two regions there were no variants significant at p<10^−4^ (chr14:104712261-105712261; rs10623258 multinomial regression p-value = 2.32×10^−4^; chr19:10923703-11923703, rs322144, multinomial regression p-value = 3.90×10^−3^). There are four main differences in the dataset used in the analysis and that used in the previous paper that account for these differences : (i) our previous paper ^2^ included data from 11 additional GWAS studies (14,910 cases and 17,588 controls) that have not been included in the present analysis in order to minimize differences in array coverage; (ii) the present analysis evaluated the risk of ER-positive and ER-negative disease concurrently, whereas in our previous paper the outcome was overall breast cancer risk. ER status was available for only 73% of the iCOGS and 79% of the OncoArray breast cancer cases (iii) for the set of samples genotyped with both arrays, ^2^ included iCOGS genotypes, while this study includes OncoArray genotypes to maximize the number of samples genotyped with a larger coverage; and (iv) imputation parameters at fine mapping-regions were modified to improve the imputation accuracy of less frequent variants.

#### Selection of a credible set of variants candidates to be causal

Variants with p-values within two orders of magnitude of the top variant for each signal, after adjusting for the index variant of preceding signals, were considered as the credible set of candidates to drive the association at each independent signal (CCVs) ^24^We defined the best model, among those including one CCVs for each signal, by iteratively conditioning on the selected CCVs in the other signals. For ten fine mapping regions (chr4:175328036-176346426, chr5:55531884-56587883, chr6:151418856-152937016, chr8:75730301-76917937, chr10:80341148-81387721, chr10:122593901-123849324, chr12:115336522-116336522, chr14:36632769-37635752, chr16:3606788-4606788, chr22:38068833-39859355) a model including the lowest conditioned p-value variants (index variant) at each signal could not be found. In that case we estimated the best model as the model with the largest chi-square, from all the possible combinations of credible variants. Variants in strong signals were conditioned by the additional strong confidence signals whereas variants for moderate confidence signals were conditioned by all the independent signals. We then redefined the credible set of candidate variants for each signal using the conditional p-values by the additional independent signals. Thus, variants with p-values within two orders of magnitude of the top variant for each signal, after adjusting for all the index variant of the additional signals at the locus, were considered as the final credible set of candidates.

#### Case-only analysis

Differences in the effect size between ER-positive and ER-negative disease for each index independent variant were assessed using a case-only analysis. We performed logistic regression with ER status as the dependent variable, and the lead variant at each strong signal in the fine mapping region as the independent variables. We use FDR (5%) to adjust for multiple testing.

### OncoArray-only stepwise analysis

To evaluate whether the lower coverage in iCOGS could affect the identification of independent signals, we ran stepwise multinomial regression using only the OncoArray dataset. We identified 249 independent signals. Ninety-two signals, in 67 fine mapping regions, achieved a genome-wide significance level (conditional p-value < 5×10-8). Two hundred and five of these signals were also identified in the meta-analysis with iCOGS. Nine independent variants across ten regions were not evaluated in the combined analysis due to their low imputation info score in iCOGS. Out of these nine signals, two signals would be classified as main primary signals, rs114709821 at region chr1:145144984-146144984 (OncoArray imputation info score = 0.72), and rs540848673 at region chr1:149406413-150420734 (OncoArray imputation info score = 0.33). Given the low number of additional signals identified in the OncoArray dataset alone, all analyses were based on the combined iCOGS/OncoArray dataset.

### CIMBA statistical analysis

CIMBA provided data from 60 retrospective cohort studies consisting of 9,445 unaffected and 9,363 affected female *BRCA1* mutation carriers of European ancestry. Unconditional (i.e. single variant) analyses were performed using a score test based on the retrospective likelihood of observing the genotype conditional on the disease phenotype ^64,65^. Conditional analyses, where more than one variant is analyzed simultaneously, cannot be performed in this score test framework. Therefore, conditional analyses were performed by Cox regression, allowing for adjustment of the conditionally independent variants identified by the BCAC/DRIVE analyses. All models were stratified by country and birth cohort, and adjusted for relatedness (unconditional models used kinship adjusted standard errors based on the estimated kinship matrix; conditional models used cluster robust standard errors based on phenotypic family data).

Data from the iCOGS array and the OncoArray were analyzed separately and combined to give an overall *BRCA1* association by fixed-effects meta-analysis. Variants were excluded from further analyses if they exhibited evidence of heterogeneity (Heterogeneity p-value < 1×10-4) between iCOGS and OncoArray, had MAF < 0.005, were poorly imputed (imputation info score < 0.3) or were imputed to iCOGS only (i.e. must have been imputed to OncoArray or iCOGS and OncoArray).

### Meta-analysis of ER-negative cases in BCAC with ***BRCA1* mutation carriers from CIMBA**

*BRCA1* mutation carrier association results were combined with the BCAC multinomial regression ER-negative association results in a fixed-effects meta-analysis. Variants considered for analysis must have passed all prior QC steps and have had MAF≥0.005. All meta-analyses were performed using the METAL software ^66^.Instances where spurious associations might occur were investigated by assessing the LD between a possible spurious association and the conditionally independent variants. High LD between a variant and a conditionally independent variant within its region causes model instability through collinearity and the convergence of the model likelihood maximization may not reliable. Where the association appeared to be driven by collinearity, the signals were excluded.

### eQTL analysis

Total RNA was extracted from normal breast tissue in formalin-fixed paraffin embedded breast cancer tissue blocks from 264 Nurses’ Health Study (NHS) participants ^32^. Transcript expression levels were measured using the Glue Grant Human Transcriptome Array version 3.0 at the Molecular Biology Core Facilities, Dana-Farber Cancer Institute. Gene expression was normalized and summarized into Log_2_ values using RMA (Affymetrix Power Tools v1.18.012); quality control was performed using GlueQC and arrayQualityMetrics v3.24.014. Genome-wide data on variants were generated using the Illumina HumanHap 550 BeadChip as part of the Cancer Genetic Markers of Susceptibility initiative ^67^. Imputation to the 1000KGP Phase 3 v5 ALL reference panel was performed using MACH to pre-phase measured genotypes and minimac to impute.

Expression analyses were performed using data from The Cancer Genome Atlas (TCGA) and Molecular Taxonomy of Breast Cancer International Consortium (METABRIC) projects ^34,38^. The TCGA eQTL analysis was based on 458 breast tumors that had matched gene expression, copy number and methylation profiles together with the corresponding germline genotypes available. All 458 individuals were of European ancestry as ascertained using the genotype data and the Local Ancestry in admixed Populations (LAMP) software package (LAMP estimate cut-off >95% European)^68^. Germline genotypes were imputed into the 1000 Genomes Project reference panel (October 2014 release) using IMPUTE version 2 ^69,70^. Gene expression had been measured on the Illumina HiSeq 2000 RNA-Seq platform (gene-level RSEM normalized counts ^71^), copy-number estimates were derived from the Affymetrix SNP 6.0 (somatic copy-number alteration minus germline copy-number variation called using the GISTIC2 algorithm ^72^), and methylation beta values measured on the Illumina Infinium HumanMethylation450. Expression QTL analysis focused on all variants within each of the 152 genomic intervals that had been subjected to fine-mapping for their association with breast cancer susceptibility. Each of these variants was evaluated for its association with the expression of every gene within 2κMb that had been profiled for each of the three data types. The effects of tumor copy number and methylation on gene expression were first regressed out using a method described previously ^73^. eQTL analysis was performed by linear regression, with residual gene expression as outcome, germline SNP genotype dosage as the covariate of interest and ESR1 expression and age as additional covariates, using the R package Matrix eQTL ^74^.

The METABRIC eQTL analysis was based on 138 normal breast tissue samples resected from breast cancer patients of European ancestry. Germline genotyping for the METABRIC study was also done on the Affymetrix SNP 6.0 array, and gene expression in the METABRIC study was measured using the Illumina HT12 microarray platform (probe-level estimates). No adjustment was implemented for somatic copy number and methylation status since we were evaluating eQTLs in normal breast tissue. All other steps were identical to the TCGA eQTL analysis described above.

### Genomic feature enrichment

We explored the overlap of candidate and excluded variants with 90 transcription factors, 10 histone marks, and DNase hypersensitivity sites in in 15 breast cell lines, and eight normal human breast tissues. We analysed data from the Encyclopedia of DNA Elements (ENCODE) Project ^75,76^, Roadmap Epigenomics Projects ^77^, the International Human Epigenome Consortium ^78^, ^27, 79^, The Cancer Genome Atlas (TCGA) ^33^, the Molecular Taxonomy of Breast Cancer International Consortium (METABRIC) ^34^, ReMap database (We included 241 TF annotations from ReMap (of 2825 total) which showed at least 2% overlap for any of the phenotype SNP sets) ^80^, and other data obtained through the National Center for Biotechnology Information (NCBI) Gene Expression Omnibus (GEO). Promoters were defined following the procedure defined in Pellacani et al. ^79^, that is +/-2Kb from a gene transcription start site, using an updated version of the RefSeq genes (refGene, version updated 2017-04-11)^81^. Transcribed regions were defined using the same version of refSeq genes. lncRNA annotation was obtained from Gencode (v19)^82^

To include eQTL results in the enrichment analysis we (i) identified all the genes for which summary statistics were available, for each study; (ii) defined the most significant eQTL variant for each gene (index eQTL variant, p-value threshold ≤ 5×10-4); (iii) classified variants with p-values within two orders of magnitude of the index eVariant as the credible set of eQTL variants candidate to drive the expression of the gene. Those variants part of at least one eQTL credible set were defined as eVariants. We evaluated the overlap between eQTL credible sets and CCVs (risk variants credible set). We evaluated the enrichment of CCVs for genomic feature using logistic regression, with CCV (vs non-CCV variants) being the outcome. To adjust for the correlation among variants in the same fine mapping region, we used robust variance estimation for clustered observations (R function *multiwaycov*). The associated variants at FDR 5% were included into a stepwise forward logistic regression procedure to select the most parsimonious model. A likelihood ratio test was used to compare multinomial logistic regression models with and without equality effect constraints to evaluate whether there was heterogeneity among the effect sizes for ER-positive, ER-negative or signals equally associated with both phenotypes (ER-neutral).

To validate the disease specificity of the regulatory regions identified through this analysis we follow the same approach for the autoimmune related CCVs from ^29^ (N = 4,192). Variants excluded as candidate causal variants, and within 500 kb upstream and downstream of the index variant for each signal were classified as excluded variants (N = 1,686,484). We then tested the enrichment for both the breast cancer and autoimmune CCVs with breast and T and B cell enhancers. We also evaluated the overlap of our CCVs with ENCODE enhancer-like and promoter-like regions for 111 tissues, primary cells, immortalized cell line, and in vitro differentiated cells. Of these, 73 had available data for both enhancer- and promoter-like regions.

### Transcription binding site motif analysis

We conducted a search to find motif occurrences for the transcription factors significantly enriched in the genomic featured. For this we used two publicly available databases, Factorbook ^83^ and JASPAR 2016 ^84^. For the search using Factorbook we included the motifs for the transcription factors discovered in the cell lines where a significant enrichment was found in our genomic features analysis. We also searched for all the available motifs for *Homo sapiens* at the JASPAR database (*JASPAR CORE 2016, TFBSTools* ^*85*^)Using as reference the USCS sequence (*BSgenome.Hsapiens.USCS.hg19*) we created fasta sequences with the reference and alternative alleles for all the variants included in our analysis plus 20 bp flanking each variant. We used FIMO (version 4.11.2, Grant et al., 2011)^86^ to scan all the fasta sequences searching for the JASPAR and Factorbook motifs to identify any overlap of any of the alleles for each of the variants (setting the p-value threshold to 10^−3^). We subsequently determined whether our CCVs were more frequency overlapping a particular TF binding motif when compared with the excluded variants. We ran these analyses for all the strong signals, but also strong signals stratified by ER status. Also, we subset this analysis to the variants located at regulatory regions in an ER-positive cell line (MCF-7 marked by H3K4me1, ENCODE id: ENCFF674BKS) and evaluated whether the ER-positive CCVs overlap any of the motifs more frequently that the excluded variants. We also evaluated the change in total binding affinity caused by the ER-positive CCCR alternative allele for all but one (2:217955891:T:<CN0>:0) of the ER-positive CCVs (*MatrixRider*^*87*^).

Subsequently, we evaluated whether the MCF-7 regions demarked by H3K4me1 (ENCODE id: ENCFF674BKS), and overlapped by ER-positive CCVs, were enriched in known TFBS motifs. We first subset the ENCODE bed file ENCFF674BKS to identify MCF-7 H3K4me1 peaks overlapped by the ER-positive CCVs (N = 107), as well as peaks only overlapped by excluded variants (N = 11,099), using BEDTools ^88^. We created fasta format sequences using genomic coordinate data from the intersected bed files. In order to create a control sequence set, we used the script included with the MEME Suite (*fasta-shuffle-letters*) to created 10 shuffled copies of each sequence overlapped by ER-positive CCVs (N = 1,070). We then used AME ^89^ to interrogate whether the 107 MCF-7 H3K4me1 genomic regions overlapped by ER-positive CCVs were enriched in know TFBS consensus motifs when compared to the shuffled control sequences, or to the MCF-7 H3K4me1 genomic regions overlapped only by excluded variants. We used the command line version of AME (version 4.12.0) selecting as scoring method the total number of positions in the sequence whose motif score p-value is less than 10^−3^, and using a one-tailed Fisher’s Exact test as the association test.

### PAINTOR analysis

To further refine the set of CCVs, we performed empirical Bayes fine-mapping using PAINTOR to integrate marginal genetic association summary statistics, linkage disequilibrium patterns, and biological features ^31,90^. PAINTOR places higher prior probability for causality on variants that fall in particular genomic features and learns the priors for different genomic features from the aggregate association results across all of the regions being mapped. PAINTOR does not assume a fixed number of causal variants in each region, although it implicitly penalizes non-parsimonious causal models. We applied PAINTOR separately to association results for overall breast cancer (in 85 regions determined to have at least one ER-neutral association or ER-positive and ER-negative association), ER-positive breast cancer (in 48 regions determined to have at least one ER-positive-specific association), and ER-negative breast cancer (in 22 regions determined to have at least one ER-negative-specific association). To avoid artifacts due to mis-matches between the LD in study samples and the LD matrix supplied to PAINTOR, we used association logistic regression summary statistics from OncoArray data only and estimated the LD structure in the OncoArray sample. For each endpoint we fit four models with increasing numbers of genomic features selected from the stepwise enrichment analyses described above: Model 0 (with no genomic features—assumes each variant is equally likely to be causal a priori), Model 1 (with those genomic features selected with stopping rule p<0.001); Model 2 (with those genomic features selected with stopping rule p<0.01); and Model 3 (with those genomic features selected with stopping rule p<0.05).

We used the Bayesian Information Criterion (BIC) to choose the best-fitting model for each outcome. As PAINTOR estimates the marginal log likelihood of the observed Z scores using Gibbs sampling, we used a shrunk mean BIC across multiple Gibbs chains to account for the stochasticity in the log-likelihood estimates. We ran PAINTOR four times to generate four independent Gibbs chains estimated the BIC difference between model *i* and model *j* as 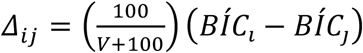. This assumes a N(0,100) prior on the difference, or roughly a 16% chance that model *i* would be decisively better than model *j* (i.e. |*BIC*_*i*_-*BIC*_*j*_|>10). We then proceeded to choose the best-fitting model in a stepwise fashion: starting with a model with no annotations, we selected a model with more annotations in favor a model with fewer if the larger model was a considerably better fit—i.e. Δ_*ij*_> 2. Model 1 was the best fit according to this process for overall and ER-positive breast cancer; Model 0 was the best fit for ER-negative breast cancer.

Differences between the PAINTOR and CCV outputs may be due to several factors. By considering functional enrichment and joint LD among all SNPs, PAINTOR may refine the set of likely causal variants; rather than imposing a hard threshold, PAINTOR allows for a gradient of evidence supporting causality; and the two sets of calculations are based on different summary statistics, CCV analyses used both iCOGS and OncoArray genotypes, while PAINTOR used only OncoArray data (**Figure 1**, Methods).

### Variant annotation

Variants genome coordinates were converted to assembly GRCh38 with liftOver and uploaded to Variant Effect Predictor ^91^ to determine their effect on genes, transcripts, and protein sequence. The commercial software Alamut^®^ Batch v1.6 batch was also used to annotate coding and splicing variants. PolyPhen-2 ^92^, SIFT ^93^, MAPP ^94^ were used to predict the consequence of missense coding variants. MaxEntScan ^95^, Splice-Site Finder, and Human Splicing Finder ^96^ were used to predict splicing effects.

### INQUISIT analysis

#### Logic underlying INQUISIT predictions

Briefly, genes were considered to potential targets of risk variants when variants affect: (1) distal regulation, (2) proximal genomic information, including chromatin interactions from capture Hi-C experiments performed in a panel of six breast cell lines ^97^, chromatin interaction analysis by paired-end tag sequencing (ChIA-PET; ^98^) and genome-wide chromosome conformation capture from HMECs (Hi-C, (Rao et al., 2014)). We used computational enhancer–promoter correlations (PreSTIGE ^99^, IM-PET (He et al., 2014), FANTOM5 ^100^ and super-enhancers ^28^), results for breast tissue-specific expression variants (eVariants) from multiple independent studies (TCGA, METABRIC, NHS, Methods), allele-specific imbalance in gene expression ^101^, transcription factor and histone modification chromatin immunoprecipitation followed by sequencing (ChIP-Seq) from the ENCODE and Roadmap Epigenomics Projects together with the genomic features found to be significantly enriched as described above, gene expression RNA-seq from several breast cancer lines and normal samples and topologically associated domain (TAD) boundaries from T47D cells (ENCODE, ^102^, Methods and Key Resources Table)). To assess the impact of intragenic variants, we evaluated their potential to alter splicing using Alamut^®^ Batch to identify new and cryptic donors and acceptors, and several tools to predict effects of coding sequence changes (Methods). The output from each tool was converted to a binary measure to indicate deleterious or tolerated predictions.

#### Scoring hierarchy

Each target gene prediction category (distal, promoter or coding) was scored according to different criteria. Genes predicted to be distally-regulated targets of CCVs were awarded points based on physical links (eg CHi-C), computational prediction methods, allele-specific expression, or eVariant associations. All CCV and HPPVs were considered as potentially involved in distal regulation. Intersection of a putative distal enhancer with genomic features found to be significantly enriched (see ‘**Genomic features enrichment’** for details) were further upweighted. CCVs and HPPVs in gene proximal regulatory regions were intersected with histone ChIP-Seq peaks characteristic of promoters and assigned to the overlapping transcription start sites (defined as −1.0 kb - +0.1 kb). Further points were awarded to such genes if there was evidence for eVariant association or allele-specific expression, while a lack of expression resulted in down-weighting as potential targets. Potential coding changes including missense, nonsense and predicted splicing alterations resulted in addition of one point to the encoded gene for each type of change, while lack of expression reduced the score. We added an additional point for predicted target genes that were also breast cancer drivers. For each category, scores ranged from 0-7 (distal); 0-3 (promoter) or 0-2 (coding). We converted these scores into ‘confidence levels’: Level 1 (highest confidence) when distal score > 4, promoter score >= 3 or coding score > 1; Level 2 when distal score <= 4 and >=1, promoter score = 1 or = 2, coding score = 1; and Level 3 when distal score < 1 and > 0, promoter score < 1 and > 0, and coding < 1 and > 0. For genes with multiple scores (for example, predicted as targets from multiple independent risk signals or predicted to be impacted in several categories), we recorded the highest score. Driver and transcription factor gene enrichment analysis was carried out using INQUISIT scores prior to adding a point for driver gene status. Modifications to the pipeline since original publication ^2^ include:

- TAD boundary definitions from ENCODE T47D Hi-C analysis. Previously, we used regions from Rao, Cell 2013;
- eQTL: Addition of NHS normal and tumor samples
- allele-specific imbalance using TCGA and GTEx RNA-seq data ^101^
- Capture Hi-C data from six breast cell lines (Beesley et al.)
- Additional biofeatures derived from global enrichment in this study.

#### Multi-signal targets

To test if more genes were targeted by multiple signals than expected by chance, we modelled the number of signals per gene by negative binomial regression (R function *glm.nb,* package MASS) and Poisson regression (R function *glm,* package stats) with ChIA-PET interactions as a covariate and adjusted by fine mapping region. Likelihood ratio tests were used to compare goodness of fit. Rootograms were created using the R function *rootogram* (package vcd).

### Pathway analysis

The pathway gene set database, dated 1 September 2018 was used ^103^ (http://download.baderlab.org/EM_Genesets/current_release/Human/symbol/). This database contains pathways from Reactome ^104^, NCI Pathway Interaction Database ^105^, GO (Gene Ontology) ^106^, HumanCyc ^107^, MSigdb ^108^, NetPath ^109^, and Panther ^110^. All duplicated pathways, defined in two or more databases, were included. To provide more biologically meaningful results, only pathways that contained ≤ 200 genes were used.

We interrogated the pathway annotation sets with the list of high-confidence (Level 1) INQUISIT gene list. The significance of over-representation of the INQUISIT genes within each pathway was assessed with a hypergeometric test, ^111^, using the R function *phyper* as follows:

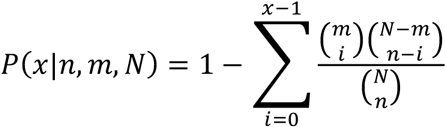

where x is the number of Level 1 genes that overlap with any of the genes in the pathway, n is the number of genes in the pathway, m is the number of Level1 genes that overlap with any of the genes in the pathway data set (m_strong GO_ = 135, m_ER-positive GO_ = 48, m_ER-negative GO_ = 26, m_ER-neutral GO_ = 73; m_strong Pathways_ = 114, m_ER-positive Pathways_ = 36, m_ER-negative Pathways_ = 21, m_ER-neutral Pathways_ = 67), and N is the number of genes in the pathway data set (N_Genes GO_ = 14,252, N_Genes Pathways_ = 10,915). We only included pathways that overlapped with at least two Level 1 genes. We used the Benjamini-Hochberg false discovery rate (FDR) ^62^ at 5% level.

## ACKNOWLEDGMENTS

We thank all the individuals who took part in these studies and all the researchers, clinicians, technicians and administrative staff who have enabled this work to be carried out. This work was supported by the European Union’s Horizon 2020 research and innovation programme under the Marie Sklodowska-Curie grant agreement No 656144. Genotyping of the OncoArray was principally funded from three sources: the PERSPECTIVE project, funded by the Government of Canada through Genome Canada and the Canadian Institutes of Health Research, the ‘Ministère de l’Économie, de la Science et de l’Innovation du Québec’ through Genome Québec, and the Quebec Breast Cancer Foundation; the NCI Genetic Associations and Mechanisms in Oncology (GAME-ON) initiative and Discovery, Biology and Risk of Inherited Variants in Breast Cancer (DRIVE) project (NIH Grants U19 CA148065 and X01HG007492); and Cancer Research UK (C1287/A10118 and C1287/A16563). BCAC is funded by Cancer Research UK (C1287/A16563), by the European Community’s Seventh Framework Programme under grant agreement 223175 (HEALTH-F2-2009-223175) (COGS) and by the European Union’s Horizon 2020 Research and Innovation Programme under grant agreements 633784 (B-CAST) and 634935 (BRIDGES). Genotyping of the iCOGS array was funded by the European Union (HEALTH-F2-2009-223175), Cancer Research UK (C1287/A10710), the Canadian Institutes of Health Research for the ‘CIHR Team in Familial Risks of Breast Cancer’ program, and the Ministry of Economic Development, Innovation and Export Trade of Quebec, grant PSR-SIIRI-701. Combining of the GWAS data was supported in part by The National Institute of Health (NIH) Cancer Post-Cancer GWAS initiative grant U19 CA 148065 (DRIVE, part of the GAME-ON initiative). For a full description of funding and acknowledgments, see Supplementary Note.

## AUTHOR CONTRIBUTIONS

Conceptualization: L.Fa., H.As., J.Be., D.R.B., J.Al., S.Ka., K.A.P., K.Mi., P.So., A.Le., M.Gh., P.D.P.P., J.C.C., M.G.C., M.K.S., R.L.M., V.N.K., J.D.E., S.L.E., A.C.A., G.C.T., J.Si., D.F.E., P.K., A.M.D. Methodology: L.Fa., H.As., J.Be., D.R.B., J.Al., J.D.E., S.L.E., A.C.A., G.C.T., J.Si., D.F.E., P.K., A.M.D. Software: J.Be., J.P.T., M.L. Formal analysis: L.Fa., H.As., J.Be., D.R.B., J.Al., S.Ka., C.Tu., M.Mor., X.J. Resources: S.A., K.A., M.R.A., I.L.A., H.A.C., N.N.A., A.A., V.A., K.J.A., B.K.A., B.A., P.L.A., J.Az., J.Ba., R.B.B., D.B., A.B.F., J.Ben., M.B., K.B., A.M.B., C.B., W.B., N.V.B., S.E.B., B.Bo., A.B., H.Bra., H.Bre., I.B., I.W.B., A.B.W., T.B., B.Bu., S.S.B., Q.C., T.C., M.A.C., N.J.C., I.C., F.C., J.S.C., B.D.C., J.E.C., J.C., H.C., W.K.C., K.B.M., C.L.C., J.M.C., S.C., F.J.C., A.C., S.S.C., C.C., K.C., M.B.D., M.D.H., P.D., O.D., Y.C.D., G.S.D., S.M.D., T.D., I.D.S., A.D., S.D., M.Dum., M.Dur., L.D., M.Dw., D.M.E., C.E., M.E., D.G.E., P.A.F., U.F., O.F., G.F., H.F., L.Fo., W.D.F., E.F., L.Fr., D.F., M.Ga., M.G.D., G.Ga., P.A.G., S.M.G., J.Ga., J.A.G., M.M.G., V.G., G.G.G., G.Gl., A.K.G., M.S.G., D.E.G., A.G.N., M.H.G., M.Gr., J.Gr., A.G., P.G., E.H., C.A.H., N.H., P.Ha., U.H., P.A.H., J.M.H., M.H., W.H., C.S.H., B.A.M., J.H., P.Hi., F.B.L., A.H., M.J.H., J.L.H., A.Ho., G.H., P.J.H., E.N.I., C.I., M.I., A.Jag., M.J., A.Jak., P.J., R.J., R.C.J., E.M.J., N.J., M.E.J., A.Juk., A.Jun., R.Ka., D.K., B.Pes., R.Ke., M.J.K., E.K., J.I.K., J.K., C.M.K., Y.K., I.K., V.K., S.Ko., K.K.S., T.K., A.K., K.K., Y.L., D.L., E.L., G.L., J.Le., F.L., A.Li., W.L., J.Lo., A.Lo., J.T.L., J.Lu., R.J.M., T.M., E.M., A.Ma., M.Ma., S.Man., S.Mag., M.E.M., K.Ma., D.M., R.M., L.M., C.M., N.Me., A.Me., P.M., A.Mi., N.Mi., M.Mo., F.M., A.M.M., V.M.M., T.A., S.A.N., R.N., K.L.N., N.Z.N., H.N., P.N., F.C.N., L.N.Z., A.N., K.O., E.O., O.I.O., H.O., N.O., A.O., V.S.P., J.Pa., S.K.P., T.W.P.S., M.T.P., J.Pau., I.S.P., B.Pei., B.Y.K., P.P., J.Pe., D.P.K., K.Pr., R.P., N.P., D.P., M.A.P., K.Py., P.R., S.J.R., J.R., R.R.M., G.R., H.A.R., M.R., A.R., C.M.R., E.S., E.S.H., D.P.S., M.Sa., C.Sa., E.J.S., M.T.S., D.F.S., R.K.S., A.S., M.J.S., B.S., P.Sc., C.Sc., R.J.S., L.S., C.M.D., M.Sh., P.Sh., C.Y.S., X.S., C.F.S., T.P.S., S.S., M.C.S., J.J.S., A.B.S., J.St., D.S.L., C.Su., A.J.S., R.M.T., Y.Y.T., W.J.T., J.A.T., M.R.T., M.Te., S.H., M.B.T., A.T., M.Th., D.L.T., M.G.T., M.Ti., A.E.T., R.A.E., I.T., D.T., G.T.M., M.A.T., N.T., M.Tz., H.U.U., C.M.V., C.J.A., L.E.K., E.J.R., A.Ve., A.Vi., J.V., M.J.V., Q.W., B.W., C.R.W., J.N.W., C.W., H.W., R.W., A.W., A.H.W., D.Y., Y.Z., W.Z. Data management and curation: K.Mi., J.D., M.K.B., Q.W., R.Ke., J.C.C. and M.K.S. Writing original draft: L.Fa., H.As., J.Be., G.C.T., D.F.E., P.K., A.M.D. Writing review and editing: D.R.B., J.Al., P.So., A.Le., V.N.K., J.D.E., S.L.E., A.C.A., J.Si. Visualization: L.Fa., H.As., J.Be., C.Tu. Supervision: A.C.A., G.C.T., J.Si., D.F.E., P.K., A.M.D. Funding acquisition: L.Fa., P.D.P.P., J.C.C., M.G.C., M.K.S., R.L.M., V.N.K., J.D.E., S.L.E., A.C.A., G.C.T., J.Si., D.F.E., P.K., A.M.D. All authors read and approved the final version of the manuscript.

## MATERIALS & CORRESPONDENCE

Further information and requests for resources should be directed to and will be fulfilled by Manjeet Bolla

## REFERENCES

[1] Milne, R.L. et al. Identification of ten variants associated with risk of estrogen-receptor-negative breast cancer. Nat Genet 49, 1767–1778 (2017).

[2] Michailidou, K. et al. Association analysis identifies 65 new breast cancer risk loci. Nature 551, 92–94 (2017).

[3] Ghoussaini, M. et al. Evidence that breast cancer risk at the 2q35 locus is mediated through IGFBP5 regulation. Nat Commun 4, 4999 (2014).

[4] Wyszynski, A. et al. An intergenic risk locus containing an enhancer deletion in 2q35 modulates breast cancer risk by deregulating IGFBP5 expression. Hum Mol Genet 25, 3863–3876 (2016).

[5] Guo, X. et al. Fine-scale mapping of the 4q24 locus identifies two independent loci associated with breast cancer risk. Cancer Epidemiol Biomarkers Prev 24, 1680–91 (2015).

[6] Glubb, D.M. et al. Fine-scale mapping of the 5q11.2 breast cancer locus reveals at least three independent risk variants regulating MAP3K1. Am J Hum Genet 96, 5–20 (2015).

[7] Dunning, A.M. et al. Breast cancer risk variants at 6q25 display different phenotype associations and regulate ESR1, RMND1 and CCDC170. Nat Genet 48, 374–86 (2016).

[8] Shi, J. et al. Fine-scale mapping of 8q24 locus identifies multiple independent risk variants for breast cancer. Int J Cancer 139, 1303–1317 (2016).

[9] Orr, N. et al. Fine-mapping identifies two additional breast cancer susceptibility loci at 9q31.2. Hum Mol Genet 24, 2966–84 (2015).

[10] Darabi, H. et al. Polymorphisms in a Putative Enhancer at the 10q21.2 Breast Cancer Risk Locus Regulate NRBF2 Expression. Am J Hum Genet 97, 22–34 (2015).

[11] Darabi, H. et al. Fine scale mapping of the 17q22 breast cancer locus using dense SNPs, genotyped within the Collaborative Oncological Gene-Environment Study (COGs). Sci Rep 6, 32512 (2016).

[12] Meyer, K.B. et al. Fine-scale mapping of the FGFR2 breast cancer risk locus: putative functional variants differentially bind FOXA1 and E2F1. Am J Hum Genet 93, 1046–60 (2013).

[13] Betts, J.A. et al. Long noncoding RNAs CUPID1 and CUPID2 mediate breast cancer risk at 11q13 by modulating the response to dna damage. Am J Hum Genet 101, 255–266 (2017).

[14] French, J.D. et al. Functional variants at the 11q13 risk locus for breast cancer regulate cyclin D1 expression through long-range enhancers. Am J Hum Genet 92, 489–503 (2013).

[15] Ghoussaini, M. et al. Evidence that the 5p12 variant rs10941679 Confers susceptibility to estrogen-receptor-positive breast cancer through FGF10 and MRPS30 regulation. Am J Hum Genet 99, 903–911 (2016).

[16] Horne, H.N. et al. Fine-mapping of the 1p11.2 Breast cancer susceptibility locus. PLoS One 11, e0160316 (2016).

[17] Zeng, C. et al. Identification of independent association signals and putative functional variants for breast cancer risk through fine-scale mapping of the 12p11 locus. Breast Cancer Res 18, 64 (2016).

[18] Lin, W.Y. et al. Identification and characterization of novel associations in the CASP8/ALS2CR12 region on chromosome 2 with breast cancer risk. Hum Mol Genet 24, 285–98 (2015).

[19] Bojesen, S.E. et al. Multiple independent variants at the TERT locus are associated with telomere length and risks of breast and ovarian cancer. Nat Genet 45, 371-84, 384e1-2 (2013).

[20] Lawrenson, K. et al. Functional mechanisms underlying pleiotropic risk alleles at the 19p13.1 breast-ovarian cancer susceptibility locus. Nat Commun 7, 12675 (2016).

[21] Amos, C.I. et al. The OncoArray Consortium: A Network for Understanding the Genetic Architecture of Common Cancers. Cancer Epidemiol Biomarkers Prev 26, 126–135 (2017).

[22] Michailidou, K. et al. Large-scale genotyping identifies 41 new loci associated with breast cancer risk. Nat Genet 45, 353-61, 361e1-2 (2013).

[23] Michailidou, K. et al. Genome-wide association analysis of more than 120,000 individuals identifies 15 new susceptibility loci for breast cancer. Nature Genetics 47, 373–U127 (2015).

[24] Udler, M.S., Tyrer, J. & Easton, D.F. Evaluating the power to discriminate between highly correlated SNPs in genetic association studies. Genet Epidemiol 34, 463–8 (2010).

[25] Mavaddat, N., Antoniou, A.C., Easton, D.F. & Garcia-Closas, M. Genetic susceptibility to breast cancer. Mol Oncol 4, 174–91 (2010).

[26] Lakhani, S.R. et al. Prediction of BRCA1 status in patients with breast cancer using estrogen receptor and basal phenotype. Clin Cancer Res 11, 5175–80 (2005).

[27] Taberlay, P.C., Statham, A.L., Kelly, T.K., Clark, S.J. & Jones, P.A. Reconfiguration of nucleosome-depleted regions at distal regulatory elements accompanies DNA methylation of enhancers and insulators in cancer. Genome Res 24, 1421–32 (2014).

[28] Hnisz, D. et al. Super-enhancers in the control of cell identity and disease. Cell 155, 934–47 (2013).

[29] Farh, K.K. et al. Genetic and epigenetic fine mapping of causal autoimmune disease variants. Nature 518, 337–43 (2015).

[30] Cowper-Sal lari, R. et al. Breast cancer risk-associated SNPs modulate the affinity of chromatin for FOXA1 and alter gene expression. Nat Genet 44, 1191–8 (2012).

[31] Kichaev, G. et al. Integrating functional data to prioritize causal variants in statistical fine-mapping studies. PLoS Genet 10, e1004722 (2014).

[32] Quiroz-Zarate, A. et al. Expression Quantitative Trait loci (QTL) in tumor adjacent normal breast tissue and breast tumor tissue. PLoS One 12, e0170181 (2017).

[33] Cancer Genome Atlas Research, N. et al. The Cancer Genome Atlas Pan-Cancer analysis project. Nat Genet 45, 1113–20 (2013).

[34] Curtis, C. et al. The genomic and transcriptomic architecture of 2,000 breast tumours reveals novel subgroups. Nature 486, 346–52 (2012).

[35] Ciriello, G. et al. Comprehensive Molecular Portraits of Invasive Lobular Breast Cancer. Cell 163, 506–19 (2015).

[36] Nik-Zainal, S. et al. Landscape of somatic mutations in 560 breast cancer whole-genome sequences. Nature 534, 47–54 (2016).

[37] Pereira, B. et al. The somatic mutation profiles of 2,433 breast cancers refines their genomic and transcriptomic landscapes. Nat Commun 7, 11479 (2016).

[38] Cancer Genome Atlas, N. Comprehensive molecular portraits of human breast tumours. Nature 490, 61–70 (2012).

[39] Bailey, M.H. et al. Comprehensive Characterization of Cancer Driver Genes and Mutations. Cell 173, 371-385 e18 (2018).

[40] Lambert, S.A. et al. The Human Transcription Factors. Cell 172, 650–665 (2018).

[41] Artero-Castro, A. et al. Disruption of the ribosomal P complex leads to stress-induced autophagy. Autophagy 11, 1499–519 (2015).

[42] Wang, X.Y. et al. Musashi1 modulates mammary progenitor cell expansion through proliferin-mediated activation of the Wnt and Notch pathways. Mol Cell Biol 28, 3589–99 (2008).

[43] Vijayan, D., Young, A., Teng, M.W.L. & Smyth, M.J. Targeting immunosuppressive adenosine in cancer. Nat Rev Cancer 17, 709–724 (2017).

[44] Takebe, N. et al. Targeting Notch, Hedgehog, and Wnt pathways in cancer stem cells: clinical update. Nat Rev Clin Oncol 12, 445–64 (2015).

[45] Thorpe, L.M., Yuzugullu, H. & Zhao, J.J. PI3K in cancer: divergent roles of isoforms, modes of activation and therapeutic targeting. Nat Rev Cancer 15, 7–24 (2015).

[46] Nusse, R. & Clevers, H. Wnt/beta-Catenin Signaling, Disease, and Emerging Therapeutic Modalities. Cell 169, 985–999 (2017).

[47] Massague, J. TGFbeta signalling in context. Nat Rev Mol Cell Biol 13, 616–30 (2012).

[48] Meeks, H.D. et al. BRCA2 polymorphic stop codon K3326X and the Risk of breast, prostate, and ovarian cancers. J Natl Cancer Inst 108(2016).

[49] CHEK2 Breast Cancer Case-Control Consortium. CHEK2*1100delC and susceptibility to breast cancer: a collaborative analysis involving 10,860 breast cancer cases and 9,065 controls from 10 studies. Am J Hum Genet 74, 1175–82 (2004).

[50] Schmidt, M.K. et al. Age- and Tumor Subtype-Specific Breast Cancer Risk Estimates for CHEK2*1100delC Carriers. J Clin Oncol 34, 2750–60 (2016).

[51] Kilpivaara, O. et al. CHEK2 variant I157T may be associated with increased breast cancer risk. Int J Cancer 111, 543–7 (2004).

[52] Muranen, T.A. et al. Patient survival and tumor characteristics associated with CHEK2:p.I157T - findings from the Breast Cancer Association Consortium. Breast Cancer Res 18, 98 (2016).

[53] Killedar, A. et al. A Common Cancer Risk-Associated Allele in the hTERT Locus Encodes a Dominant Negative Inhibitor of Telomerase. PLoS Genet 11, e1005286 (2015).

[54] De Blasio, A. et al. Unusual roles of caspase-8 in triple-negative breast cancer cell line MDA-MB-231. Int J Oncol 48, 2339–48 (2016).

[55] Haupt, S. et al. Targeting Mdmx to treat breast cancers with wild-type p53. Cell Death Dis 6, e1821 (2015).

[56] Pandya, P.H., Murray, M.E., Pollok, K.E. & Renbarger, J.L. The Immune System in Cancer Pathogenesis: Potential Therapeutic Approaches. J Immunol Res 2016, 4273943 (2016).

[57] Gionet, N., Jansson, D., Mader, S. & Pratt, M.A.NF-kappaB and estrogen receptor alpha interactions: Differential function in estrogen receptor-negative and -positive hormone-independent breast cancer cells. J Cell Biochem 107, 448–59 (2009).

[58] Fleischer, T. et al. DNA methylation at enhancers identifies distinct breast cancer lineages. Nat Commun 8, 1379 (2017).

[59] Couch, F.J. et al. Genome-wide association study in BRCA1 mutation carriers identifies novel loci associated with breast and ovarian cancer risk. PLoS Genet 9, e1003212 (2013).

[60] Gaudet, M.M. et al. Identification of a BRCA2-specific modifier locus at 6p24 related to breast cancer risk. PLoS Genet 9, e1003173 (2013).

[61] Marchini, J., Howie, B., Myers, S., McVean, G. & Donnelly, P. A new multipoint method for genome-wide association studies by imputation of genotypes. Nat Genet 39, 906–13 (2007).

[62] Benjamini, Y. & Hochberg, Y. Controlling the False Discovery Rate - a Practical and Powerful Approach to Multiple Testing. Journal of the Royal Statistical Society Series B-Methodological 57, 289–300 (1995).

[63] Liu, J.Z. et al. Meta-analysis and imputation refines the association of 15q25 with smoking quantity. Nat Genet 42, 436–40 (2010).

[64] Antoniou, A.C. et al. RAD51 135G-->C modifies breast cancer risk among BRCA2 mutation carriers: results from a combined analysis of 19 studies. Am J Hum Genet 81, 1186–200 (2007).

[65] Barnes, D.R. et al. Evaluation of association methods for analysing modifiers of disease risk in carriers of high-risk mutations. Genet Epidemiol 36, 274–91 (2012).

[66] Willer, C.J., Li, Y. & Abecasis, G.R. METAL: fast and efficient meta-analysis of genomewide association scans. Bioinformatics 26, 2190–1 (2010).

[67] Hunter, D.J. et al. A genome-wide association study identifies alleles in FGFR2 associated with risk of sporadic postmenopausal breast cancer. Nat Genet 39, 870–4 (2007).

[68] Baran, Y. et al. Fast and accurate inference of local ancestry in Latino populations. Bioinformatics 28, 1359–67 (2012).

[69] Howie, B., Fuchsberger, C., Stephens, M., Marchini, J. & Abecasis, G.R. Fast and accurate genotype imputation in genome-wide association studies through pre-phasing. Nat Genet 44, 955–9 (2012).

[70] Genomes Project, C. et al. An integrated map of genetic variation from 1,092 human genomes. Nature 491, 56–65 (2012).

[71] Li, B. & Dewey, C.N. RSEM: accurate transcript quantification from RNA-Seq data with or without a reference genome. BMC Bioinformatics 12, 323 (2011).

[72] Mermel, C.H. et al. GISTIC2.0 facilitates sensitive and confident localization of the targets of focal somatic copy-number alteration in human cancers. Genome Biol 12, R41 (2011).

[73] Li, Q. et al. Integrative eQTL-based analyses reveal the biology of breast cancer risk loci. Cell 152, 633–41 (2013).

[74] Shabalin, A.A. Matrix eQTL: ultra fast eQTL analysis via large matrix operations. Bioinformatics 28, 1353–8 (2012).

[75] ENCODE Project Consortium. An integrated encyclopedia of DNA elements in the human genome. Nature 489, 57–74 (2012).

[76] Sloan, C.A. et al. ENCODE data at the ENCODE portal. Nucleic Acids Res 44, D726–32 (2016).

[77] Roadmap Epigenomics, C. et al. Integrative analysis of 111 reference human epigenomes. Nature 518, 317–30 (2015).

[78] Stunnenberg, H.G., International Human Epigenome, C. & Hirst, M. The International Human Epigenome Consortium: A Blueprint for Scientific Collaboration and Discovery. Cell 167, 1897 (2016).

[79] Pellacani, D. et al. Analysis of Normal Human Mammary Epigenomes Reveals Cell-Specific Active Enhancer States and Associated Transcription Factor Networks. Cell Rep 17, 2060–2074 (2016).

[80] Cheneby, J., Gheorghe, M., Artufel, M., Mathelier, A. & Ballester, B. ReMap 2018: an updated atlas of regulatory regions from an integrative analysis of DNA-binding ChIP-seq experiments. Nucleic Acids Res 46, D267–D275 (2018).

[81] Pruitt, K.D. et al. RefSeq: an update on mammalian reference sequences. Nucleic Acids Res 42, D756–63 (2014).

[82] Harrow, J. et al. GENCODE: the reference human genome annotation for The ENCODE Project. Genome Res 22, 1760–74 (2012).

[83] Wang, J. et al. Sequence features and chromatin structure around the genomic regions bound by 119 human transcription factors. Genome Res 22, 1798–812 (2012).

[84] Mathelier, A. et al. JASPAR 2016: a major expansion and update of the open-access database of transcription factor binding profiles. Nucleic Acids Res 44, D110–5 (2016).

[85] Tan, G. & Lenhard, B. TFBSTools: an R/bioconductor package for transcription factor binding site analysis. Bioinformatics 32, 1555–6 (2016).

[86] Grant, C.E., Bailey, T.L. & Noble, W.S. FIMO: scanning for occurrences of a given motif. Bioinformatics 27, 1017–8 (2011).

[87] Grassi, E., Zapparoli, E., Molineris, I. & Provero, P. Total Binding Affinity Profiles of Regulatory Regions Predict Transcription Factor Binding and Gene Expression in Human Cells. PLoS One 10, e0143627 (2015).

[88] Quinlan, A.R. & Hall, I.M. BEDTools: a flexible suite of utilities for comparing genomic features. Bioinformatics 26, 841–2 (2010).

[89] McLeay, R.C. & Bailey, T.L. Motif Enrichment Analysis: a unified framework and an evaluation on ChIP data. BMC Bioinformatics 11, 165 (2010).

[90] Kichaev, G. et al. Improved methods for multi-trait fine mapping of pleiotropic risk loci. Bioinformatics 33, 248–255 (2017).

[91] McLaren, W. et al. The Ensembl Variant Effect Predictor. Genome Biol 17, 122 (2016).

[92] Adzhubei, I.A. et al. A method and server for predicting damaging missense mutations. Nat Methods 7, 248–9 (2010).

[93] Kumar, P., Henikoff, S. & Ng, P.C. Predicting the effects of coding non-synonymous variants on protein function using the SIFT algorithm. Nat Protoc 4, 1073–81 (2009).

[94] Stone, E.A. & Sidow, A. Physicochemical constraint violation by missense substitutions mediates impairment of protein function and disease severity. Genome Res 15, 978–86 (2005).

[95] Yeo, G. & Burge, C.B. Maximum entropy modeling of short sequence motifs with applications to RNA splicing signals. J Comput Biol 11, 377–94 (2004).

[96] Desmet, F.O. et al. Human Splicing Finder: an online bioinformatics tool to predict splicing signals. Nucleic Acids Res 37, e67 (2009).

[97] Beesley, J. et al. Chromatin interactome mapping at 139 independent breast cancer risk signals. Submitted.

[98] Fullwood, M.J. et al. An oestrogen-receptor-alpha-bound human chromatin interactome. Nature 462, 58–64 (2009).

[99] Corradin, O. et al. Combinatorial effects of multiple enhancer variants in linkage disequilibrium dictate levels of gene expression to confer susceptibility to common traits. Genome Res 24, 1–13 (2014).

[100] Andersson, R. et al. An atlas of active enhancers across human cell types and tissues. Nature 507, 455–461 (2014).

[101] Moradi Marjaneh, M. et al. High-throughput allelic expression imbalance analyses identify 14 candidate breast cancer risk genes. Submitted.

[102] Dixon, J.R. et al. Integrative detection and analysis of structural variation in cancer genomes. Nat Genet 50, 1388–1398 (2018).

[103] Merico, D., Isserlin, R. & Bader, G.D. Visualizing gene-set enrichment results using the Cytoscape plug-in enrichment map. Methods Mol Biol 781, 257–77 (2011).

[104] Vastrik, I. et al. Reactome: a knowledge base of biologic pathways and processes. Genome Biol 8, R39 (2007).

[105] Schaefer, C.F. et al. PID: the Pathway Interaction Database. Nucleic Acids Res 37, D674-9 (2009).

[106] Ashburner, M. et al. Gene ontology: tool for the unification of biology. The Gene Ontology Consortium. Nat Genet 25, 25–9 (2000).

[107] Romero, P. et al. Computational prediction of human metabolic pathways from the complete human genome. Genome Biol 6, R2 (2005).

[108] Subramanian, A. et al. Gene set enrichment analysis: a knowledge-based approach for interpreting genome-wide expression profiles. Proc Natl Acad Sci U S A 102, 15545–50 (2005).

[109] Kandasamy, K. et al. NetPath: a public resource of curated signal transduction pathways. Genome Biol 11, R3 (2010).

[110] Thomas, P.D. et al. PANTHER: a library of protein families and subfamilies indexed by function. Genome Res 13, 2129–41 (2003).

[111] Herwig, R., Hardt, C., Lienhard, M. & Kamburov, A. Analyzing and interpreting genome data at the network level with ConsensusPathDB. Nat Protoc 11, 1889–907 (2016).

