## Supplementary Information for "Fine-mapping of 150 breast cancer risk regions identifies 178 high confidence target genes"

**Supplementary Excel Table guide, supplied as individual files**

**Table S1.** Breast cancer risk regions identified through genome-wide association studies.

(a) Fine-mapping regions. (b) Imputation quality metrics across fine-mapping regions.

**Table S2**. Breast cancer risk signals and credible candidate variants (CCVs).

(a) Multinomial Regression models. (b) Strong signals (BCAC and CIMBA) multinomial regression models.

**Table S3.** Bio-features enrichment.

**Table S4.** Consensus transcription factor binding motif enrichment.

(a) Transcription Factor consensus binding motif enrichment analysis. (b) Transcription Factor enrichment at MCF-7 H3K4me1 regions . (c) ER-positive CCVs overlap with transcription factor binding motifs significantly enriched

**Table S5.** Coding, splicing CCVs and overlap of CCVs with variant drivers of local gene expression.

(a) CCVs collocating with eQTL variants in normal breast tissue. (b) CCVs collocating with eQTL variants in breast tumor tissue. (c) CCVs coding annotation. (d) CCVs predicted to affect splicing

**Table S6.** 178 Level 1 predicted target genes.

**Table S7.** INQUISIT results for coding/splicing variants.

**Table S8.** INQUISIT results for promoter variants.

**Table S9.** INQUISIT results for distal variants.

**Table S10.** Pathways significantly enriched in CCV and high posterior probability predicted target genes.

### SUPPLEMENTARY FIGURES

### Figure S1. Bio-features enrichment.

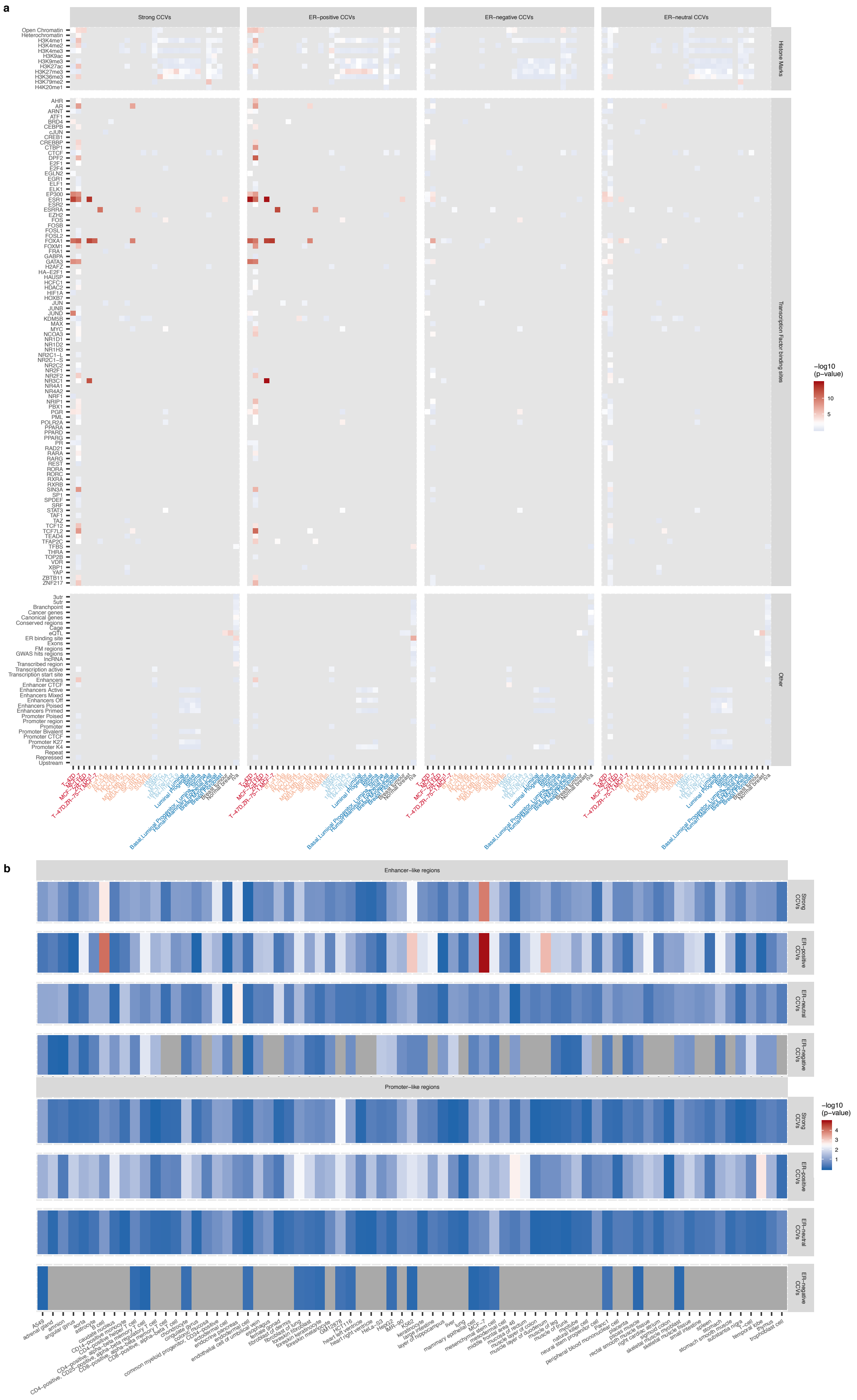

(a) Intersection between CCVs and known bio-features. (b) ENCODE enhancer-like and promoter-like enrichment. ENCODE enhancer-like regions, top, ENCODE promoter like tissues, bottom. Each bar shows the overlap p-value for each subset of CCVs (Strong, ER-positive, ER-negative and ER-neutral) with regulatory regions defined by ENCODE at 73 tissues, primary cells, immortalized cell line, and in vitro differentiated cells (from most significant, dark red, to less significant, blue; grey bars indicate regions where there is <5 CCVs overlapping the region).

**Figure S2. Correlation between CCVs overlapping significantly enriched bio-features.**

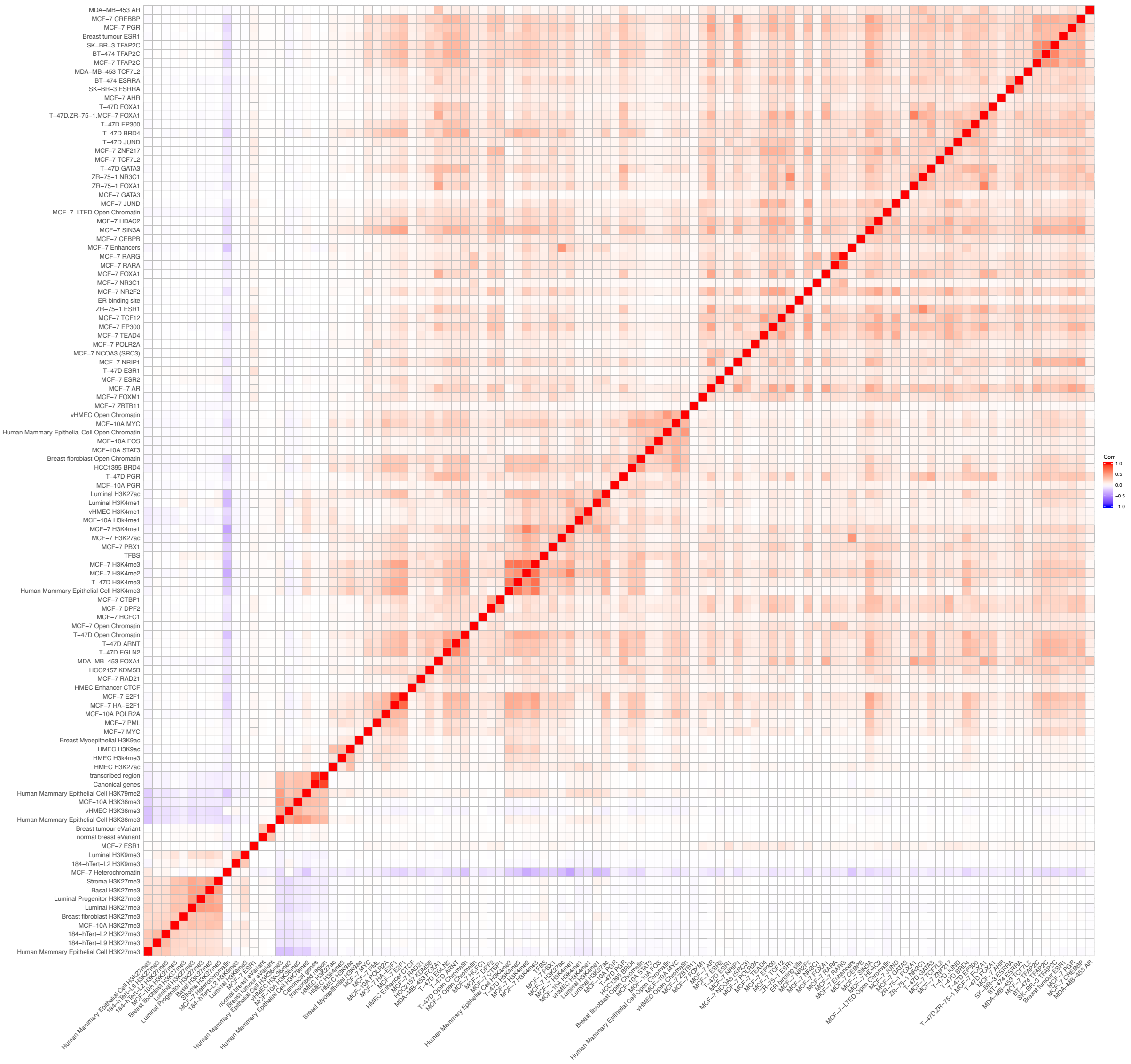

**Figure S3. Bayesian fine mapping.**

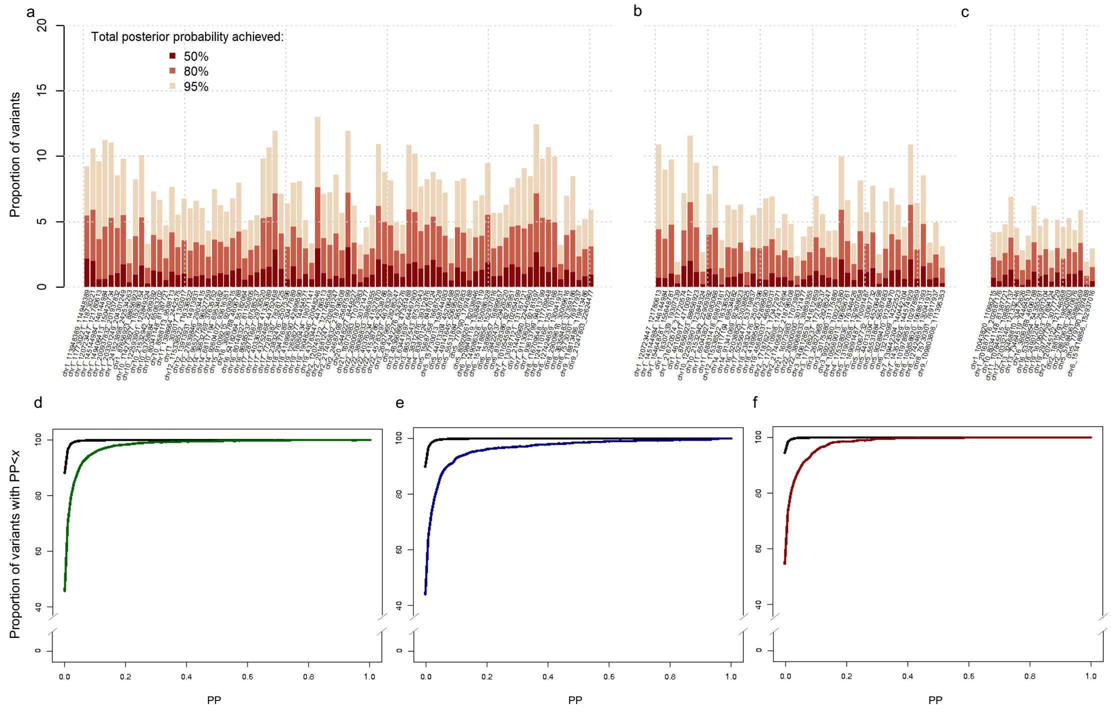


Number of Variants per total PP from (a) ER-all model (b) ER-positive PAINTOR model (c) ER-negative PAINTOR model. Cumulative distributions of PP for variants in strong signals for overall breast cancer (d, green), strong signals for ER-positive breast cancer (e, blue), and strong signals for ER-negative breast cancer (f, red), compared to cumulative distributions of variants outside of these signals (black).

**Figure S4. Predicted target genes over-dispersion analysis and enrichment analysis.**

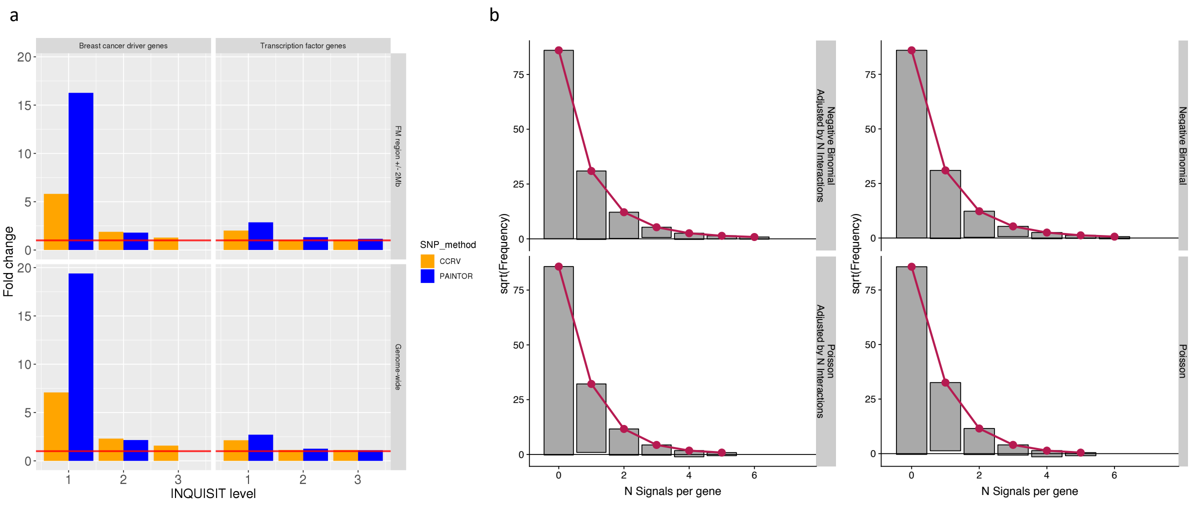


(a) Predicted target genes are enriched in known breast cancer driver genes and TFs. (b) Hanging rootograms for the negative binomial model (glm.ng), and the Poisson model (glm.pois). The red line represents the expected counts given the model. The bars denote the observed counts. X-axis shows the count bin. Y-axis shows the square root of the observed or expected count

**BCAC FUNDING AND ACKNOWLEDGMENTS**

**Funding**

BCAC is funded by Cancer Research UK [C1287/A16563, C1287/A10118], the European Union's Horizon 2020 Research and Innovation Programme (grant numbers 634935 and 633784 for BRIDGES and B-CAST respectively), and by the European Community´s Seventh Framework Programme under grant agreement number 223175 (grant number HEALTH-F2-2009-223175) (COGS). The EU Horizon 2020 Research and Innovation Programme funding source had no role in study design, data collection, data analysis, data interpretation or writing of the report.

Genotyping of the OncoArray was funded by the NIH Grant U19 CA148065, and Cancer UK Grant C1287/A16563 and the PERSPECTIVE project supported by the Government of Canada through Genome Canada and the Canadian Institutes of Health Research (grant GPH-129344) and, the Ministère de l'Économie, Science et Innovation du Québec through Genome Québec and the PSRSIIRI-701 grant, and the Quebec Breast Cancer Foundation. Funding for the iCOGS infrastructure came from: the European Community's Seventh Framework Programme under grant agreement n° 223175 (HEALTH-F2-2009-223175) (COGS), Cancer Research UK (C1287/A10118, C1287/A10710, C12292/A11174, C1281/A12014, C5047/A8384, C5047/A15007, C5047/A10692, C8197/A16565), the National Institutes of Health (CA128978) and Post-Cancer GWAS initiative (1U19 CA148537, 1U19 CA148065 and 1U19 CA148112 - the GAME-ON initiative), the Department of Defence (W81XWH-10-1-0341), the Canadian Institutes of Health Research (CIHR) for the CIHR Team in Familial Risks of Breast Cancer, and Komen Foundation for the Cure, the Breast Cancer Research Foundation, and the Ovarian Cancer Research Fund. The DRIVE Consortium was funded by U19 CA148065.

The Australian Breast Cancer Family Study (ABCFS) was supported by grant UM1 CA164920 from the National Cancer Institute (USA). The content of this manuscript does not necessarily reflect the views or policies of the National Cancer Institute or any of the collaborating centers in the Breast Cancer Family Registry (BCFR), nor does mention of trade names, commercial products, or organizations imply endorsement by the USA Government or the BCFR. The ABCFS was also supported by the National Health and Medical Research Council of Australia, the New South Wales Cancer Council, the Victorian Health Promotion Foundation (Australia) and the Victorian Breast Cancer Research Consortium. J.L.H. is a National Health and Medical Research Council (NHMRC) Senior Principal Research Fellow. M.C.S. is a NHMRC Senior Research Fellow. The ABCS study was supported by the Dutch Cancer Society [grants NKI 2007-3839; 2009 4363]. The Australian Breast Cancer Tissue Bank (ABCTB) is generously supported by the National Health and Medical Research Council of Australia, The Cancer Institute NSW and the National Breast Cancer Foundation. The ACP study is funded by the Breast Cancer Research Trust, UK. The AHS study is supported by the intramural research program of the National Institutes of Health, the National Cancer Institute (grant number Z01-CP010119), and the National Institute of Environmental Health Sciences (grant number Z01-ES049030). The work of the BBCC was partly funded by ELAN-Fond of the University Hospital of Erlangen. The BBCS is funded by Cancer Research UK and Breast Cancer Now and acknowledges NHS funding to the NIHR Biomedical Research Centre, and the National Cancer Research Network (NCRN). The BCEES was funded by the National Health and Medical Research Council, Australia and the Cancer Council Western Australia and acknowledges funding from the National Breast Cancer Foundation (JS). For the BCFR-NY, BCFR-PA, BCFR-UT this work was supported by grant UM1 CA164920 from the National Cancer Institute. The content of this manuscript does not necessarily reflect the views or policies of the National Cancer Institute or any of the collaborating centers in the Breast Cancer Family Registry (BCFR), nor does mention of trade names, commercial products, or organizations imply endorsement by the US Government or the BCFR. For BIGGS, ES is supported by NIHR Comprehensive Biomedical Research Centre, Guy's & St. Thomas' NHS Foundation Trust in partnership with King's College London, United Kingdom. IT is supported by the Oxford Biomedical Research Centre. BOCS is supported by funds from Cancer Research UK (C8620/A8372/A15106) and the Institute of Cancer Research (UK). BOCS acknowledges NHS funding to the Royal Marsden / Institute of Cancer Research NIHR Specialist Cancer Biomedical Research Centre. The BREast Oncology GAlician Network (BREOGAN) is funded by Acción Estratégica de Salud del Instituto de Salud Carlos III FIS PI12/02125/Cofinanciado FEDER; Acción Estratégica de Salud del Instituto de Salud Carlos III FIS Intrasalud (PI13/01136); Programa Grupos Emergentes, Cancer Genetics Unit, Instituto de Investigacion Biomedica Galicia Sur. Xerencia de Xestion Integrada de Vigo-SERGAS, Instituto de Salud Carlos III, Spain; Grant 10CSA012E, Consellería de Industria Programa Sectorial de Investigación Aplicada, PEME I + D e I + D Suma del Plan Gallego de Investigación, Desarrollo e Innovación Tecnológica de la Consellería de Industria de la Xunta de Galicia, Spain; Grant EC11-192. Fomento de la Investigación Clínica Independiente, Ministerio de Sanidad, Servicios Sociales e Igualdad, Spain; and Grant FEDER-Innterconecta. Ministerio de Economia y Competitividad, Xunta de Galicia, Spain. The BSUCH study was supported by the Dietmar-Hopp Foundation, the Helmholtz Society and the German Cancer Research Center (DKFZ). The CAMA study was funded by Consejo Nacional de Ciencia y Tecnología (CONACyT) (SALUD-2002-C01-7462). Sample collection and processing was funded in part by grants from the National Cancer Institute (NCI R01CA120120 and K24CA169004). CBCS is funded by the Canadian Cancer Society (grant # 313404) and the Canadian Institutes of Health Research. CCGP is supported by funding from the University of Crete. The CECILE study was supported by Fondation de France, Institut National du Cancer (INCa), Ligue Nationale contre le Cancer, Agence Nationale de Sécurité Sanitaire, de l'Alimentation, de l'Environnement et du Travail (ANSES), Agence Nationale de la Recherche (ANR). The CGPS was supported by the Chief Physician Johan Boserup and Lise Boserup Fund, the Danish Medical Research Council, and Herlev and Gentofte Hospital. The CNIO-BCS was supported by the Instituto de Salud Carlos III, the Red Temática de Investigación Cooperativa en Cáncer and grants from the Asociación Española Contra el Cáncer and the Fondo de Investigación Sanitario (PI11/00923 and PI12/00070). COLBCCC is supported by the German Cancer Research Center (DKFZ), Heidelberg, Germany. Diana Torres was in part supported by a postdoctoral fellowship from the Alexander von Humboldt Foundation. The American Cancer Society funds the creation, maintenance, and updating of the CPS-II cohort. The CTS was initially supported by the California Breast Cancer Act of 1993 and the California Breast Cancer Research Fund (contract 97-10500) and is currently funded through the National Institutes of Health (R01 CA77398, UM1 CA164917, and U01 CA199277). Collection of cancer incidence data was supported by the California Department of Public Health as part of the statewide cancer reporting program mandated by California Health and Safety Code Section 103885. HAC receives support from the Lon V Smith Foundation (LVS39420). The University of Westminster curates the DietCompLyf database funded by Against Breast Cancer Registered Charity No. 1121258 and the NCRN. The coordination of EPIC is financially supported by the European Commission (DG-SANCO) and the International Agency for Research on Cancer. The national cohorts are supported by: Ligue Contre le Cancer, Institut Gustave Roussy, Mutuelle Générale de l'Education Nationale,

Institut National de la Santé et de la Recherche Médicale (INSERM) (France); German Cancer Aid, German Cancer Research Center (DKFZ), Federal Ministry of Education and Research (BMBF) (Germany); the Hellenic Health Foundation, the Stavros Niarchos Foundation (Greece); Associazione Italiana per la Ricerca sul Cancro-AIRC-Italy and National Research Council (Italy); Dutch Ministry of Public Health, Welfare and Sports (VWS), Netherlands Cancer Registry (NKR), LK Research Funds, Dutch Prevention Funds, Dutch ZON (Zorg Onderzoek Nederland), World Cancer Research Fund (WCRF), Statistics Netherlands (The Netherlands); Health Research Fund (FIS), PI13/00061 to Granada, PI13/01162 to EPIC-Murcia, Regional Governments of Andalucía, Asturias, Basque Country, Murcia and Navarra, ISCIII RETIC (RD06/0020) (Spain); Cancer Research UK (14136 to EPIC-Norfolk; C570/A16491 and C8221/A19170 to EPIC-Oxford), Medical Research Council (1000143 to EPIC-Norfolk, MR/M012190/1 to EPIC-Oxford) (United Kingdom). The ESTHER study was supported by a grant from the Baden Württemberg Ministry of Science, Research and Arts. Additional cases were recruited in the context of the VERDI study, which was supported by a grant from the German Cancer Aid (Deutsche Krebshilfe). FHRISK is funded from NIHR grant PGfAR 0707-10031. The GC-HBOC (German Consortium of Hereditary Breast and Ovarian Cancer) is supported by the German Cancer Aid (grant no 110837, coordinator: Rita K. Schmutzler, Cologne). This work was also funded by the European Regional Development Fund and Free State of Saxony, Germany (LIFE - Leipzig Research Centre for Civilization Diseases, project numbers 713-241202, 713-241202, 14505/2470, 14575/2470). The GENICA was funded by the Federal Ministry of Education and Research (BMBF) Germany grants 01KW9975/5, 01KW9976/8, 01KW9977/0 and 01KW0114, the Robert Bosch Foundation, Stuttgart, Deutsches Krebsforschungszentrum (DKFZ), Heidelberg, the Institute for Prevention and Occupational Medicine of the German Social Accident Insurance, Institute of the Ruhr University Bochum (IPA), Bochum, as well as the Department of Internal Medicine, Evangelische Kliniken Bonn gGmbH, Johanniter Krankenhaus, Bonn, Germany. The GEPARSIXTO study was conducted by the German Breast Group GmbH. The GESBC was supported by the Deutsche Krebshilfe e. V. [70492] and the German Cancer Research Center (DKFZ). GLACIER was supported by Breast Cancer Now, CRUK and Biomedical Research Centre at Guy's and St Thomas' NHS Foundation Trust and King's College London. The HABCS study was supported by the Claudia von Schilling Foundation for Breast Cancer Research, by the Lower Saxonian Cancer Society, and by the Rudolf Bartling Foundation. The HEBCS was financially supported by the Helsinki University Central Hospital Research Fund, Academy of Finland (266528), the Finnish Cancer Society, and the Sigrid Juselius Foundation. The HERPACC was supported by MEXT Kakenhi (No. 170150181 and 26253041) from the Ministry of Education, Science, Sports, Culture and Technology of Japan, by a Grant-in-Aid for the Third Term Comprehensive 10-Year Strategy for Cancer Control from Ministry Health, Labour and Welfare of Japan, by Health and Labour Sciences Research Grants for Research on Applying Health Technology from Ministry Health, Labour and Welfare of Japan, by National Cancer Center Research and Development Fund, and "Practical Research for Innovative Cancer Control (15ck0106177h0001)" from Japan Agency for Medical Research and development, AMED, and Cancer Bio Bank Aichi. The HMBCS was supported by a grant from the Friends of Hannover Medical School and by the Rudolf Bartling Foundation. The HUBCS was supported by a grant from the German Federal Ministry of Research and Education (RUS08/017), and by the Russian Foundation for Basic Research and the Federal Agency for Scientific Organizations for support the Bioresource collections and RFBR grants 14-04-97088, 17-29-06014 and 17-44-020498. ICICLE was supported by Breast Cancer Now, CRUK and Biomedical Research

Centre at Guy's and St Thomas' NHS Foundation Trust and King's College London. Financial support for KARBAC was provided through the regional agreement on medical training and clinical research (ALF) between Stockholm County Council and Karolinska Institutet, the Swedish Cancer Society, The Gustav V Jubilee foundation and Bert von Kantzows foundation. The KARMA study was supported by Märit and Hans Rausings Initiative Against Breast Cancer. The KBCP was financially supported by the special Government Funding (EVO) of Kuopio University Hospital grants, Cancer Fund of North Savo, the Finnish Cancer Organizations, and by the strategic funding of the University of Eastern Finland. kConFab is supported by a grant from the National Breast Cancer Foundation, and previously by the National Health and Medical Research Council (NHMRC), the Queensland Cancer Fund, the Cancer Councils of New South Wales, Victoria, Tasmania and South Australia, and the Cancer Foundation of Western Australia. Financial support for the AOCS was provided by the United States Army Medical Research and Materiel Command [DAMD17-01-1-0729], Cancer Council Victoria, Queensland Cancer Fund, Cancer Council New South Wales, Cancer Council South Australia, The Cancer Foundation of Western Australia, Cancer Council Tasmania and the National Health and Medical Research Council of Australia (NHMRC; 400413, 400281, 199600). G.C.T. and P.W. are supported by the NHMRC. RB was a Cancer Institute NSW Clinical Research Fellow. The KOHBRA study was partially supported by a grant from the Korea Health Technology R&D Project through the Korea Health Industry Development Institute (KHIDI), and the National R&D Program for Cancer Control, Ministry of Health & Welfare, Republic of Korea (HI16C1127; 1020350; 1420190). LAABC is supported by grants (1RB-0287, 3PB-0102, 5PB-0018, 10PB-0098) from the California Breast Cancer Research Program. Incident breast cancer cases were collected by the USC Cancer Surveillance Program (CSP) which is supported under subcontract by the California Department of Health. The CSP is also part of the National Cancer Institute's Division of Cancer Prevention and Control Surveillance, Epidemiology, and End Results Program, under contract number N01CN25403. LMBC is supported by the 'Stichting tegen Kanker'. DL is supported by the FWO. The MABCS study is funded by the Research Centre for Genetic Engineering and Biotechnology "Georgi D. Efremov" and supported by the German Academic Exchange Program, DAAD. The MARIE study was supported by the Deutsche Krebshilfe e.V. [70-2892-BR I, 106332, 108253, 108419, 110826, 110828], the Hamburg Cancer Society, the German Cancer Research Center (DKFZ) and the Federal Ministry of Education and Research (BMBF) Germany [01KH0402]. MBCSG is supported by grants from the Italian Association for Cancer Research (AIRC) and by funds from the Italian citizens who allocated the 5/1000 share of their tax payment in support of the Fondazione IRCCS Istituto Nazionale Tumori, according to Italian laws (INT-Institutional strategic projects "5x1000"). The MCBCS was supported by the NIH grants CA192393, CA116167, CA176785 an NIH Specialized Program of Research Excellence (SPORE) in Breast Cancer [CA116201], and the Breast Cancer Research Foundation and a generous gift from the David F. and Margaret T. Grohne Family Foundation. MCCS cohort recruitment was funded by VicHealth and Cancer Council Victoria. The MCCS was further supported by Australian NHMRC grants 209057 and 396414, and by infrastructure provided by Cancer Council Victoria. Cases and their vital status were ascertained through the Victorian Cancer Registry (VCR) and the Australian Institute of Health and Welfare (AIHW), including the National Death Index and the Australian Cancer Database. The MEC was support by NIH grants CA63464, CA54281, CA098758, CA132839 and CA164973. The MISS study is supported by funding from ERC-2011-294576 Advanced grant, Swedish Cancer Society, Swedish Research Council, Local hospital funds, Berta Kamprad Foundation, Gunnar Nilsson. The MMHS study was supported by NIH grants CA97396, CA128931, CA116201, CA140286 and CA177150. MSKCC is supported by grants from the Breast Cancer Research Foundation and Robert and Kate Niehaus Clinical Cancer Genetics Initiative. The work of MTLGEBCS was supported by the Quebec Breast Cancer Foundation, the Canadian Institutes of Health Research for the “CIHR Team in Familial Risks of Breast Cancer” program – grant # CRN-87521 and the Ministry of Economic Development, Innovation and Export Trade – grant # PSR-SIIRI-701. MYBRCA is funded by research grants from the Malaysian Ministry of Higher Education (UM.C/HlR/MOHE/06) and Cancer Research Malaysia. MYMAMMO is supported by research grants from Yayasan Sime Darby LPGA Tournament and Malaysian Ministry of Higher Education (RP046B-15HTM). The NBCS has received funding from the K.G. Jebsen Centre for Breast Cancer Research; the Research Council of Norway grant 193387/V50 (to A-L Børresen-Dale and V.N. Kristensen) and grant 193387/H10 (to A-L Børresen-Dale and V.N. Kristensen), South Eastern Norway Health Authority (grant 39346 to A-L Børresen-Dale) and the Norwegian Cancer Society (to A-L Børresen-Dale and V.N. Kristensen). The NBHS was supported by NIH grant R01CA100374. Biological sample preparation was conducted the Survey and Biospecimen Shared Resource, which is supported by P30 CA68485. The Northern California Breast Cancer Family Registry (NC-BCFR) and Ontario Familial Breast Cancer Registry (OFBCR) were supported by grant UM1 CA164920 from the National Cancer Institute (USA). The content of this manuscript does not necessarily reflect the views or policies of the National Cancer Institute or any of the collaborating centers in the Breast Cancer Family Registry (BCFR), nor does mention of trade names, commercial products, or organizations imply endorsement by the USA Government or the BCFR. The Carolina Breast Cancer Study was funded by Komen Foundation, the National Cancer Institute (P50 CA058223, U54 CA156733, U01 CA179715), and the North Carolina University Cancer Research Fund. The NGOBCS was supported by Grants-in-Aid for the Third Term Comprehensive Ten-Year Strategy for Cancer Control from the Ministry of Health, Labor and Welfare of Japan, and for Scientific Research on Priority Areas, 17015049 and for Scientific Research on Innovative Areas, 221S0001, from the Ministry of Education, Culture, Sports, Science, and Technology of Japan. The NHS was supported by NIH grants P01 CA87969, UM1 CA186107, and U19 CA148065. The NHS2 was supported by NIH grants UM1 CA176726 and U19 CA148065. The OBCS was supported by research grants from the Finnish Cancer Foundation, the Academy of Finland (grant number 250083, 122715 and Center of Excellence grant number 251314), the Finnish Cancer Foundation, the Sigrid Juselius Foundation, the University of Oulu, the University of Oulu Support Foundation and the special Governmental EVO funds for Oulu University Hospital-based research activities. The ORIGO study was supported by the Dutch Cancer Society (RUL 1997-1505) and the Biobanking and Biomolecular Resources Research Infrastructure (BBMRI-NL CP16). The PBCS was funded by Intramural Research Funds of the National Cancer Institute, Department of Health and Human Services, USA. Genotyping for PLCO was supported by the Intramural Research Program of the National Institutes of Health, NCI, Division of Cancer Epidemiology and Genetics. The PLCO is supported by the Intramural Research Program of the Division of Cancer Epidemiology and Genetics and supported by contracts from the Division of Cancer Prevention, National Cancer Institute, National Institutes of Health. The POSH study is funded by Cancer Research UK (grants C1275/A11699, C1275/C22524, C1275/A19187, C1275/A15956 and Breast Cancer Campaign 2010PR62, 2013PR044. PROCAS is funded from NIHR grant PGfAR 0707-10031. The RBCS was funded by the Dutch Cancer Society (DDHK 2004-3124, DDHK 2009-4318). The SASBAC study was supported by funding from the Agency for Science, Technology and Research of Singapore (A*STAR), the US National Institute of Health (NIH) and the Susan G.

Komen Breast Cancer Foundation. The SBCGS was supported primarily by NIH grants R01CA64277, R01CA148667, UMCA182910, and R37CA70867. Biological sample preparation was conducted the Survey and Biospecimen Shared Resource, which is supported by P30 CA68485. The scientific development and funding of this project were, in part, supported by the Genetic Associations and Mechanisms in Oncology (GAME-ON) Network U19 CA148065. The SBCS was supported by Sheffield Experimental Cancer Medicine Centre and Breast Cancer Now Tissue Bank. The SCCS is supported by a grant from the National Institutes of Health (R01 CA092447). Data on SCCS cancer cases used in this publication were provided by the Alabama Statewide Cancer Registry; Kentucky Cancer Registry, Lexington, KY; Tennessee Department of Health, Office of Cancer Surveillance; Florida Cancer Data System; North Carolina Central Cancer Registry, North Carolina Division of Public Health; Georgia Comprehensive Cancer Registry; Louisiana Tumor Registry; Mississippi Cancer Registry; South Carolina Central Cancer Registry; Virginia Department of Health, Virginia Cancer Registry; Arkansas Department of Health, Cancer Registry, 4815 W. Markham, Little Rock, AR 72205. The Arkansas Central Cancer Registry is fully funded by a grant from National Program of Cancer Registries, Centers for Disease Control and Prevention (CDC). Data on SCCS cancer cases from Mississippi were collected by the Mississippi Cancer Registry which participates in the National Program of Cancer Registries (NPCR) of the Centers for Disease Control and Prevention (CDC). The contents of this publication are solely the responsibility of the authors and do not necessarily represent the official views of the CDC or the Mississippi Cancer Registry. SEARCH is funded by Cancer Research UK [C490/A10124, C490/A16561] and supported by the UK National Institute for Health Research Biomedical Research Centre at the University of Cambridge. The University of Cambridge has received salary support for PDPP from the NHS in the East of England through the Clinical Academic Reserve. SEBCS was supported by the BRL (Basic Research Laboratory) program through the National Research Foundation of Korea funded by the Ministry of Education, Science and Technology (2012-0000347). SGBCC is funded by the NUS start-up Grant, National University Cancer Institute Singapore (NCIS) Centre Grant and the NMRC Clinician Scientist Award. Additional controls were recruited by the Singapore Consortium of Cohort Studies-Multi-ethnic cohort (SCCS-MEC), which was funded by the Biomedical Research Council, grant number: 05/1/21/19/425. The Sister Study (SISTER) is supported by the Intramural Research Program of the NIH, National Institute of Environmental Health Sciences (Z01-ES044005 and Z01-ES049033). The Two Sister Study (2SISTER) was supported by the Intramural Research Program of the NIH, National Institute of Environmental Health Sciences (Z01-ES044005 and Z01-ES102245), and, also by a grant from Susan G. Komen for the Cure, grant FAS0703856. SKKDKFZS is supported by the DKFZ. The SMC is funded by the Swedish Cancer Foundation. The SZBCS was supported by Grant PBZ_KBN_122/P05/2004. The TBCS was funded by The National Cancer Institute Thailand. The TNBCC was supported by: a Specialized Program of Research Excellence (SPORE) in Breast Cancer (CA116201), a grant from the Breast Cancer Research Foundation, a generous gift from the David F. and Margaret T. Grohne Family Foundation. The TWBCS is supported by the Taiwan Biobank project of the Institute of Biomedical Sciences, Academia Sinica, Taiwan. The UCIBCS component of this research was supported by the NIH [CA58860, CA92044] and the Lon V Smith Foundation [LVS39420]. The UKBGS is funded by Breast Cancer Now and the Institute of Cancer Research (ICR), London. ICR acknowledges NHS funding to the NIHR Biomedical Research Centre. The UKOPS study was funded by The Eve Appeal (The Oak Foundation) and supported by the National Institute for Health Research University College London Hospitals Biomedical Research Centre. The US3SS study was supported by

Massachusetts (K.M.E., R01CA47305), Wisconsin (P.A.N., R01 CA47147) and New Hampshire (L.T.-E., R01CA69664) centers, and Intramural Research Funds of the National Cancer Institute, Department of Health and Human Services, USA. The USRT Study was funded by Intramural Research Funds of the National Cancer Institute, Department of Health and Human Services, USA. The WAABCS study was supported by grants from the National Cancer Institute of the National Institutes of Health (R01 CA89085 and P50 CA125183 and the D43 TW009112 grant), Susan G. Komen (SAC110026), the Dr. Ralph and Marian Falk Medical Research Trust, and the Avon Foundation for Women. The WHI program is funded by the National Heart, Lung, and Blood Institute, the US National Institutes of Health and the US Department of Health and Human Services (HHSN268201100046C, HHSN268201100001C, HHSN268201100002C, HHSN268201100003C, HHSN268201100004C and HHSN271201100004C). This work was also funded by NCI U19 CA148065-01.

**Acknowledgements**
We thank all the individuals who took part in these studies and all the researchers, clinicians, technicians and administrative staff who have enabled this work to be carried out. The COGS study would not have been possible without the contributions of the following:) Andrew Berchuck (OCAC), Rosalind A. Eeles, Ali Amin Al Olama, Zsofia Kote-Jarai, Sara Benlloch (PRACTICAL), Andrew Lee, and Ed Dicks, Craig Luccarini and the staff of the Centre for Genetic Epidemiology Laboratory, and the staff of the CNIO genotyping unit, Daniel C. Tessier, Francois Bacot, Daniel Vincent, Sylvie LaBoissière and Frederic Robidoux and the staff of the McGill University and Génome Québec Innovation Centre, Sune F. Nielsen, Borge G. Nordestgaard, and the staff of the Copenhagen DNA laboratory, and Julie M. Cunningham, Sharon A. Windebank, Christopher A. Hilker, Jeffrey Meyer and the staff of Mayo Clinic Genotyping Core Facility. ABCFS thank Maggie Angelakos, Judi Maskiell, Gillian Dite. ABCS thanks the Blood bank Sanquin, The Netherlands. ABCTB Investigators: Christine Clarke,Rosemary Balleine, Robert Baxter,Stephen Braye, Jane Carpenter, Jane Dahlstrom, John Forbes, Soon Lee, Debbie Marsh, Adrienne Morey, Nirmala Pathmanathan, Rodney Scott, Allan Spigelman, Nicholas Wilcken, Desmond Yip. Samples are made available to researchers on a non-exclusive basis. The ACP study wishes to thank the participants in the Thai Breast Cancer study. Special Thanks also go to the Thai Ministry of Public Health (MOPH), doctors and nurses who helped with the data collection process. Finally, the study would like to thank Dr Prat Boonyawongviroj, the former Permanent Secretary of MOPH and Dr Pornthep Siriwanarungsan, the Department Director-General of Disease Control who have supported the study throughout. BBCS thanks Eileen Williams, Elaine Ryder-Mills, Kara Sargus. BCEES thanks Allyson Thomson, Christobel Saunders, Terry Slevin, BreastScreen Western Australia, Elizabeth Wylie, Rachel Lloyd. The BCINIS study would not have been possible without the contributions of Dr. K. Landsman, Dr. N. Gronich, Dr. A. Flugelman, Dr. W. Saliba, Dr. E. Liani, Dr. I. Cohen, Dr. S. Kalet, Dr. V. Friedman, Dr. O. Barnet of the NICCC in Haifa, and all the contributing family medicine, surgery, pathology and oncology teams in all medical institutes in Northern Israel. BIGGS thanks Niall McInerney, Gabrielle Colleran, Andrew Rowan, Angela Jones. The BREOGAN study would not have been possible without the contributions of the following: Manuela Gago-Dominguez, Jose Esteban Castelao, Angel Carracedo, Victor Muñoz Garzón, Alejandro Novo Domínguez, Maria Elena Martinez, Sara Miranda Ponte, Carmen Redondo Marey, Maite Peña Fernández, Manuel Enguix Castelo, Maria Torres, Manuel Calaza (BREOGAN), José Antúnez, Máximo Fraga and the staff of the Department of Pathology and Biobank of the University Hospital Complex of Santiago-CHUS,

Instituto de Investigación Sanitaria de Santiago, IDIS, Xerencia de Xestion Integrada de Santiago-SERGAS; Joaquín González-Carreró and the staff of the Department of Pathology and Biobank of University Hospital Complex of Vigo, Instituto de Investigacion Biomedica Galicia Sur, SERGAS, Vigo, Spain. BSUCH thanks Peter Bugert, Medical Faculty Mannheim. The CAMA study would like to recognize CONACyT for the financial support provided for this work and all physicians responsible for the project in the different participating hospitals: Dr. Germán Castelazo (IMSS, Ciudad de México, DF), Dr. Sinhué Barroso Bravo (IMSS, Ciudad de México, DF), Dr. Fernando Mainero Ratchelous (IMSS, Ciudad de México, DF), Dr. Joaquín Zarco Méndez (ISSSTE, Ciudad de México, DF), Dr. Edelmiro Pérez Rodríguez (Hospital Universitario, Monterrey, Nuevo León), Dr. Jesús Pablo Esparza Cano (IMSS, Monterrey, Nuevo León), Dr. Heriberto Fabela (IMSS, Monterrey, Nuevo León), Dr. Fausto Hernández Morales (ISSSTE, Veracruz, Veracruz), Dr. Pedro Coronel Brizio (CECAN SS, Xalapa, Veracruz) and Dr. Vicente A. Saldaña Quiroz (IMSS, Veracruz, Veracruz). CBCS thanks study participants, co-investigators, collaborators and staff of the Canadian Breast Cancer Study, and project coordinators Agnes Lai and Celine Morissette. CCGP thanks Styliani Apostolaki, Anna Margiolaki, Georgios Nintos, Maria Perraki, Georgia Saloustrou, Georgia Sevastaki, Konstantinos Pompodakis. CGPS thanks staff and participants of the Copenhagen General Population Study. For the excellent technical assistance: Dorthe Uldall Andersen, Maria Birna Arnadottir, Anne Bank, Dorthe Kjeldgård Hansen. The Danish Cancer Biobank is acknowledged for providing infrastructure for the collection of blood samples for the cases. CNIO-BCS thanks Guillermo Pita, Charo Alonso, Nuria Álvarez, Pilar Zamora, Primitiva Menendez, the Human Genotyping-CEGEN Unit (CNIO). COLBCCC thanks all patients, the physicians Justo G. Olaya, Mauricio Tawil, Lilian Torregrosa, Elias Quintero, Sebastian Quintero, Claudia Ramírez, José J. Caicedo, and Jose F. Robledo, the researchers Diana Torres, Ignacio Briceno, Fabian Gil, Angela Umana, Angela Beltran and Viviana Ariza, and the technician Michael Gilbert for their contributions and commitment to this study. Investigators from the CPS-II cohort thank the participants and Study Management Group for their invaluable contributions to this research. They also acknowledge the contribution to this study from central cancer registries supported through the Centers for Disease Control and Prevention National Program of Cancer Registries, as well as cancer registries supported by the National Cancer Institute Surveillance Epidemiology and End Results program. The CTS Steering Committee includes Leslie Bernstein, Susan Neuhausen, James Lacey, Sophia Wang, Huiyan Ma, and Jessica Clague DeHart at the Beckman Research Institute of City of Hope, Dennis Deapen, Rich Pinder, and Eunjung Lee at the University of Southern California, Pam Horn-Ross, Peggy Reynolds, Christina Clarke Dur and David Nelson at the Cancer Prevention Institute of California, Hoda Anton-Culver, Argyrios Ziogas, and Hannah Park at the University of California Irvine, and Fred Schumacher at Case Western University. DIETCOMPLYF thanks the patients, nurses and clinical staff involved in the study. The DietCompLyf study was funded by the charity Against Breast Cancer (Registered Charity Number 1121258) and the NCRN. We thank the participants and the investigators of EPIC (European Prospective Investigation into Cancer and Nutrition). ESTHER thanks Hartwig Ziegler, Sonja Wolf, Volker Hermann, Christa Stegmaier, Katja Butterbach. FHRISK thanks NIHR for funding. GC-HBOC thanks Stefanie Engert, Heide Hellebrand, Sandra Kröber and LIFE - Leipzig Research Centre for Civilization Diseases (Markus Loeffler, Joachim Thiery, Matthias Nüchter, Ronny Baber). The GENICA Network: Dr. Margarete Fischer-Bosch-Institute of Clinical Pharmacology, Stuttgart, and University of Tübingen, Germany [HB, Wing-Yee Lo, Christina Justenhoven], German Cancer Consortium (DKTK) and German Cancer Research Center (DKFZ) [HB], Department of Internal Medicine, Evangelische Kliniken Bonn gGmbH, Johanniter Krankenhaus, Bonn, Germany [Yon-Dschun Ko, Christian Baisch], Institute of Pathology, University of Bonn, Germany [Hans-Peter Fischer], Molecular Genetics of Breast Cancer, Deutsches Krebsforschungszentrum (DKFZ), Heidelberg, Germany [Ute Hamann], Institute for Prevention and Occupational Medicine of the German Social Accident Insurance, Institute of the Ruhr University Bochum (IPA), Bochum, Germany [Thomas Brüning, Beate Pesch, Sylvia Rabstein, Anne Lotz]; and Institute of Occupational Medicine and Maritime Medicine, University Medical Center Hamburg-Eppendorf, Germany [Volker Harth]. GLACIER thanks Kelly Kohut, Patricia Gorman, Maria Troy. HABCS thanks Michael Bremer. HEBCS thanks Sofia Khan, Johanna Kiiski, Carl Blomqvist, Kristiina Aittomäki, Rainer Fagerholm, Kirsimari Aaltonen, Karl von Smitten, Irja Erkkilä. HKBCS thanks Hong Kong Sanatorium and Hospital, Dr Ellen Li Charitable Foundation, The Kerry Group Kuok Foundation, National Institute of Health 1R03CA130065 and the North California Cancer Center for support. HMBCS thanks Peter Hillemanns, Hans Christiansen and Johann H. Karstens. HUBCS thanks Shamil Gantsev. ICICLE thanks Kelly Kohut, Michele Caneppele, Maria Troy. KARMA and SASBAC thank the Swedish Medical Research Counsel. KBCP thanks Eija Myöhänen, Helena Kemiläinen. kConFab/AOCS wish to thank Heather Thorne, Eveline Niedermayr, all the kConFab research nurses and staff, the heads and staff of the Family Cancer Clinics, and the Clinical Follow Up Study (which has received funding from the NHMRC, the National Breast Cancer Foundation, Cancer Australia, and the National Institute of Health (USA)) for their contributions to this resource, and the many families who contribute to kConFab. We thank all investigators of the KOHBRA (Korean Hereditary Breast Cancer) Study. LAABC thanks all the study participants and the entire data collection team, especially Annie Fung and June Yashiki. LMBC thanks Gilian Peuteman, Thomas Van Brussel, EvyVanderheyden and Kathleen Corthouts. MABCS thanks Milena Jakimovska (RCGEB "Georgi D. Efremov), Katerina Kubelka, Mitko Karadjozov (Adzibadem-Sistina" Hospital), Andrej Arsovski and Liljana Stojanovska (Re-Medika" Hospital) for their contributions and commitment to this study. MARIE thanks Petra Seibold, Dieter Flesch-Janys, Judith Heinz, Nadia Obi, Alina Vrieling, Sabine Behrens, Ursula Eilber, Muhabbet Celik, Til Olchers and Stefan Nickels. MBCSG (Milan Breast Cancer Study Group): Paolo Radice, Paolo Peterlongo, Siranoush Manoukian, Bernard Peissel, Roberta Villa, Cristina Zanzottera, Bernardo Bonanni, Irene Feroce, and the personnel of the Cogentech Cancer Genetic Test Laboratory. We thank the coordinators, the research staff and especially the MMHS participants for their continued collaboration on research studies in breast cancer. MSKCC thanks Marina Corines, Lauren Jacobs. MTLGEBCS would like to thank Martine Tranchant (CHU de Québec Research Center), Marie-France Valois, Annie Turgeon and Lea Heguy (McGill University Health Center, Royal Victoria Hospital; McGill University) for DNA extraction, sample management and skillful technical assistance. J.S. is Chairholder of the Canada Research Chair in Oncogenetics. MYBRCA thanks study participants and research staff (particularly Patsy Ng, Nurhidayu Hassan, Yoon Sook-Yee, Daphne Lee, Lee Sheau Yee, Phuah Sze Yee and Norhashimah Hassan) for their contributions and commitment to this study. The following are NBCS Collaborators: Kristine K. Sahlberg (PhD), Lars Ottestad (MD), Rolf Kåresen (Prof. Em.) Dr. Ellen Schlichting (MD), Marit Muri Holmen (MD), Toril Sauer (MD), Vilde Haakensen (MD), Olav Engebråten (MD), Bjørn Naume (MD), Alexander Fosså (MD), Cecile E. Kiserud (MD), Kristin V. Reinertsen (MD), Åslaug Helland (MD), Margit Riis (MD), Jürgen Geisler (MD), OSBREAC and Grethe I. Grenaker Alnæs (MSc). NBHS and SBCGS thank study participants and research staff for their contributions and commitment to the studies. We would like to thank the participants and staff of the Nurses' Health Study and Nurses' Health Study II for their valuable contributions as well as the following state cancer registries for their help: AL, AZ, AR, CA, CO, CT, DE, FL, GA, ID, IL, IN, IA, KY, LA, ME, MD, MA, MI, NE, NH, NJ, NY, NC, ND, OH, OK, OR, PA, RI, SC, TN, TX, VA, WA, WY. The authors assume full responsibility for analyses and interpretation of these data. OBCS thanks Arja Jukkola-Vuorinen, Mervi Grip, Saila Kauppila, Meeri Otsukka, Leena Keskitalo and Kari Mononen for their contributions to this study. OFBCR thanks Teresa Selander, Nayana Weerasooriya. ORIGO thanks E. Krol-Warmerdam, and J. Blom for patient accrual, administering questionnaires, and managing clinical information. The LUMC survival data were retrieved from the Leiden hospital-based cancer registry system (ONCDOC) with the help of Dr. J. Molenaar. PBCS thanks Louise Brinton, Mark Sherman, Neonila Szeszenia-Dabrowska, Beata Peplonska, Witold Zatonski, Pei Chao, Michael Stagner. The ethical approval for the POSH study is MREC /00/6/69, UKCRN ID: 1137. We thank staff in the Experimental Cancer Medicine Centre (ECMC) supported Faculty of Medicine Tissue Bank and the Faculty of Medicine DNA Banking resource. PREFACE thanks Sonja Oeser and Silke Landrith. PROCAS thanks NIHR for funding. RBCS thanks Petra Bos, Jannet Blom, Ellen Crepin, Elisabeth Huijskens, Anja Kromwijk-Nieuwlaat, Annette Heemskerk, the Erasmus MC Family Cancer Clinic. SBCS thanks Sue Higham, Helen Cramp, Dan Connley, Ian Brock, Sabapathy Balasubramanian and Malcolm W.R. Reed. We thank the SEARCH and EPIC teams. SGBCC thanks the participants and research coordinator Ms Tan Siew Li. SKKDKFZS thanks all study participants, clinicians, family doctors, researchers and technicians for their contributions and commitment to this study. We thank the SUCCESS Study teams in Munich, Duessldorf, Erlangen and Ulm. SZBCS thanks Ewa Putresza. UCIBCS thanks Irene Masunaka. UKBGS thanks Breast Cancer Now and the Institute of Cancer Research for support and funding of the Breakthrough Generations Study, and the study participants, study staff, and the doctors, nurses and other health care providers and health information sources who have contributed to the study. We acknowledge NHS funding to the Royal Marsden/ICR NIHR Biomedical Research Centre. The authors thank the WHI investigators and staff for their dedication and the study participants for making the program possible.

**CIMBA FUNDING AND ACKNOWLEDGEMENTS**

**Funding**

CIMBA: The CIMBA data management and data analysis were supported by Cancer Research – UK grants C12292/A20861, C12292/A11174. ACA is a Cancer Research -UK Senior Cancer Research Fellow. GCT and ABS are NHMRC Research Fellows. iCOGS: the European Community's Seventh Framework Programme under grant agreement n° 223175 (HEALTH-F2-2009-223175) (COGS), Cancer Research UK (C1287/A10118, C1287/A 10710, C12292/A11174, C1281/A12014, C5047/A8384, C5047/A15007, C5047/A10692, C8197/A16565), the National Institutes of Health (CA128978) and Post-Cancer GWAS initiative (1U19 CA148537, 1U19 CA148065 and 1U19 CA148112 - the GAME-ON initiative), the Department of Defence (W81XWH-10-1-0341), the Canadian Institutes of Health Research (CIHR) for the CIHR Team in Familial Risks of Breast Cancer (CRN-87521), and the Ministry of Economic Development, Innovation and Export Trade (PSR-SIIRI-701), Komen Foundation for the Cure, the Breast Cancer Research Foundation, and the Ovarian Cancer Research Fund. The PERSPECTIVE project was supported by the Government of Canada through Genome Canada and the Canadian Institutes of Health Research, the Ministry of Economy, Science and Innovation through Genome Québec, and The Quebec Breast Cancer Foundation.

BCFR: UM1 CA164920 from the National Cancer Institute. The content of this manuscript does not necessarily reflect the views or policies of the National Cancer Institute or any of the collaborating centers in the Breast Cancer Family Registry (BCFR), nor does mention of trade names, commercial products, or organizations imply endorsement by the US Government or the BCFR. BFBOCC: Lithuania (BFBOCC-LT): Research Council of Lithuania grant SEN-18/2015. BIDMC: Breast Cancer Research Foundation. BMBSA: Cancer Association of South Africa (PI Elizabeth J. van Rensburg). CNIO: Spanish Ministry of Health PI16/00440 supported by FEDER funds, the Spanish Ministry of Economy and Competitiveness (MINECO) SAF2014-57680-R and the Spanish Research Network on Rare diseases (CIBERER). COH-CCGCRN: Research reported in this publication was supported by the National Cancer Institute of the National Institutes of Health under grant number R25CA112486, and RC4CA153828 (PI: J. Weitzel) from the National Cancer Institute and the Office of the Director, National Institutes of Health. The content is solely the responsibility of the authors and does not necessarily represent the official views of the National Institutes of Health. CONSIT TEAM: Associazione Italiana Ricerca sul Cancro (AIRC; IG2014 no.15547) to P. Radice. Funds from Italian citizens who allocated the 5x1000 share of their tax payment in support of the Fondazione IRCCS Istituto Nazionale Tumori, according to Italian laws (INT-Institutional strategic projects '5x1000') to S. Manoukian. **UNIROMA1**: Italian Association for Cancer Research (AIRC; grant no.16933) to L. Ottini. **IST:** Funds from Italian citizens who allocated the 5x1000 share of their tax payment in support of the Ospedale Policlinico San Martino IRCCS according to Italian laws (institutional project) to L. Varesco. **UNINS:** Associazione CAOS Varese to M.G. Tibiletti. **IFOM:** Associazione Italiana Ricerca sul Cancro (AIRC; IG2015 no.16732) to P. Peterlongo**.** DEMOKRITOS: European Union (European Social Fund – ESF) and Greek national funds through the Operational Program "Education and Lifelong Learning" of the National Strategic Reference Framework (NSRF) - Research Funding Program of the General Secretariat for Research & Technology: SYN11_10_19 NBCA. Investing in knowledge society through the European Social Fund. DFKZ: German Cancer Research Center. EMBRACE: Cancer Research UK Grants C1287/A10118 and C1287/A11990. D. Gareth Evans and Fiona Lalloo are supported by an NIHR grant to the Biomedical Research Centre, Manchester. The Investigators at The Institute of Cancer Research and The Royal Marsden NHS Foundation Trust are supported by an NIHR grant to the Biomedical Research Centre at The Institute of Cancer Research and The Royal Marsden NHS Foundation Trust. Ros Eeles and Elizabeth Bancroft are supported by Cancer Research UK Grant C5047/A8385. Ros Eeles is also supported by NIHR support to the Biomedical Research Centre at The Institute of Cancer Research and The Royal Marsden NHS Foundation Trust. FCCC: The University of Kansas Cancer Center (P30 CA168524) and the Kansas Bioscience Authority Eminent Scholar Program. A.K.G. was funded by R0 1CA140323, R01 CA214545, and by the Chancellors Distinguished Chair in Biomedical Sciences Professorship. FPGMX: FISPI05/2275 and Mutua Madrileña Foundation (FMMA). GC-HBOC: German Cancer Aid (grant no 110837, Rita K. Schmutzler) and the European Regional Development Fund and Free State of Saxony, Germany (LIFE - Leipzig Research Centre for Civilization Diseases, project numbers 713-241202, 713-241202, 14505/2470, 14575/2470). GEMO: Ligue Nationale Contre le Cancer; the Association "Le cancer du sein, parlons-en!" Award, the Canadian Institutes of Health Research for the "CIHR Team in Familial Risks of Breast Cancer" program and the French National Institute of Cancer (INCa). GEORGETOWN: the Non-Therapeutic Subject Registry Shared Resource at Georgetown University (NIH/NCI grant P30-CA051008), the Fisher Center for Hereditary Cancer and Clinical Genomics Research, and Swing Fore the Cure. G-FAST: Bruce Poppe is a senior clinical investigator of

FWO. Mattias Van Heetvelde obtained funding from IWT. HCSC: Spanish Ministry of Health PI15/00059, PI16/01292, and CB-161200301 CIBERONC from ISCIII (Spain), partially supported by European Regional Development FEDER funds. HEBCS: Helsinki University Hospital Research Fund, Academy of Finland (266528), the Finnish Cancer Society and the Sigrid Juselius Foundation. HEBON: the Dutch Cancer Society grants NKI1998-1854, NKI2004-3088, NKI2007-3756, the Netherlands Organization of Scientific Research grant NWO 91109024, the Pink Ribbon grants 110005 and 2014-187.WO76, the BBMRI grant NWO 184.021.007/CP46 and the Transcan grant JTC 2012 Cancer 12-054. HEBON thanks the registration teams of Dutch Cancer Registry (IKNL; S. Siesling, J. Verloop) and the Dutch Pathology database (PALGA; L. Overbeek) for part of the data collection. HRBCP: Hong Kong Sanatorium and Hospital, Dr Ellen Li Charitable Foundation, The Kerry Group Kuok Foundation, National Institute of Health1R 03CA130065, and North California Cancer Center. HUNBOCS: Hungarian Research Grants KTIA-OTKA CK-80745 and NKFI_OTKA K-112228. ICO: The authors would like to particularly acknowledge the support of the Asociación Española Contra el Cáncer (AECC), the Instituto de Salud Carlos III (organismo adscrito al Ministerio de Economía y Competitividad) and "Fondo Europeo de Desarrollo Regional (FEDER), una manera de hacer Europa" (PI10/01422, PI13/00285, PIE13/00022, PI15/00854, PI16/00563 and CIBERONC) and the Institut Català de la Salut and Autonomous Government of Catalonia (2009SGR290, 2014SGR338 and PERIS Project MedPerCan). IHCC: PBZ_KBN_122/P05/2004. ILUH: Icelandic Association "Walking for Breast Cancer Research" and by the Landspitali University Hospital Research Fund. INHERIT: Canadian Institutes of Health Research for the "CIHR Team in Familial Risks of Breast Cancer" program – grant # CRN-87521 and the Ministry of Economic Development, Innovation and Export Trade – grant # PSR-SIIRI-701. IOVHBOCS: Ministero della Salute and "5x1000" Istituto Oncologico Veneto grant. IPOBCS: Liga Portuguesa Contra o Cancro. kConFab: The National Breast Cancer Foundation, and previously by the National Health and Medical Research Council (NHMRC), the Queensland Cancer Fund, the Cancer Councils of New South Wales, Victoria, Tasmania and South Australia, and the Cancer Foundation of Western Australia. KOHBRA: the Korea Health Technology R&D Project through the Korea Health Industry Development Institute (KHIDI), and the National R&D Program for Cancer Control, Ministry of Health & Welfare, Republic of Korea (HI16C1127; 1020350; 1420190). MAYO: NIH grants CA116167, CA192393 and CA176785, an NCI Specialized Program of Research Excellence (SPORE) in Breast Cancer (CA116201),and a grant from the Breast Cancer Research Foundation. MCGILL: Jewish General Hospital Weekend to End Breast Cancer, Quebec Ministry of Economic Development, Innovation and Export Trade. Marc Tischkowitz is supported by the funded by the European Union Seventh Framework Program (2007Y2013)/European Research Council (Grant No. 310018). MODSQUAD: MH CZ - DRO (MMCI, 00209805), **MEYS - NPS I - LO1413** to LF and by the European Regional Development Fund and the State Budget of the Czech Republic (RECAMO, CZ.1.05/2.1.00/03.0101) to LF, and by Charles University in Prague project UNCE204024 (MZ). MSKCC: the Breast Cancer Research Foundation, the Robert and Kate Niehaus Clinical Cancer Genetics Initiative, the Andrew Sabin Research Fund and a Cancer Center Support Grant/Core Grant (P30 CA008748). NAROD: 1R01 CA149429-01. NCI: the Intramural Research Program of the US National Cancer Institute, NIH, and by support services contracts NO2-CP-11019-50, N02-CP-21013-63 and N02-CP-65504 with Westat, Inc, Rockville, MD. NICCC: Clalit Health Services in Israel, the Israel Cancer Association and the Breast Cancer Research Foundation (BCRF), NY. NNPIO: the Russian Federation for Basic Research (grants 15-04-01744, 16-54-00055 and 17-54-12007). NRG Oncology: U10

CA180868, NRG SDMC grant U10 CA180822, NRG Administrative Office and the NRG Tissue Bank (CA 27469), the NRG Statistical and Data Center (CA 37517) and the Intramural Research Program, NCI. OSUCCG: Ohio State University Comprehensive Cancer Center. PBCS: Italian Association of Cancer Research (AIRC) [IG 2013 N.14477] and Tuscany Institute for Tumors (ITT) grant 2014-2015-2016. SEABASS: Ministry of Science, Technology and Innovation, Ministry of Higher Education (UM.C/HlR/MOHE/06) and Cancer Research Initiatives Foundation. SMC: the Israeli Cancer Association. SWE-BRCA: the Swedish Cancer Society. UCHICAGO: NCI Specialized Program of Research Excellence (SPORE) in Breast Cancer (CA125183), R01 CA142996, 1U01CA161032 and by the Ralph and Marion Falk Medical Research Trust, the Entertainment Industry Fund National Women's Cancer Research Alliance and the Breast Cancer research Foundation. OIO is an ACS Clinical Research Professor. UCLA: Jonsson Comprehensive Cancer Center Foundation; Breast Cancer Research Foundation. UCSF: UCSF Cancer Risk Program and Helen Diller Family Comprehensive Cancer Center. UKFOCR: Cancer Researc h UK. UPENN: National Institutes of Health (NIH) (R01-CA102776 and R01-CA083855; Breast Cancer Research Foundation; Susan G. Komen Foundation for the cure, Basser Research Center for BRCA. UPITT/MWH: Hackers for Hope Pittsburgh. VFCTG: Victorian Cancer Agency, Cancer Australia, National Breast Cancer Foundation. WCP: Dr Karlan is funded by the American Cancer Society Early Detection Professorship (SIOP-06-258-01-COUN) and the National Center for Advancing Translational Sciences (NCATS), Grant UL1TR000124.

**Acknowledgements**

All the families and clinicians who contribute to the studies; Sue Healey, in particular taking on the task of mutation classification with the late Olga Sinilnikova; Maggie Angelakos, Judi Maskiell, Gillian Dite, Helen Tsimiklis; members and participants in the New York site of the Breast Cancer Family Registry; members and participants in the Ontario Familial Breast Cancer Registry; Vilius Rudaitis and Laimonas Griškevičius; Drs Janis Eglitis, Anna Krilova and Aivars Stengrevics; Yuan Chun Ding and Linda Steele for their work in participant enrollment and biospecimen and data management; Bent Ejlertsen and Anne-Marie Gerdes for the recruitment and genetic counseling of participants; Alicia Barroso, Rosario Alonso and Guillermo Pita; all the individuals and the researchers who took part in CONSIT TEAM (Consorzio Italiano Tumori Ereditari Alla Mammella), in particular: Cristina Zanzottera, Roberta Villa, Daniela Zaffaroni, Irene Feroce, Mariarosaria Calvello, Riccardo Dolcetti, Laura Ottini, Giuseppe Giannini, Laura Papi, Gabriele Lorenzo Capone, Liliana Varesco, Viviana Gismondi, , Daniela Furlan, Antonella Savarese, Aline Martayan, Stefania Tommasi, Brunella Pilato, and the personnel of the Cogentech Cancer Genetic Test Laboratory, Milan, Italy. Ms. JoEllen Weaver and Dr. Betsy Bove; Marta Santamariña, Ana Blanco, Miguel Aguado, Uxía Esperón and Belinda Rodríguez; IFE - Leipzig Research Centre for Civilization Diseases (Markus Loeffler, Joachim Thiery, Matthias Nüchter, Ronny Baber); We thank all participants, clinicians, family doctors, researchers, and technicians for their contributions and commitment to the DKFZ study and the collaborating groups in Lahore, Pakistan (Muhammad U. Rashid, Noor Muhammad, Sidra Gull, Seerat Bajwa, Faiz Ali Khan, Humaira Naeemi, Saima Faisal, Asif Loya, Mohammed Aasim Yusuf) and Bogota, Colombia (Diana Torres, Ignacio Briceno, Fabian Gil). Genetic Modifiers of Cancer Risk in BRCA1/2 Mutation Carriers (GEMO) study is a study from the National Cancer Genetics Network UNICANCER Genetic Group, France. We wish to pay a tribute to Olga M. Sinilnikova, who with Dominique Stoppa-Lyonnet initiated and coordinated GEMO until she sadly passed away on the 30th June 2014. The team in Lyon (Olga Sinilnikova, Mélanie Léoné, Laure Barjhoux, Carole Verny-Pierre, Sylvie Mazoyer, Francesca Damiola, Valérie Sornin) managed the GEMO samples until the biological resource centre was transferred to Paris in December 2015 (Noura Mebirouk, Fabienne Lesueur, Dominique Stoppa-Lyonnet). We want to thank all the GEMO collaborating groups for their contribution to this study: Coordinating Centre, Service de Génétique, Institut Curie, Paris, France: Muriel Belotti, Ophélie Bertrand, Anne-Marie Birot, Bruno Buecher, Sandrine Caputo, Anaïs Dupré, Emmanuelle Fourme, Marion Gauthier-Villars, Lisa Golmard, Claude Houdayer, Marine Le Mentec, Virginie Moncoutier, Antoine de Pauw, Claire Saule, Dominique Stoppa-Lyonnet, and Inserm U900, Institut Curie, Paris, France: Fabienne Lesueur, Noura Mebirouk.Contributing Centres : Unité Mixte de Génétique Constitutionnelle des Cancers Fréquents, Hospices Civils de Lyon - Centre Léon Bérard, Lyon, France: Nadia Boutry-Kryza, Alain Calender, Sophie Giraud, Mélanie Léone. Institut Gustave Roussy, Villejuif, France: Brigitte Bressac-de-Paillerets, Olivier Caron, Marine Guillaud-Bataille. Centre Jean Perrin, Clermont–Ferrand, France: Yves-Jean Bignon, Nancy Uhrhammer. Centre Léon Bérard, Lyon, France: Valérie Bonadona, Christine Lasset. Centre François Baclesse, Caen, France: Pascaline Berthet, Laurent Castera, Dominique Vaur. Institut Paoli Calmettes, Marseille, France: Violaine Bourdon, Catherine Noguès, Tetsuro Noguchi, Cornel Popovici, Audrey Remenieras, Hagay Sobol. CHU Arnaud-de-Villeneuve, Montpellier, France: Isabelle Coupier, Pascal Pujol. Centre Oscar Lambret, Lille, France: Claude Adenis, Aurélie Dumont, Françoise Révillion. Centre Paul Strauss, Strasbourg, France: Danièle Muller. Institut Bergonié, Bordeaux, France: Emmanuelle Barouk-Simonet, Françoise Bonnet, Virginie Bubien, Michel Longy, Nicolas Sevenet, Institut Claudius Regaud, Toulouse, France: Laurence Gladieff, Rosine Guimbaud, Viviane Feillel, Christine Toulas. CHU Grenoble, France: Hélène Dreyfus, Christine Dominique Leroux, Magalie Peysselon, Rebischung. CHU Dijon, France: Amandine Baurand, Geoffrey Bertolone, Fanny Coron, Laurence Faivre, Caroline Jacquot, Sarab Lizard. CHU St-Etienne, France: Caroline Kientz, Marine Lebrun, Fabienne Prieur. Hôtel Dieu Centre Hospitalier, Chambéry, France: Sandra Fert Ferrer. Centre Antoine Lacassagne, Nice, France: Véronique Mari. CHU Limoges, France: Laurence Vénat-Bouvet. CHU Nantes, France: Stéphane Bézieau, Capucine Delnatte. CHU Bretonneau, Tours and Centre Hospitalier de Bourges France: Isabelle Mortemousque. Groupe Hospitalier Pitié-Salpétrière, Paris, France: Chrystelle Colas, Florence Coulet, Florent Soubrier, Mathilde Warcoin. CHU Vandoeuvre-les-Nancy, France: Myriam Bronner, Johanna Sokolowska. CHU Besançon, France: Marie-Agnès Collonge-Rame, Alexandre Damette. CHU Poitiers, Centre Hospitalier d'Angoulême and Centre Hospitalier de Niort, France: Paul Gesta. Centre Hospitalier de La Rochelle : Hakima Lallaoui. CHU Nîmes Carémeau, France: Jean Chiesa. CHI Poissy, France: Denise Molina-Gomes. CHU Angers, France : Olivier Ingster; Ilse Coene en Brecht Crombez; Ilse Coene and Brecht Crombez; Alicia Tosar and Paula Diaque; Drs .Sofia Khan, Taru A. Muranen, Carl Blomqvist, Irja Erkkilä and Virpi Palola; The Hereditary Breast and Ovarian Cancer Research Group Netherlands (HEBON) consists of the following Collaborating Centers: Coordinating center: Netherlands Cancer Institute, Amsterdam, NL: M.A. Rookus, F.B.L. Hogervorst, F.E. van Leeuwen, S. Verhoef, M.K. Schmidt, N.S. Russell, D.J. Jenner; Erasmus Medical Center, Rotterdam, NL: J.M. Collée, A.M.W. van den Ouweland, M.J. Hooning, C. Seynaeve, C.H.M. van Deurzen, I.M. Obdeijn; Leiden University Medical Center, NL: C.J. van Asperen, J.T. Wijnen, R.A.E.M. Tollenaar, P. Devilee, T.C.T.E.F. van Cronenburg; Radboud University Nijmegen Medical Center, NL: C.M. Kets, A.R. Mensenkamp; University Medical Center Utrecht, NL: M.G.E.M. Ausems, R.B. van der Luijt, C.C. van der Pol; Amsterdam Medical Center, NL: C.M. Aalfs, T.A.M. van Os; VU University Medical Center, Amsterdam, NL: J.J.P. Gille, Q. Waisfisz, H.E.J. Meijers-Heijboer;

University Hospital Maastricht, NL: E.B. Gómez-Garcia, M.J. Blok; University Medical Center Groningen, NL: J.C. Oosterwijk, A.H. van der Hout, M.J. Mourits, G.H. de Bock; The Netherlands Foundation for the detection of hereditary tumours, Leiden, NL: H.F. Vasen; The Netherlands Comprehensive Cancer Organization (IKNL): S. Siesling, J.Verloop; The Dutch Pathology Registry (PALGA): L.I.H. Overbeek; Hong Kong Sanatorium and Hospital; the Hungarian Breast and Ovarian Cancer Study Group members (Aniko Bozsik, Timea Pocza, Zoltan Matrai, Miklos Kasler, Judit Franko, Maria Balogh, Gabriella Domokos, Judit Ferenczi, Department of Molecular Genetics, National Institute of Oncology, Budapest, Hungary) and the clinicians and patients for their contributions to this study; the Oncogenetics Group (VHIO) and the High Risk and Cancer Prevention Unit of the University Hospital Vall d'Hebron, Miguel Servet Progam (CP10/00617), and the Cellex Foundation for providing research facilities and equipment; the ICO Hereditary Cancer Program team led by Dr. Gabriel Capella; the ICO Hereditary Cancer Program team led by Dr. Gabriel Capella; Ana Peixoto, Catarina Santos and Pedro Pinto; members of the Center of Molecular Diagnosis, Oncogenetics Department and Molecular Oncology Research Center of Barretos Cancer Hospital; Heather Thorne, Eveline Niedermayr, all the kConFab research nurses and staff, the heads and staff of the Family Cancer Clinics, and the Clinical Follow Up Study (which has received funding from the NHMRC, the National Breast Cancer Foundation, Cancer Australia, and the National Institute of Health (USA)) for their contributions to this resource, and the many families who contribute to kConFab; the KOBRA Study Group; Csilla Szabo (National Human Genome Research Institute, National Institutes of Health, Bethesda, MD, USA); Eva Machackova (Department of Cancer Epidemiology and Genetics, Masaryk Memorial Cancer Institute and MF MU, Brno, Czech Republic); and Michal Zikan, Petr Pohlreich and Zdenek Kleibl (Oncogynecologic Center and Department of Biochemistry and Experimental Oncology, First Faculty of Medicine, Charles University, Prague, Czech Republic); Anne Lincoln, Lauren Jacobs; the participants in Hereditary Breast/Ovarian Cancer Study and Breast Imaging Study for their selfless contributions to our research; the NICCC National Familial Cancer Consultation Service team led by Sara Dishon, the lab team led by Dr. Flavio Lejbkowicz, and the research field operations team led by Dr. Mila Pinchev; the investigators of the Australia New Zealand NRG Oncology group; members and participants in the Ontario Cancer Genetics Network; Leigha Senter, Kevin Sweet, Caroline Craven, Julia Cooper, Amber Aielts, and Michelle O'Conor; Yip Cheng Har, Nur Aishah Mohd Taib, Phuah Sze Yee, Norhashimah Hassan and all the research nurses, research assistants and doctors involved in the MyBrCa Study for assistance in patient recruitment, data collection and sample preparation, Philip Iau, Sng Jen-Hwei and Sharifah Nor Akmal for contributing samples from the Singapore Breast Cancer Study and the HUKM-HKL Study respectively; the Meirav Comprehensive breast cancer center team at the Sheba Medical Center; Christina Selkirk, Helena Jernström, Karin Henriksson, Katja Harbst, Maria Soller, Ulf Kristoffersson; from Gothenburg Sahlgrenska University Hospital: Anna Öfverholm, Margareta Nordling, Per Karlsson, Zakaria Einbeigi; from Stockholm and Karolinska University Hospital: Anna von Wachenfeldt, Annelie Liljegren, Brita Arver, Gisela Barbany Bustinza; from Umeå University Hospital: Beatrice Melin, Christina Edwinsdotter Ardnor, Monica Emanuelsson; from Uppsala University: Hans Ehrencrona, Maritta Hellström Pigg, Richard Rosenquist; from Linköping University Hospital: Marie Stenmark-Askmalm, Sigrun Liedgren; Cecilia Zvocec, Qun Niu; Joyce Seldon and Lorna Kwan; Dr. Robert Nussbaum, Beth Crawford, Kate Loranger, Julie Mak, Nicola Stewart, Robin Lee, Amie Blanco and Peggy Conrad and Salina Chan; Simon Gayther, Carole Pye, Patricia

Harrington and Eva Wozniak; Geoffrey Lindeman, Marion Harris, Martin Delatycki, Sarah Sawyer, Rebecca Driessen, and Ella Thompson for performing all DNA amplification.
